# The Arabidopsis *HOP2* Gene Has a Role in Preventing illegitimate Exchanges between Nonhomologous Chromosomes

**DOI:** 10.1101/2021.08.04.455073

**Authors:** Yisell Farahani-Tafreshi, Chun Wei, Peilu Gan, Jenya Daradur, C. Daniel Riggs, Clare A. Hasenkampf

## Abstract

Meiotic homologous chromosomes pair up and undergo crossing over. In many eukaryotes both intimate pairing and crossing over require the induction of double stranded breaks (DSBs) and subsequent repair via Homologous Recombination (HR). In these organisms, two key proteins are the recombinases RAD51 and DMC1. Recombinase-modulators HOP2 and MND1 have been identified as proteins that assist RAD51 and DMC1 and are needed to promote stabilized pairing. We have probed the nature of the genetic lesions seen in *hop2* mutants and looked at the role of HOP2 in the fidelity of genetic exchanges. Using γH2Ax as a marker for unrepaired DSBs we found that *hop2-1* and *mnd1* mutants have different appearance/disappearance for DSBs than wild type, but all DSBs are repaired by mid-late pachytene. Therefore, the bridges and fragments seen from metaphase I onward are due to mis-repaired DSBs, not unrepaired ones. Studying Arabidopsis haploid meiocytes we found that wild type haploids produced the expected five univalents, but *hop2-1* haploids suffered many illegitimate exchanges that were stable enough to produce bridged chromosomes during segregation. Our results suggest that HOP2 has a significant active role in preventing nonhomologous associations. We also found evidence that HOP2 plays a role in preventing illegitimate exchanges during repair of radiation-induced DSBs in rapidly dividing petal cells. Thus, HOP2 plays both a positive role in promoting homologous chromosome synapsis and a separable role in preventing nonhomologous chromosome exchanges. Possible mechanisms for this second important role are discussed.

**SIGNIFICANCE STATEMENT:** The fidelity of Homologous Recombination (HR) during meiosis is essential to the production of viable gametes and for maintaining genome integrity in vegetative cells. HOP2 is an important protein for accurate meiotic HR in plants. We find high levels of illegitimate repairs between nonhomologous chromosomes during meiosis and in irradiated petal cells in *HOP2* mutants. We consider mechanisms of how this second role might be accomplished.

## INTRODUCTION

Meiosis is a remarkably complex process wherein the action of proteins governing chromosome movement have been mechanistically linked with proteins that participate in repairing DNA damage through Homologous Recombination (HR). Meiosis provides for new allelic combinations in the resulting nuclei; it occurs in conjunction with sexual reproduction and, through its two divisions of the chromosomes, reduces the ploidy level in anticipation of fertilization. In most eukaryotes the regular segregation at the first meiotic division is accomplished by having homologous chromosomes pair intimately, form synaptonemal complex along their combined pairing axes, and have one sister chromatid from each of the two homologs undergo at least one, precise, reciprocal genetic exchange (*aka* crossover). The occurrence of crossovers combined with sister chromatid cohesion creates physical ties between homologs that allow them to coordinately orient on the metaphase I plate (Sansam and Pezza 2015). At anaphase I the cohesins are removed along non-centromeric portions of the chromosome arms allowing homologs to segregate to opposite poles. Thus, crossovers (CO) are required for high fidelity homolog segregation and are also a means of creating new allelic combinations of genes on homologs. (Reviewed in Mercier *et al.,* 2015).

Eukaryotes with a standard meiosis absolutely rely on HR (Keeney *et al.* 2014). Meiotic HR proceeds as summarized in reviews by Ranjha *et al.* (2018) and Crickard and Greene (2018). Early in prophase I double strand breaks (DSBs) are intentionally induced into the DNA by SPO11, which remains attached to the 5’ end of the break. This type of break attracts the MRN/MRX complex, which sets in motion 1) the removal of a small section of ssDNA bound to SPO11, 2) activation of signalling molecules ATM/ATR, 3) further re-sectioning of the 5’ end to create a longer segment of ssDNA and 4) successive addition of proteins that ultimately lead to the binding of RecA type recombinases RAD51 and in many organisms DMC1. RAD51/DMC1 bind to create a nucleoprotein filament that can then invade dsDNA and create a D-loop intermediate. The resulting displaced ssDNA can then serve as a template to replace the genetic information lost during the removal of SPO11 and subsequent re-sectioning. Once this repair replication has occurred, there are several different ways the D-loop can be processed to complete the repair, yet only some of these pathways can generate a CO. Another important aspect of DSB repair is the phosphorylation of histone H2AX by ATM/ATR in a wide region flanking a DSB; this γH2AX is important in the recruitment of key proteins to the damage site (Lowndes and Toh 2006; Amiard *et al.*, 2010, Yao et al., 2020).

During meiosis at least one of the D-loops that formed between the DNA of homologs must be resolved in a manner that produces a CO. For this CO to create two genetically balanced DNA helices, the homologous chromosomes must be in precise register. Many eukaryotes (*eg*. *S. cerevisae*, mammals, higher plants) rely on the HR system to assist in this precise stabilized alignment of the homologs (Keeney *et al.,* 2014), but other eukaryotes (*eg. C. elegans* and *D. melanogaster*) have HR-independent mechanisms to promote intimate pairing of homologs and these organisms only rely on HR to generate the COs themselves (Jang *et al.*, 2003). As reviewed in Keeney *et al.* (2014), organisms with HR-independent pairing mechanisms induce a modest number of DSBs. In contrast, eukaryotes that rely on HR for both the intimate pairing of homologous chromosomes and for the generation of COs have approximately ten times as many DSBs per CO produced. Induction of DSBs is a dangerous undertaking, and repairing these breaks in a manner that generates COs introduces additional risk. Even small misalignments of homologous regions can generate duplications and deletions. COs occurring between nonhomologous chromosomes can create translocations which can produce acentric, dicentric or even polycentric chromosomes that cannot segregate properly at anaphase. Because of the hazards of incorrect repair of DSBs, it is not surprising that meiotic repair of DSBs is carefully regulated, especially for CO-outcomes. Nor is it surprising that organisms that use HR for stabilizing intimate pairing, with their increased number of DSBs, appear to require additional proteins to accomplish this pairing and coordinate it with CO production.

Three proteins that are only found in eukaryotes using HR for stabilization of pairing are DMC1, HOP2 and MND1. DMC1 is a RecA-like protein with homology to RAD51 (Bishop *et al.*, 1992). DMC1 probes homology through D-loop formation and triggers structural changes that can lead to stabilized pairing followed by SC formation; DMC1 also is essential for CO formation (Bishop *et al.*, 1992, Pittman *et al.*, 1998, Couteau *et al.*, 1999). DMC1 and RAD51 arose from a gene duplication and have very similar biochemical properties; both can form nucleofilaments on resected ssDNA and form D-loops (Crickard and Greene 2018). Immunolocalization studies have revealed that DMC1 and RAD51 do not always colocalize; individual DMC1 and RAD51 foci are regularly seen, but so are doublets of DMC1 with RAD51 [yeast (Shinohara *et al.*, 2000) and Arabidopsis (Kurzbauer *et al.*, 2012)]. Occurrence of these doublets has led to the notion that RAD51 and DMC1 might occur on opposite ssDNA overhangs of a DSB. More recent studies in yeast [Super resolution/dSTORM microscopy (Brown *et al.*, 2015) and reconstitution studies (Crickard *et al.,* 2019)] suggest that DMC1 and RAD51 may form mixed recombinase nucleofilaments on each of the ssDNA overhangs of a DSB. The precise roles of DMC1 and RAD51 in the production of COs is yet to be determined, but in organisms with both proteins they are both necessary to achieve normal levels and distributions of COs (Su *et al.*, 2017), and their proximity to each other suggests their combined activities must be carefully coordinated.

HOP2 and MND1 are not RecA family members, but HOP2, in combination with MND1, is required to achieve normal levels of stabilized pairing, followed by SC formation and COs between homologous chromosomes (Leu *et al.*, 1998, Petukhova *et al.,* 2003, Kerzendorfer *et al.* 2006, Vignard *et al.*, 2007, Stronghill *et al.*,2010, Shi *et al*, 2019). Using Super resolution (SIM) microscopy with rice meiotic cells, Shi and coworkers (2019) showed that HOP2 occurs in the looped domains, along the pairing axis and even within the central region. *In vitro* biochemical analyses suggest that HOP2-MND1 is a modulator of DMC1 and/or RAD51 activity (Petukhova *et al.*, 2005, Neale and Keeney 2006, Ploquin *et al.* (2007). Analyses of the structure of HOP2-MND1 heterodimers suggest they form a complex that can bind both ssDNA and dsDNA, and allow tethering of dsDNA to resected sections of ssDNA that have been coated with DMC1 and RAD51 (Zhao *et al.*, 2014, Moktan *et al.*, 2014, Kang *et al.*, 2015).

Mutants of *hop2* have been studied extensively in yeast and mammals. These mutants do not complete meiosis. In yeast SC formation is delayed and DMC1 foci start to form normally but accumulate to higher-than-normal levels and are still present at the time of prophase I arrest. DSBs appeared to be unrepaired in the arrested cells, and nonhomologously formed SC was observed (Leu *et al.*, 1998). Tsubouchi and Roeder (2003), using a FISH pairing assay found evidence of nonhomologous pairing in *hop2* and at higher levels than observed in *dmc1* or *rad51* mutants. In mice *hop2* mutants have reduced levels of SC formation and much of what does form appears to be nonhomologous in nature. The meiocytes arrest in pachytene, with some unrepaired DSBs and persistent DMC1 and RAD51 foci (Petukhova *et al.* 2003). Thus, in *hop2* mutants in mice and yeast, the mutants arrest in prophase I with incomplete repair of DSBs, greatly reduced levels of homologous SC, and evidence of some nonhomologous chromosome associations. The model plant Arabidopsis provides an interesting perspective on the interdependence of pairing stabilization and HR-based DSB repair because many meiotic mutants do not experience meiotic prophase I arrest and thus complete meiosis.

Previously we examined *hop2-1* diploid mutants and found very little SC formation in zygotene or pachytene (Pathan *et al.* 2013), despite a prolonged zygotene stage. We also observed complex chromosome interconnections at metaphase I and chromatin bridges at both the first and second meiotic anaphase (Stronghill *et al.*, 2010). The entangled chromosomes under tension at metaphase I and the dicentric chromosomes that produce chromatin bridges are indicative of DSBs that are <u>mis-</u>repaired during prophase I. In this paper we have further explored mis-repair in haploid *hop2-1* mutants, determined the durations of leptotene and zygotene in diploid *hop2-1* and *mnd1* plants, and determined the timing of appearance/disappearance of DSBs as monitored by γH2AX foci. We also investigated the *hop2-1* phenotype in the rapidly dividing diploid cells of flower petals.

## RESULTS

### Duration of Leptotene and Zygotene in hop2-1 and mnd1 Mutants

We determined the duration of leptotene and zygotene stages for L*er*, *hop2-1*, Columbia and *mnd1* plants by incubating inflorescences in media with EdU for two hours, transferring inflorescences to media without EdU, then collecting inflorescences at two-hour intervals with subsequent fixation. Here we used a preservation protocol that retains the microsporocytes as a filament (sac) of meiotic cells surrounded by a single layer of cells collectively known as the tapetum. Each filament, containing ~30 meiotic cells, was processed for both detection of EdU incorporation and immunolocalization of histone γH2AX. Table S1 contains the complete data set for all 2086 filaments. This table includes information on meiotic and tapetal nuclei characteristics, EdU incorporation in meiotic and tapetal cells, and immunostaining for histone γH2AX. Negative controls yielded no EdU signals and no histone γH2AX staining.

Using the time point of the first appearance of EdU incorporation in meiotic nuclei, we determined the duration of the leptotene and zygotene stages for L*er*, *hop2-1*, Col and *mnd1* genotypes. Our estimates of the leptotene and zygotene durations are provided in Table 1. The values for L*er* control plants agree with our previous work (Stronghill *et al.*, 2014) and those for Col control plants agree with those reported by Armstrong (2013). Our data for *hop2-1* plants are suggestive of it having a shorter leptotene (*hop2-1* is 3-5 hrs; L*er* is 5-7 hr), but since the estimates of leptotene for L*er* and *hop2-1* plants overlap (because we collected samples every two hrs) we cannot definitively conclude that leptotene is shorter in duration. Relative to zygotene, our data does clearly show that *hop2-1* has a prolonged zygotene. In *mnd1* plants leptotene appears to be the same duration as the Col control. The *mnd1* mutant, like *hop2-1* has a prolonged zygotene.

**Table 1.** Durations of Early Meiotic Prophase I Substages

| Genotype | Leptotene | Zygotene |
| --- | --- | --- |
| Landsberg *erecta* | 5-7 hours | 5-7 hours |
| *hop2-1* | 3-5 hours | 9-11 hours |
| Columbia | 5-7 hours | 7-9 hours |
| *mnd1* | 5-7 hours | 11-13 hours |

Filaments of meiotic cells were collected every two hours after removal of EdU from the media. Buds were preserved and meiotic stage(s) found were recorded as was the presence of EdU labeling. The duration of each meiotic stage was calculated based on the time point of first appearance of label in that stage and in the next stage, subtracting the time of appearance in the later from that of the former.

### Assessment of meiotic DSBs using the appearance and disappearance of γH2AX foci

Next, we collected γH2AX foci data for each meiotic stage for those nuclei that had incorporated EdU; staging and time point details are found in the Experimental Procedures. Three categories of small γH2AX foci were recorded: ‘no small foci’, ‘1-50 small foci’, and ‘51-150 foci’. In addition to small discrete foci of these three categories, we also saw a few, larger, brighter staining regions in the wild type L*er* and Col controls in early pachytene; these few bright foci only appeared after the small foci had disappeared. Representative micrographs of meiotic cells with their extent of γH2AX immunostaining are provided in Fig. 1 for L*er* and *hop2-1* plants and in Fig. 2 for Col and *mnd1* plants. Figure 3 summarizes the levels and extent of histone γH2AX formation seen at each prophase I substage in each of the four genotypes. Our profiles of L*er* and Col were identical, with small foci first appearing in leptotene and peaking in zygotene. By early pachytene, the L*er* and Col controls had no small foci, but had 2-5 brighter staining regions; examples of these special foci can be seen in Fig. 2 (eP in L*er*) and Fig. 3 (eP in Col). By mid-to-late pachytene, L*er* and Col meiotic cells had no histone γH2AX staining. Our Col results agree fairly well with those of Sanchez Moran *et al.* (2007), taking into account the differences in chromosome preparations and in sampling times (we pulled samples every two hours from the addition of EdU; they pulled samples at 0,3,5,12,18,24 and 30 hrs). They reported first seeing γH2AX foci in the G2-early leptotene interval; we first saw foci in leptotene; in our study and theirs γH2AX foci had disappeared by the end of prophase I. Our observations of the few brighter γH2AX foci seen in the wild type controls, also corresponds to the pattern reported for HEI10, a protein required for the formation of class I crossovers.

**Figure 1.**
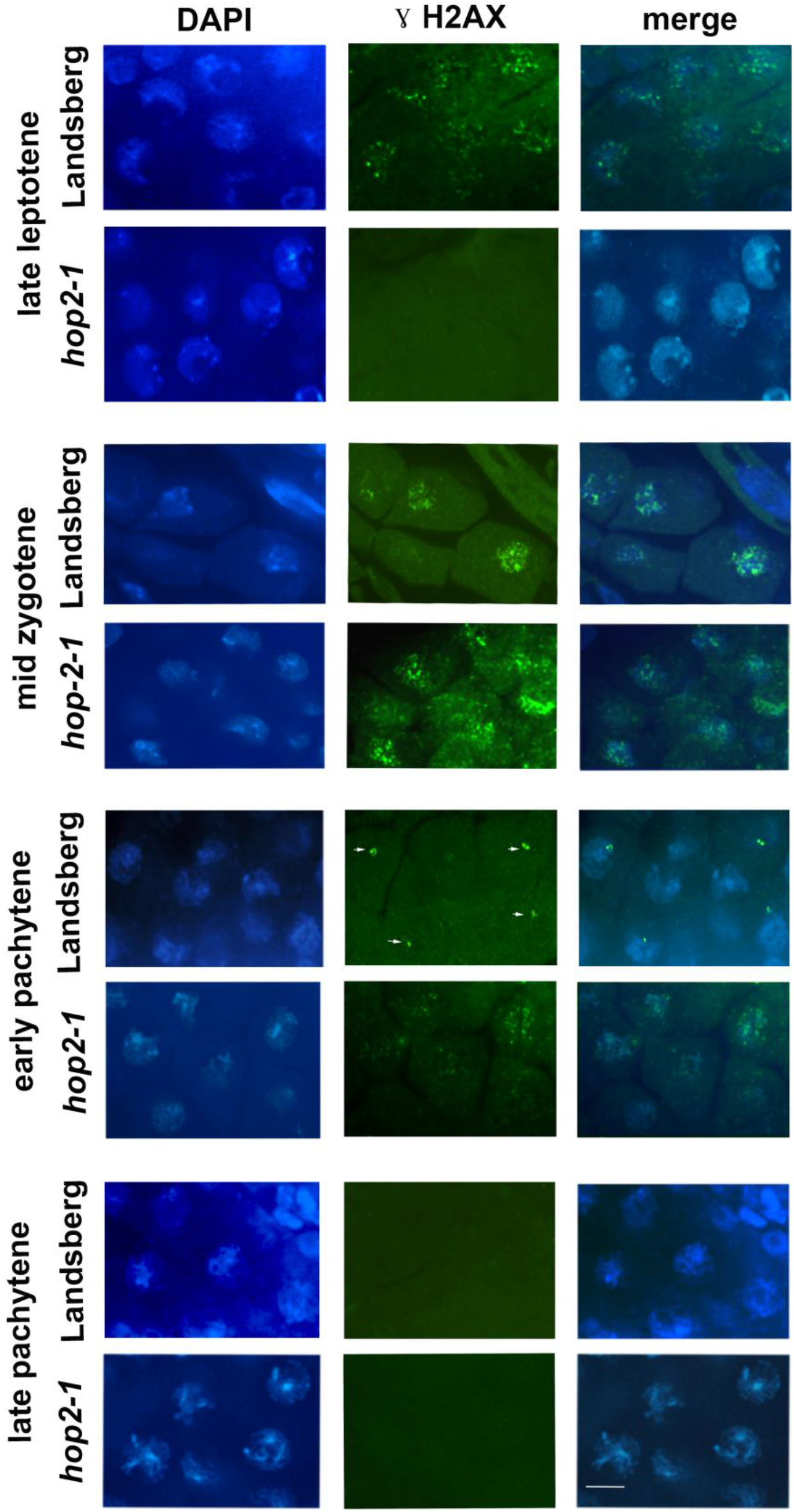
γH2AX staining in prophase I substages in *Arabidopsis thaliana* Landsberg *erecta* (L*er*) and *hop2-1* diploids. Panel 1 provides light micrographs of DAPI-labeled chromatin. Panel 2 shows the same nuclei immunostained for γH2AX, detected with Alexa 488 (green fluorescence). Panel 2’s arrows point to the larger bright staining regions that are only seen in the L*er* (and Col) wild type early pachytene nuclei. Panel 3 micrographs are merged images for each filament region. Scale bar is 5μm.

**Figure 2.**
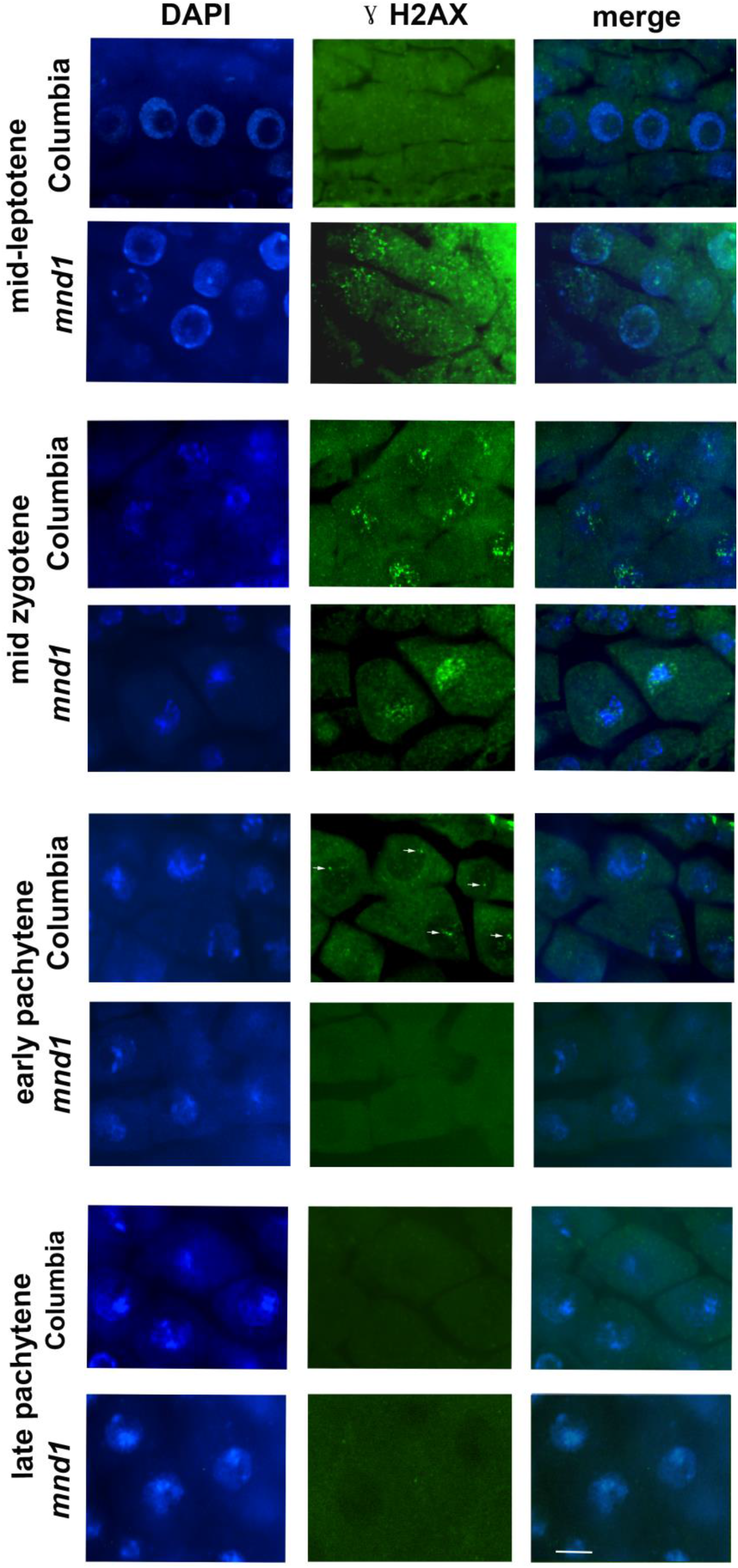
γH2AX staining in prophase I substages in Columbia (Col) and *mnd1* diploids. Panel 1 provides light micrographs of DAPI-labeled chromatin. Panel 2 has the same nuclei immune-stained for γH2AX, detected with Alexa 488 (green fluorescence). Panel 2 arrows point to the large bright staining regions that are only seen in the Col (and L*er*) wild type early pachytene nuclei. Panel 3 micrographs are merged images for each filament region. Scale bar is 5μm.

**Figure 3.**
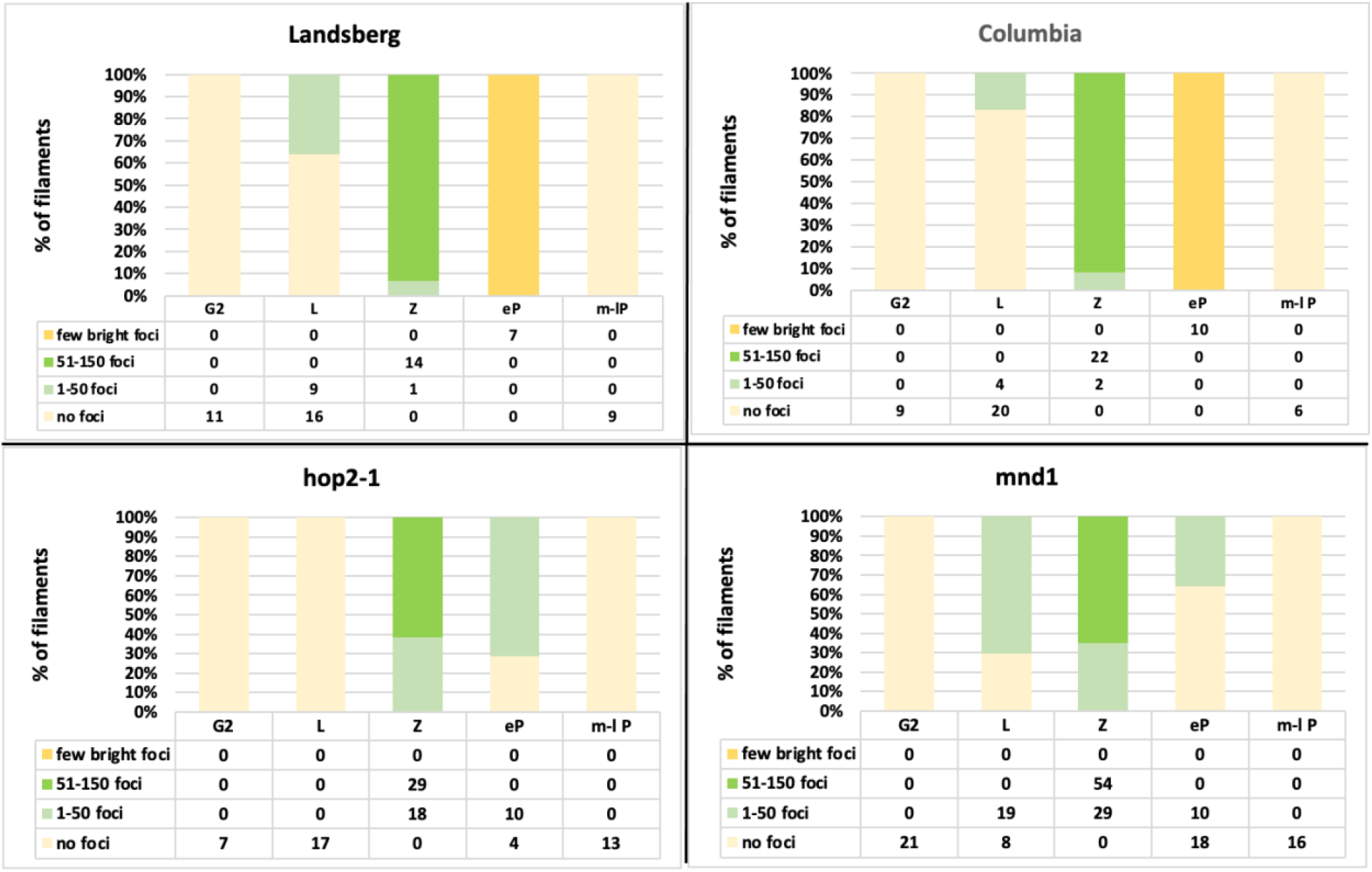
Timing of γH2AX foci in genotypes Landsberg *erecta* (L*er*) and *hop2-1* (L*er* background) and Columbia (Col) and *mnd1* (Columbia background), from samples that incorporated EdU. The duration for each meiotic stage for each genotype was determined based on the timepoint at which EdU staining was first observed for each meiotic stage. Then all filaments of meiotic cells that had EdU incorporated for those stages, in that time window, were assessed for histone γH2Ax levels. Each filament contains ~30 meiotic cells, surrounded by the tapetum. Each filament was classified as having ‘no foci’, ‘1-50 small foci’, ‘51-150 small foci’, or ‘a few brighter larger foci’. The number of filaments within each category are given in tabular form below each genotype’s percentage of nuclei in each category. Stages assessed were G2 preceding meiosis, leptotene (L), zygotene (Z), early pachytene (eP) and mid-to late pachytene (m-lP). The mutant *hop2-1* had significantly fewer foci than L*er* at both leptotene and zygotene and the differences were statistically significant (p<0.003 and p<0.00001 respectively). In contrast the *mnd1* mutant had significantly more foci visible than Col at leptotene (p<0.00001), but significantly fewer than Col by zygotene (p<0.0001).

Hurel *et al.* (2018) observed that as synapsis progressed from zygotene to pachytene HEI10 staining was reduced to only a few bright foci.

Compared to its wild type control, the *hop2-1* mutant lags a little behind L*er* in the appearance of small γH2AX foci at leptotene and zygotene; the differences were statistically significant (p< 0.003 and p<0.00001 respectively). In contrast, the *mnd1* mutant had a significantly higher foci level than its Col control at leptotene (p< 0.00001), but fewer at zygotene (p< 0.00001). Both mutants still have some small foci at early pachytene. Neither *hop2-1* nor *mnd1* ever had any of the larger, brighter γH2AX foci that were observed in L*er* and Col in early pachytene. Importantly, we observed that all histone γH2AX foci disappeared in all genotypes by mid-late pachytene.

We also looked at γH2AX appearance and disappearance in our complete data collection, including all meiotic cells, whether or not they had incorporated EdU. This is a much larger sample; the results are shown in Fig.S1. This larger sample showed the same temporal patterns that we saw in the samples that had incorporated EdU, and again, all γH2AX foci had disappeared by mid-late pachytene in all genotypes. The repair of all DSBs by mid-late pachytene in these diploid mutants is consistent with the fact that no fragments were reported in *hop2-1* (Stronghill *et al.* 2010) or *mnd1* (Vignard *et al.*, 2007) plants in prophase I or metaphase I. Thus our γH2AX results confirm that the extremely entangled chromosomes seen in late prophase I and metaphase I in *hop2-1* and *mnd1* are due to incorrectly repaired DSBs, <u>not</u> unrepaired ones. The chromosome fragments and bridges seen in the later stages in both mutants are generated as di- or polycentric chromosomes attempt to move to opposite poles during segregation (Domenichi *et al.*, 2006, Panoli *et al.*, 2006, Kerzendorfer *et al.*, 2006; Vignard *et al.*, 2007, Stronghill *et al.* 2010).

### Meiosis in haploids of Ler and hop2-1

To further explore the role of *HOP2* in preventing incorrect repairs with illegitimate chromosome segments, and to separate it from the events of synapsis, we examined meiosis in spread chromosome preparations of L*er* and *hop2-1* haploids that have no opportunity to engage with a homolog. Cifuentes *et al.*, (2013) examined meiosis in wild type haploids and reported that the haploids formed and repaired DSBs without forming crossovers or illegitimate chromosome exchanges. Only intact univalents were observed in metaphase I and these segregated randomly to the poles. Thus, wild type haploids have regulatory mechanisms in place to prevent DSB HR repair from using nonhomologous chromosome regions, even in the absence of synapsis. As such, Arabidopsis haploids provide an excellent opportunity to tease apart the established role of HOP2 in promoting stabilized homologous chromosome associations from our proposed role for HOP2 in actively preventing nonhomologous chromosome exchanges. We reasoned that if HOP2 was important in actively preventing nonhomologous chromosome regions from being joined together during repair, then nonhomologous exchanges in haploid *hop2-1* plants would be much more extensive than in wild type haploids. Indeed, this is what we found as described below.

For our analysis we examined five plants for L*er* control haploids and five *hop2-1* haploids. For each haploid plant used in the analysis, the mitotic karyotype was determined (Fig.S2a and b), and we confirmed the presence of each different L*er* chromosome using chromosome-specific SSLPs (Fig.S2c). Once we were certain that we had identified genuine haploids, we genotyped the plants to distinguish *HOP2* from *hop2-1* haploids (Fig.S2c, two bottom lanes). For the confirmed haploids we collected and preserved flower buds and made meiotic, spread-style chromosome preparations. We focused on nuclei ranging from metaphase I through telophase II. We analyzed 427 meiotic nuclei for the L*er (HOP2*) haploids and 292 nuclei from *hop2-1* haploids.

Our results for the L*er* plants (Fig.4 a, c, e, g) were very similar to those observed for Columbia haploids (Cifuentes *et al.*, 2013). The majority (84%) of first division segregations resulted in five univalents segregating randomly (Figure 4c). Premature sister chromatid separation was observed in 16% of anaphase I nuclei (Figure 4e), a phenomenon also observed by Cifuentes *et al.*, (2013). In both the first and second meiotic divisions only 3% of nuclei had illegitimate connections. Thus, the haploid control nuclei had some problems with premature sister chromatid separation but exhibited only a low level of nonhomologous associations.

**Figure 4.**
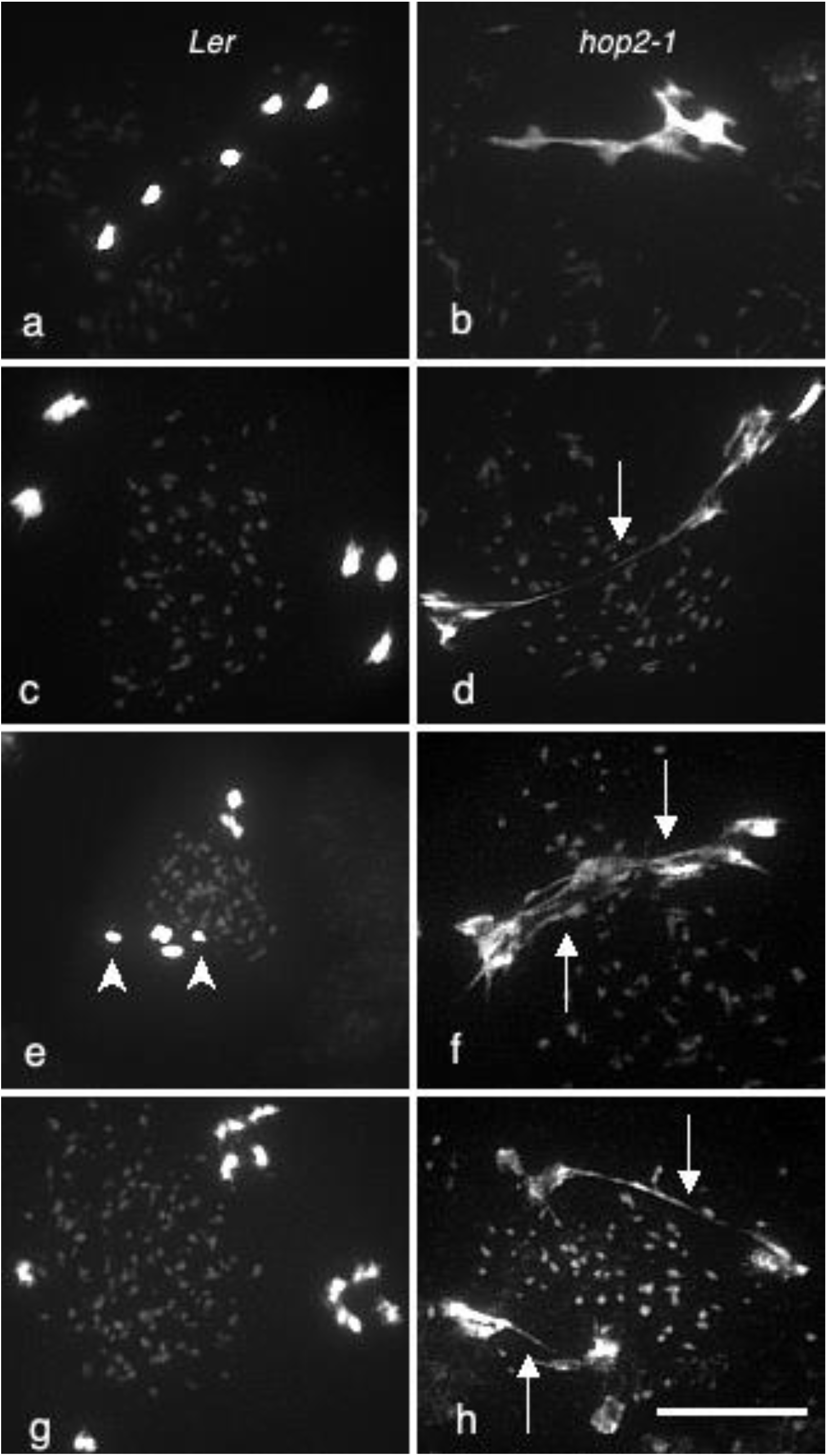
L*er* and *hop2-1* Haploid Meiotic Divisions. Panels 4 a,c,e and g are of L*er* haploids; panels b,d,f and h show *hop2-1* mutants. Panels a and b show Metaphase I; c,d,e and f are of anaphase I nuclei and g and h are from second division nuclei. The arrowheads in panel e point to sister chromatids that prematurely segregated. The long arrows of panels d, f, and h point to bridges caused by illegitimate chromosome connections in *hop2-1*. Scale bar= 5μm

Chromosome segregation behavior in *hop2-1* haploids was strikingly different. Examples of the aberrations are shown in Fig.4 b, d, f and h. The meiotic nuclei of *hop2-1* haploids regularly demonstrated nonhomologous associations that resisted resolution during segregation. We found that 99% (248/251) of first division nuclei had multi-chromosome associations at metaphase I (Figure 4b) or chromatin bridges at anaphase I or telophase I (Fig. 4 d and f). Of the 41 different second division nuclei we observed, all showed evidence of chromatin bridges (e.g. Fig 4h).

From our analysis of haploid meiosis, we conclude that a functional *HOP2* gene is required to prevent nonhomologous chromosome regions from being used to repair DSBs during prophase I of meiosis. Comparing our results with *hop2-1* haploids to published work on diploid *hop2* and *mnd1* plants, it is clear that both diploid and haploid Arabidopsis meiotic nuclei require HOP2 to prevent large scale DSB repairs that connect nonhomologous regions together as part of the repair process. We did not look at γH2AX appearance/disappearance in our haploid plants but we did look specifically for any evidence of fragments in the metaphase I nuclei from our haploid *hop2-1* plants. In the 122 metaphase I nuclei examined, only 3 nuclei had a possible chromosome fragment before anaphase I. Thus, in the vast majority of haploid *hop2-1* nuclei, all DSBs are repaired by metaphase I, with fragments only being generated after opposing anaphase I segregation forces begin to break di- or multicentric chromosomes.

### hop2-1 plants have increased numbers of incorrectly repaired DSBs outside of meiosis, after γ -irradiation

While *hop2-1* plants in Arabidopsis are sterile, no obvious vegetative phenotype has been reported in previous studies when plants are grown under normal growth chamber conditions (Schommer *et al.* 2003; Stronghill *et al.*, 2010). However, *HOP2* expression has been reported in vegetative tissues that have high mitotic indices (Klepikova *et al.*, 2016). In light of our findings that during meiosis, *hop2-1* exhibits illegitimate exchanges, we decided to investigate whether petal mitotic nuclei of *hop2-1* plants had more illegitimate chromatin exchanges than did the L*er* control. We looked at unirradiated plants and plants exposed to 25 Gy of γ -irradiation with 70 min or 24 hr recovery times. Petals taken from stage 9/10 flowers (Smyth *et al.*, 1990) are easy to collect and have a fairly high mitotic index (Fig. S3). We examined ~15,000 cells from each petal, determining the mitotic index and recording the presence or absence of chromatin bridges in all observed anaphase and telophase cells. Examples of bridges are shown in Figure 5, and the results are summarized in Table 2. In the unirradiated plants the petal mitotic index was 2.4 and 2.3%, respectively for L*er* and *hop2-1,* and no chromatin bridges were observed for either. This result is consistent with the lack of vegetative phenotype observed for growth chamber grown plants of *hop2-1*.

**Table 2.** Mitotic Index and Extent of Bridges for Irradiated Petals

| Treatment | # Mitoses |  | Mitotic Index (%) |  | Bridged Mitosis (%) |  |
| --- | --- | --- | --- | --- | --- | --- |
|  | *Ler* | *hop2-1* | *Ler* | *hop2-1* | *Ler* | *hop2-1* |
| No irradiation | 300 | 300 | 2.4 | 2.3<sup>†</sup> | 0.0 | 0.0 |
| Irradiation + 70 min | 86 | 85<sup>†</sup> | 0.30 | 0.27<sup>†</sup> | 30.2 | 49.4<sup>‡</sup> |
| Irradiation + 24 hours | 278 | 316<sup>†</sup> | 1.0 | 1.3<sup>†</sup> | 6.8 | 12.0<sup>‡</sup> |

The first 15,000 cells observed for each petal had the mitotic index determined as a percentage of cells with condensed chromosomes.
<sup>†</sup> The observed small differences in mitotic indices between *Ler* with *hop2-1* were not statistically significant.
<sup>‡</sup> The observed differences in % mitoses with chromatin bridges when comparing *Ler* to *hop2-1* were statistically significant for both of the irradiation treatments (p-value = 0.01513 and 0.04511, respectively, for the 70 min and 24 hr recovery times).

**Figure 5.**
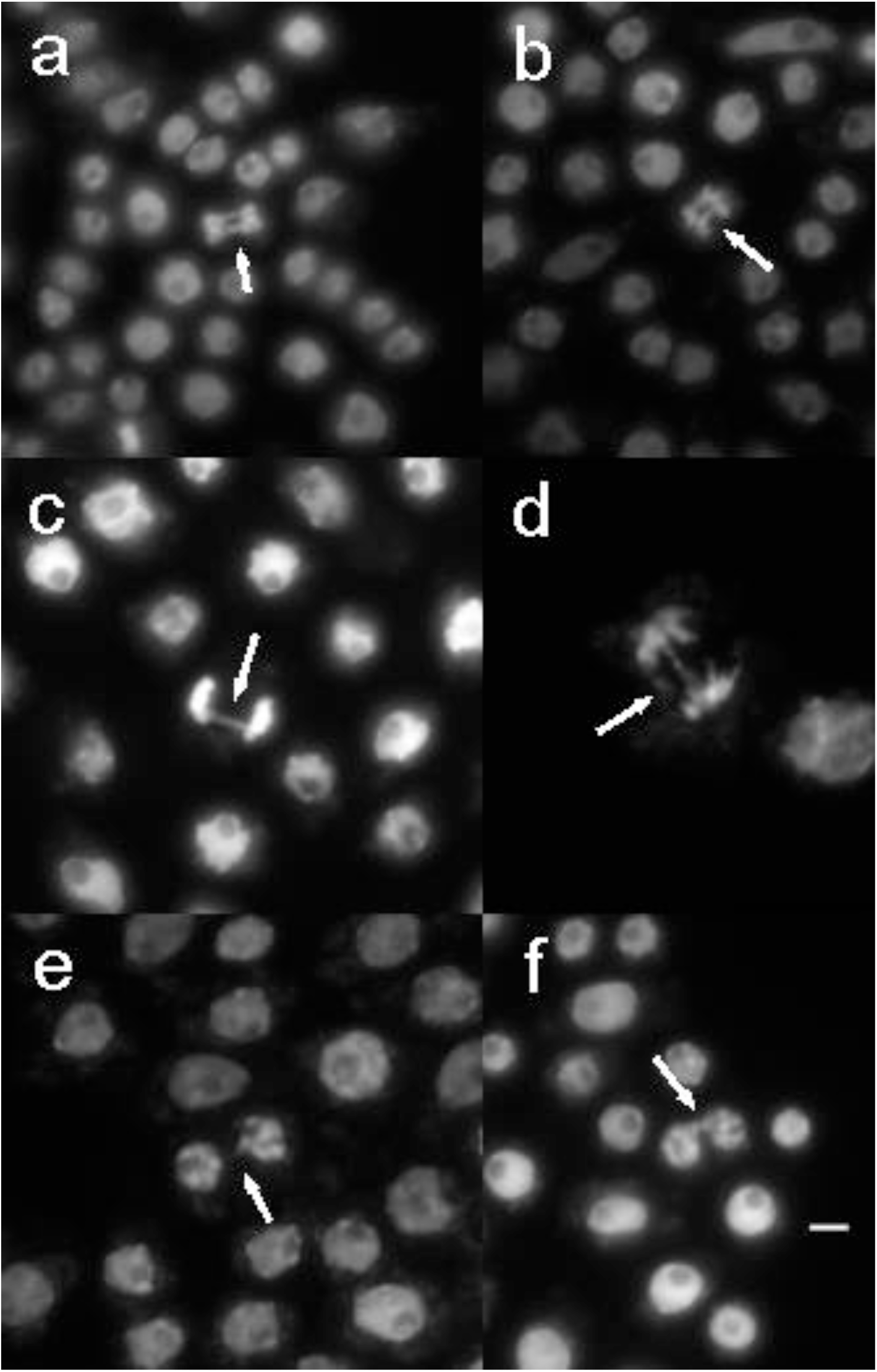
Abnormal chromosome segregations were found in both L*er* and *hop2-1* plants, 70mins after irradiation. Panels a,c and e are from L*er* plants; b,d and f are from *hop2-1* plants. Arrows point to bridges seen in anaphase and telophases of a-d. Panels e and f show post-mitotic cells that retain chromatin connections (arrows) between the two products of an aberrant mitosis. Scale bar is 10 μm.

In our first irradiation experiment the plants were irradiated and then flower buds were processed 70 min after irradiation. While cells at all stages of the cell cycle were affected by the radiation, only cells already in late G2 or mitosis would have been able to produce mitotic figures only 70 min after irradiation. Both L*er* and *hop2-1* exhibited a marked reduction in the mitotic index (to 0.3 and 0.27%, respectively). This likely is due to the inability of most cells to repair the breaks and transit the G2 checkpoint in just 70 min; most of the mitotic cells we observed likely were already past the G2 checkpoint when irradiated. We observed chromatin bridges in both L*er* and *hop2-1* samples (Fig. 5), but bridges were more frequently seen in *hop2-1* than in L*er* (49.4 versus 30%). This difference was statistically significant (p-value = 0.01513).

We also looked at irradiated plants that were allowed to recover for 24 hours after irradiation. By allowing 24 hrs for repair and for progression through the cell cycle, cells damaged by the radiation at all stages of the cell cycle could have had time to enter mitosis. For these plants the mitotic index rebounded to 1.0% and 1.3% in L*er* and *hop2-1* plants, respectively. Not only was the mitotic index closer to normal, but the frequency of chromatin bridging was reduced to 6.8% in L*er* and 12.0% in *hop2-1*. Even though the frequency of bridges was reduced overall, the *hop2-1* plants still had a statistically significant higher frequency of abnormal repairs than did the L*er* control (p-value of 0.04511). Thus, the *hop2-1* plants appear to be able to repair DSBs at a similar rate to wild type (because both L*er* and *hop2-1* plants had similar mitotic indices in both irradiation experiments) but *hop2-1* plants allow more repairs that generate illegitimate exchanges. Therefore, even in petal nuclei, HOP2 appears to have a role in reducing the likelihood of conducting illegitimate repair with a nonhomologous chromosome region.

## Discussion

In yeast (Leu *et al.* 1998), mammals (Petukhova *et al.* 2003), Arabidopsis (Stronghill *et al.* 2010) and rice (Shi *et al.* 2019), HOP2 appears to have a direct or indirect role in stabilizing chromosome pairing for effective chromosome synapsis (SC formation). In yeast (Leu *et al.* 1998) and mammals (Petukhova *et al.* 2003) *hop2* mutants arrest in prophase I with unrepaired DSBs. In Arabidopsis, and plants in general, there is no prophase I arrest so meiosis can be followed to completion. Thus, even though *hop2-1* (Stronghill *et al.* 2010) and *mnd1*(Vignard *et al.* 2007) mutants have a prolonged zygotene, and no appreciable amounts of synaptonemal complex, they proceed through meiosis. Metaphase I nuclei in *hop2-1* and *mnd1* have extensively interconnected chromosomes, but no distinct fragments. But chromosome entanglements persist into anaphase I and anaphase II where chromatin bridges with consequent chromosome fragmentation are observed. In our work here we have demonstrated that all DSBs that are generated during prophase I are repaired prior to the completion of prophase I in both *hop2-1* and *mnd1* diploids. Additionally, Uanschou *et al.* (2013) reported that DMC1 and RAD51 foci appear and disappear normally in *hop2-1*, *hop2-2* and *hop2-3* alleles. Therefore, in Arabidopsis lack of SC formation between homologs does not prevent repair of all DSBs.

Cifuentes *et al.* (2013) looked at Arabidopsis wild type haploids and saw that they form DSBs normally, repair all DSBs, and form only intact univalents during segregation. The haploids also form DMC1 foci at 50% the level seen in diploids, proportional to their having 50% fewer chromosomes. Thus, in WT haploids, which lack synapsis, there are no COs, but all DSBs are repaired. As well, illegitimate exchanges with nonhomologous chromosomes are prevented in the wild type haploids. This contrasts sharply with the meiotic phenotype of *hop2-1* haploids where we found extensive entanglements with bridges and fragments forming once the chromosomes started segregating at anaphase I. Thus, our haploid *hop2-1* data strongly reinforce the concept that HOP2 has an important role in preventing illegitimate repair of DSBs that is separable from the role of HOP2-MND1 in assisting in SC and normal CO formation. The work of Uanschou *et al.* (2013) sheds additional light on the second role of HOP2 in preventing illegitimate reciprocal exchanges. They examined meiotic behavior in the *hop2-2* mutant allele that has a 147 bp deletion upstream of the start codon; plants homozygous for this allele make the normal protein but at dramatically lower levels than wild type. In this hypomorphic allele the positive role of HOP2-MND1 in promoting synapsis and homologous CO formation is lacking, but the role in preventing nonhomologous exchanges is intact; *hop2-2* plants generally form univalents and nonhomologous associations are rarely seen.

To think about the two different roles of HOP2 during meiosis we consider two different locations for DSBs at the time of DSB repair intermediates formation and completion. (1) Axially located DSBs – these would be DSBs that could originate either along the chromosome axis or be brought within the pairing axis, and which will be repaired using HR with a non-sister homologous chromosome in a manner that could give rise to a CO. (2) Radial DSBs – these would be DSBs that are found in the looped domains of the chromatin, which do not become axially located, and should be repaired by RAD51 using a sister chromatid or which might be repaired using DMC1 but in a manner that prevents second end capture and Double Holliday Junction (DHJ) intermediates.

In Arabidopsis the axial repairs that can generate crossovers require the concerted action of DMC1, the RAD51 complex of RAD51, RAD51C, XRCC2 (Su *et al.* 2017), and the assistance of MND1and HOP2 (Uanschou *et al.*, 2013). The role of MND1-HOP2 for axial events may be to help stabilize HR intermediates generated by the RecA-like recombinases (Zhao *et al.*, 2014, Moktan *et al.*, 2014, Kang *et al.*, 2015) and perhaps help drive DSB repair intermediates, which are not sufficiently homologous, toward synthesis dependent strand annealing and a non-CO outcome.

We propose that the <u>nonaxial</u>, radial DSB repairs must be actively prevented from using a nonhomologous chromosome for an HR repair path that could produce an illegitimate exchange, and that HOP2 has a role in this process. The need for HOP2 likely relates to the large excess of DSBs seen in eukaryotes that use the DSB pathway to achieve stabilized pairing, and their requirement for DMC1 activity in prophase I. We propose that HOP2’s role in preventing nonhomologous exchanges in radial regions in Arabidopsis is accomplished by effectively preventing DMC1 from acting in radial regions, and preventing it from creating DSB intermediates that get to the second end capture intermediate; this would prevent formation of DHJs in nonaxial regions. The behavior of single and double mutants of *dmc1 and hop2-1* in Arabidopsis support this hypothesis. The *dmc1* single mutant forms intact univalents with DSBs being repaired by RAD51 without illegitimate exchanges (Couteau *et al.* 1999); single mutants of *hop2-1* complete DSB repairs but many of the repairs create illegitimate exchanges (Stronghill *et al.* 2010). Relative to prevention of nonhomologous exchanges, the lack of DMC1 suppresses illegitimate exchanges seen in the *hop2-1* single mutant; the double mutant of *dmc1-hop2-1* mainly forms intact univalents (Uanschou *et al.* 2013). Our work with irradiated petal nuclei, also supports a role for HOP2 in preventing illegitimate exchanges outside of the context of meiosis.

How might HOP2 act? One important clue is revealed by the results of Uanschou *et al.*, (2013) in their study of the *hop2-2* hypomorphic allele, which has very reduced expression of the *hop2* gene. The *hop2-2* mutant has normal levels of DMC1 and RAD51 foci, but only forms univalents. These univalents do not show evidence of illegitimate exchanges. Thus, while a large amount of HOP2 is required to promote stabilized pairing and subsequent crossing over, a much smaller amount of HOP2 is required to prevent nonhomologous exchanges. Therefore, the role of HOP2 in preventing nonhomologous exchanges is unlikely to involve direct binding to DMC1 and altering its activity. It seems more likely that HOP2 in this second role is acting in more of a regulatory fashion; previously published work points in two possible directions. Zhang *et al.* (2019) have shown in mice that HOP2 can function to regulate the activity of the transcriptional regulator ATF4. Thus, one possibility is that HOP2 might regulate the production or longevity of another important protein that acts during meiosis. Possible candidates for interaction would be ATM and ATR. In Arabidopsis, single *atr* mutants appear to have a normal meiosis (Culligan and Britt 2008); *atm* single mutants have some nonhomologous associations and reduced fertility (Garcia *et al.* 2003), but *atm-atr* double mutants have extensive chromosome entanglements and are completely sterile (Culligan and Britt (2008). These authors conclude that ATR and ATM act redundantly during meiosis to prevent ectopic nonhomologous chromosome interactions. Thus, one possibility is that HOP2 may, in some manner, be regulating ATR and ATM.

Another intriguing possibility is that HOP2 impacts chromosome movements during prophase I. Sepsi and Schwarzacher (2020) reviewed the literature on chromosome-nuclear envelope tethering in plants and proposed that nuclear envelope bridge complexes control chromosome tethering to the nuclear envelope and subsequent chromosome movement during prophase I. They further consider that chromosome movement, as prophase I progresses, could serve to disrupt ectopic nonhomologous DSB repair intermediates prior to the commitment of the type of DSB repair intermediates that could produce a CO outcome. We previously did serial reconstructions of *hop2-1* and wild type nuclei (Pathan *et al.*, 2013). In addition to the fact that we found very little SC in the *hop2-1* mutant we also found abnormal structures on the nuclear envelope. The most striking difference observed was that in the mutant there exists a round structure approximately 1um in diameter that spanned the nuclear envelope. This anomalous structure was observed in all eight of the reconstructed mutant nuclei but was not seen in any of the twenty wild type reconstructions. It is possible that in *hop2-1* mutants, the apparatus for chromosome tethering and subsequent movement is disrupted and without the subsequent chromosome movements, ectopically-formed DSB early intermediates might be allowed to repair in a manner that can generate illegitimate exchanges. Recent technological innovations in Arabidopsis cytology that preserve three-dimensional structure and allow chromosome movements to be studied (Hurel *et al.* 2018) could shed light on this possibility.

## EXPERIMENTAL PROCEDURES

### Plant materials

Seeds of *A. thaliana* ecotypes Landsberg *erecta* (L*er*) and Columbia (Col), as well as the *mnd1* mutant and the haploid inducer line *cenh3-1*, were obtained from the Arabidopsis Biological Resource Center (http://www.arabidopsis.org/abrc). Seeds for *hop2-1* were kindly provided by Dr. Robert Sablowski of the John Innes Center (Schommer *et al.*, 2003). All seeds were sown on Premier PROMIX PGX (Plant Products), and plants were propagated at 22°C in Conviron AC60 environmental chambers for 16hour light/8hour dark periods employing fluorescent lighting at ~120μE/m^2^.

### Identification of Haploid Plants

Haploid plants were generated by crossing *hop2-1* heterozygotes (L*er* background) with *cenh3-1* (Columbia background) as described by Ravi *et al.* (2014). Sterile progenies were identified as possible haploids and karyotyping of the first flower buds, using the method of Mandácová and Lysak (2016), identified non-meiotic cells with only 5 chromosomes. DNA from rosette leaves of these plants was purified and used in SSLP analysis (see Table S2 for primers) to confirm that for each of the five chromosomes, L*er*-specific sequences were present. The *HOP2* allele status was assessed by PCR to determine the genotype of each haploid plant as having either the HOP2 wild type allele or the mutant *hop2-1* allele. We employed the wildtype primers (FOR 5’GCACTTGATAGTCTTGCTGATGCTG 3’ and BACK 5’CTCACCAATGTAATCCCTTCACG3’) to generate a 489bp product. The forward primer was used with the T-DNA primer GABI-KATo8549 (5’ GCTTTCGCCTATAAATACGACGG 3’) to generate a PCR product of ~340bp in samples containing an interrupted *HOP2* gene (*hop2-1* mutant). We generated at least five haploids having either a wildtype or mutant *HOP2* gene.

### Cytological analysis of haploid plants

Meiotic chromosome preparations from genotyped haploids were created as indicated in Mandácová and Lysak (2016), except our cell wall digestion was done for 2.5 hrs (vs their 3hrs) and we used 45% acetic acid (vs their 60%) in the dispersal of cellular material during spreading. Slides were analysed with a Zeiss Axiophot epi-fluorescent microscope using a 63X PlanApo lens and Sensicam CCD camera. Images were processed and saved with Northern Eclipse software. Figure images were generated using ImageJ (Image J 1.52a National Institute of Health). Statistical software R was used to perform Chi-square analysis.

### EdU labeling and γH2Ax immunolabeling experiments

EdU labeling was done as described by Armstrong (2013), with modifications as indicated in Stronghill *et al.*, 2014, with 4-5 inflorescences used for each time point. After fixation was completed, inflorescences were rinsed in meiotic buffer [15 mM pipes-NaOH (pH 6.8), 80 mM KCl, 20 mM NaCl, 0.5 mM EGTA, 2 mM EDTA, 0.15 mM spermine tetra-HCl, 0.05 mM spermidine, 1 mM dithiothreitol, 0.32 M sorbitol (pH = 8.2)]. Petals were removed from buds and the resulting pistil stamen units (PSUs) from each inflorescence were placed in a slide chamber containing meiotic buffer. Subsequent processing steps were done within the slide chamber, changing solutions as follows. Meiotic buffer was replaced with sodium citrate buffer (100mM, pH 5.0), then replaced with 200ul of cell wall enzyme mixture in citrate buffer (3% w/v each of cellulose, pectolyase, cytohelicase, glucanase and PVP-40). Covered slide chambers were incubated in a moist chamber at 40°C for 2 hrs.

Once the cell walls were softened, EdU detection was performed. PSUs in chambers were rinsed 3 times in phospho-buffered saline (PBS), pH 7.4, then incubated with PBS with 0.5% Triton-X for 1 hr. The PSUs were then rinsed in PBS once, then twice in PBS with 3% Bovine serum albumin (BSA). The PBS-BSA was then replaced with EdU Click-It colour reaction cocktail (as per manufacturer’s instructions found in the ‘Click It EdU Alexa Fluor 594’ – Imaging Kit, from Molecular Probes) for one hour at room temperature in the dark. All subsequent steps were done in dim lighting. The PSUs were rinsed twice in 3% BSA in PBS and kept in the dark until used for immunocytochemistry. For detection of γH2Ax, the PBS-BSA was replaced with incubation buffer (IB) that consisted of PBS-BSA with 0.1% Tween-20, then incubated with IB containing 150ul of anti-γH2AX (Ser 139) clone JBW301 (Millipore). Lidded chambers were incubated with primary antibody overnight at 4°C. The next day the solution was exchanged with PBS and incubated for 15 min on a flat rotator. PBS was replaced with 150ul of goat anti-mouse secondary antibody conjugated to Alexa-488 (Jackson ImmunoResearch) in IB and the slides were incubated in the dark at RT for 2 hrs. Slides were rinsed twice with PBS, each rinse for 15 min on a flat rotator.

Individual PSUs were transferred to a microscope slide containing 10ul of DAPI in Vectashield mounting media. Using a dissecting microscope at a low light setting, individual anthers were removed from the PSUs and spaced apart on the slide, other flower parts were removed, then a coverslip was added and slides were sealed with fingernail polish.

Slides were examined using a Quorum Wave-FX spinning disk confocal microscope and a Leica 63X oil immersion Plan-apo objective (NA 1.4). DAPI was detected using the blue laser line 406nm and pass band filter 460/50nm, Alexa 488 (γH2AX) was detected with the green laser line 491nm and pass band filter 525nm/50nm; EdU was detected with the red laser line 561nm) and pass band filter 620nm/60nm. The CCD camera used to capture images was a Hamamatsu Orca R2. The software used for image capture and analysis were Metamorph version 7.7.9.0 and Velocity version 6.1.1. Photoshop CS2 was used to label images and create figure montages.

### Staging criteria for EdU and γH2AX work

EdU and γH2Ax detection was done with anthers removed from individual flower buds. Within the anther-chambers each sac of meiocytes was examined and classified into relevant categories. The retention of intact cells within these sacs provides additional staging criteria using nucleolus shape and location, tapetum division status and meiocyte callose wall thickness (Stronghill *et al.* 2014). In brief, G2 and leptotene both have round nucleoli and a uninucleate tapetum, but leptotene is distinguishable from G2 by the presence of its uniformly, mildly condensed chromatin. Zygotene is distinguishable from leptotene because zygotene nucleoli becomes disk-shaped and peripheral (and pairing occurs in wild type). Additionally, at the onset of zygotene a wave of mitosis starts in the tapetal layer, such that filaments containing early zygotene nuclei have a layer of tapetal cells with frequent mitotic figures and creation of binucleate tapetal cells. By the end of zygotene, the tapetum is entirely binucleate. At the onset of pachytene, in wild types, pairing extends all along the chromosomes. In all of the genotypes we studied, regardless of pairing status, the nucleolus rounds up at the onset of pachytene and the neighboring tapetal layer is binucleate. As pachytene progresses the callose cell wall surrounding each meiocyte thickens. Thus, by combining chromosome attributes, nucleolus shape and location, division status of the tapetum and callose wall thickness we were able to readily distinguish G2, leptotene, zygotene, early pachytene and mid-late pachytene stages from each other. For the temporal distribution of γH2AX during prophase I reported for Figure 3, only meiotic nuclei that had incorporated EdU and were collected in the relevant time points for the early occurrence of each stage were used. For example, EdU-labeled leptotene nuclei were first seen in the L*er* genotype at the 6 hr time point and EdU labeled zygotene nuclei began appearing in the 12 hr time point. We collected data on EdU-labeled leptotenes of the L*er* genotype from the 6 -12 hr time points. Time points for the other meiotic stage for each genotype were similarly determined for Figure 3. For Figure S1 we did a similar analysis except that we assessed all meiotic cells for γH2AX status. Chi-Square tests (2×2) were done to test the significance of the observed differences between wild type and the mutants for the leptotene and zygotene stages.

### Mitotic Cell analysis

For the work on mitosis, petals were collected from buds in the size range of 0.4-0.5 mm as these buds contain petals with high mitotic indices (Smyth *et al.*, 1990). Irradiation of plants was carried out using a RAD SOURCE 2000 device. Plants with 1-2 open flowers were selected and positioned in the device to delivered 2.9 Gy/min, for 8.6 minutes such that the dosage received for each inflorescence was 25Gy. Irradiated plants were returned back to normal lighting conditions (growth chamber) after the treatment, and the buds were collected and processed after a recovery period. Unirradiated and irradiated buds were processed after the irradiated plants recovery period was complete.

Buds were harvested, the sepals were removed and the de-sepaled buds were individually placed in small depression slides for processing; all steps were carried out at RT unless stated otherwise. Fixation was for 2 hrs in 1ml of 3.7% paraformaldehyde dissolved in meiotic buffer, followed by three rinses in 10mM Sodium Citrate buffer pH 4.5 (CB). During incubations the slides were covered with a coverslip to prevent desiccation. After fixation, buds were rinsed in CB three times then the cell wall was softened by replacing the rinse buffer with 200 uls of 0.3% Pectolyase for 1 hr. Next, a post-fixation and permeabilization step was carried out by transferring the buds to 500ul of a solution of 3.7% of paraformaldehyde, 0.5% Triton-X in PBS for 15 min. Buds were rinsed three times using 3% BSA in PBS buffer in a depression slide. Petals were removed from the buds and transferred to a microscope slide with 10ul of Vectashield Mounting Media containing 4’,6-Diamidino-2-Phenylindole (DAPI) (1.5ug/mL) (Vector Laboratories, Canada). A 22×22 mm coverslip was added and mild pressure was applied to the coverslip, then the coverslip edges were sealed using nail polish.

Cytological analysis of the petal preparations was done using an Axioplan Fluorescence Microscope with a Zeiss 63X oil immersion PLAN-APO CHROMAT (NA 1.4) lens for the detailed imaging. An image capturing software Axiovision Ver. 4.8 was used for image captures. Statistical software R project was used to conduct data analysis, “Two sample” T-Test and 2×2 Chi-square tests were used to determine whether there were any significant differences between L*er* and *hop2-1*.

## ACKNOWLEDGEMENTS

The authors appreciate the advice, sharing of equipment and/or technical assistance of Dr. Sohee Kang and Bruno Chue and colleagues in the Plant Science group in the Department of Biological Sciences at the University of Toronto Scarborough. We also acknowledge undergraduates Nick Garrido, Diane Hamdan and Ellie Kubisz who played small but important roles in this research. Special thanks to Dr. Patti Stronghill who did most of the EdU and H2AX data collection. The research was funded by a grant from the Natural Sciences and Engineering Research Council of Canada (NSERC) to CAH and CDR, grant RGPGP‐2015‐00071 and a NSERC summer research award to YF-T.

## AUTHOR CONTRIBUTIONS

CAH designed the broad strokes of the experiments and wrote the paper. CDR developed and supervised all molecular biology experiments and provided major editorial assistance. PG did the preliminary work on generating and analyzing haploid plants and assisted in processing inflorescences for the EdU labeling. YF-T refined the cytological protocols, did genotyping, karyotyping and cytological analysis for the haploid experiments. CW developed protocols and collected all of the data for the petal irradiation experiments. JD performed genotyping and trained undergraduate personnel.

## SHORT LEGENDS FOR SUPPORTING MATERIALS

Supplemental Table 1 contains the entire data set for all meiotic filaments collected for the EdU incorporation and γHistone H2AX immunocytochemistry experiments.

Supplemental Table 2 provides the primers used in SSLP analysis to confirm the presence of each L*er* chromosome of the haploids.

Supplemental Figure 1 contains a summary of the time of appearance/disappearance of γH2AX foci in L*er*, *hop2-1*, Col and *mnd1* plants for each meiotic stage regardless of whether or not the nuclei had incorporated EdU (as compared to Figure 3 that only included the relevant meiotic stages if they had incorporated EdU).

Supplemental Figure 2 provides the data for karyotype, SSLP and PCR analyses to confirm the haploids as having five Ler chromosomes and having either *HOP2* or *hop2-1* alleles.

Supplemental Figure 3 provides a large section of one petal to indicate the level of mitotic activity.

## REFERENCES

Amiard, S., Charbonnel, C., Allain, E., Depeiges, A., White, C.I. and Gallego, M.E. (2010) Distinct Roles of the ATR Kinase and the Mre11-Rad50-Nbs1 Complex in the Maintenance of Chromosomal Stability in Arabidopsis. Plant Cell, 22, 3020–3033.

Armstrong, S. (2013) A time course for the analysis of meiotic progression in Arabidopsis thaliana. Methods Mol Biol, 990, 119–123.

Bishop, D.K., Park, D., Xu, L.Z. and Kleckner, N. (1992) DMC1-a meiosis-specific yeast homolog of *Escherichia coli* RecA required for recombination, synaptonemal complex formation, and cell cycle progression. Cell, 69, 439–456.

Brown, M.S., Grubb, J., Zhang, A., Rust, M.J. and Bishop, D.K. (2015) Small Rad51 and Dmc1 Complexes Often Co-occupy Both Ends of a Meiotic DNA Double Strand Break. Plos Genetics, 11.

Cifuentes, M., Rivard, M., Pereira, L., Chelysheva, L. and Mercier, R. (2013) Haploid Meiosis in Arabidopsis: Double-Strand Breaks Are Formed and Repaired but Without Synapsis and Crossovers. Plos One, 8.

Couteau, F., Belzile, F., Horlow, C., Grandjean, O., Vezon, D. and Doutriaux, M.P. (1999) Random chromosome segregation without meiotic arrest in both male and female meiocytes of a dmc1 mutant of Arabidopsis. Plant Cell, 11, 1623–1634.

Crickard, J.B. and Greene, E.C. (2018) The biochemistry of early meiotic recombination intermediates. Cell Cycle, 17, 2520–2530.

Crickard, J.B., Kwon, Y., Sung, P. and Greene, E.C. (2019) Dynamic interactions of the homologous pairing 2 (Hop2)-meiotic nuclear divisions 1 (Mnd1) protein complex with meiotic presynaptic filaments in budding yeast. Journal of Biological Chemistry, 294, 490–501.

Culligan, K.M. and Britt, A.B. (2008) Both ATM and ATR promote the efficient and accurate processing of programmed meiotic double-strand breaks. The Plant Journal, 55, 629–638.

Domenichini, S., Raynaud, C., Ni, D.A., Henry, Y. and Bergounioux, C. (2006) Atmnd1-Delta 1 is sensitive to gamma-irradiation and defective in meiotic DNA repair. DNA Repair, 5, 455–464.

Hurel, A., Phillips, D., Vrielynck, N., Mézard, C., Grelon, M. and Christophorou, N. (2018) A cytological approach to studying meiotic recombination and chromosome dynamics in Arabidopsis thaliana male meiocytes in three dimensions. The Plant Journal, 95, 385–396.

Jang, J.K., Sherizen, D.E., Bhagat, R., Manheim, E.A. and McKim, K.S. (2003) Relationship of DNA double-strand breaks to synapsis in *Drosophila*. Journal of Cell Science, 116, 3069–3077.

Kang, H.A., Shin, H.C., Kalantzi, A.S., Toseland, C.P., Kim, H.M., Gruber, S., Dal Peraro, M. and Oh, B.H. (2015) Crystal structure of Hop2-Mnd1 and mechanistic insights into its role in meiotic recombination. Nucleic Acids Research, 43, 3841–3856.

Keeney, S., Giroux, C.N. and Kleckner, N. (1997) Meiosis-Specific DNA Double-Strand Breaks Are Catalyzed by Spo11, a Member of a Widely Conserved Protein Family. Cell, 88, 375–384.

Keeney, S., Lange, J. and Mohibullah, N. (2014) Self-Organization of Meiotic Recombination Initiation: General Principles and Molecular Pathways. In Annual Review of Genetics, Vol 48 (Bassler, B.L. ed, pp. 187–214.

Kerzendorfer, C., Vignard, J., Pedrosa-Harand, A., Siwiec, T., Akimcheva, S., Jolivet, S., Sablowski, R., Armstrong, S., Schweizer, D., Mercier, R. and Schlogelhofer, P. (2006) The Arabidopsis thaliana MND1 homologue plays a key role in meiotic homologous pairing, synapsis and recombination. Journal of Cell Science, 119, 2486–2496.

Klepikova, A.V., Kasianov, A.S., Gerasimov, E.S., Logacheva, M.D. and Penin, A.A. (2016) A high resolution map of the *Arabidopsis thaliana* developmental transcriptome based on RNA-seq profiling. Plant J, 88, 1058–1070.

Kurzbauer, M.T., Uanschou, C., Chen, D. and Schlogelhofer, P. (2012) The Recombinases DMC1 and RAD51 Are Functionally and Spatially Separated during Meiosis in Arabidopsis. Plant Cell, 24, 2058–2070.

Leu, J.Y., Chua, P.R. and Roeder, G.S. (1998) The meiosis-specific hop2 protein of S-cerevisiae ensures synapsis between homologous chromosomes. Cell, 94, 375–386.

Lowndes, N.F. and Toh, G.W.L. (2005) DNA repair: The importance of phosphorylating histone H2AX. Current Biology, 15, R99–R102.

Mandáková, T. and Lysak, M.A. (2016) Chromosome Preparation for Cytogenetic Analyses in Arabidopsis. Current protocols in plant biology, 1, 43–51.

Mercier, R., Mézard, C., Jenczewski, E., Macaisne, N. and Grelon, M. (2015) The Molecular Biology of Meiosis in Plants. Annual Review of Plant Biology, 66, 297–327.

Moktan, H., Guiraldelli, M.F., Eyster, C.A., Zhao, W.X., Lee, C.Y., Mather, T., Camerini-Otero, R.D., Sung, P., Zhou, D.H.H. and Pezza, R.J. (2014) Solution Structure and DNA-binding Properties of the Winged Helix Domain of the Meiotic Recombination HOP2 Protein. Journal of Biological Chemistry, 289, 14682–14691.

Neale, M.J. and Keeney, S. (2006) Clarifying the mechanics of DNA strand exchange in meiotic recombination. Nature, 442, 153–158.

Panoli, A.P., Ravi, M., Sebastian, J., Nishal, B., Reddy, T.V., Marimuthu, M.P.A., Subbiah, V., Vijaybhaskar, V. and Siddiqi, I. (2006) AtMND1 is required for homologous pairing during meiosis in Arabidopsis. Bmc Molecular Biology, 7.

Pathan, N., Stronghill, P. and Hasenkampf, C. (2013) Transmission electron microscopy and serial reconstructions reveal novel meiotic phenotypes for the ahp2 mutant of Arabidopsis thaliana. Genome, 56, 139–145.

Petukhova, G.V., Pezza, R.J., Vanevski, F., Ploquin, M., Masson, J.Y. and Camerini-Otero, R.D. (2005) The Hop2 and Mnd1 proteins act in concert with Rad51 and Dmc1 in meiotic recombination. Nature Structural & Molecular Biology, 12, 449–453.

Petukhova, G.V., Romanienko, P.J. and Camerini-Otero, R.D. (2003) The Hop2 protein has a direct role in promoting interhomolog interactions during mouse meiosis. Developmental Cell, 5, 927–936.

Pittman, D.L., Cobb, J., Schimenti, K.J., Wilson, L.A., Cooper, D.M., Brignull, E., Handel, M.A. and Schimenti, J.C. (1998) Meiotic prophase arrest with failure of chromosome synapsis in mice deficient for Dmc1, a germline-specific RecA homolog. Molecular Cell, 1, 697–705.

Ploquin, M., Petukhova, G.V., Morneau, D., Dery, U., Bransi, A., Stasiak, A., Camerini-Otero, R.D. and Masson, J.Y. (2007) Stimulation of fission yeast and mouse Hop2-Mnd1 of the Dmc1 and Rad51 recombinases. Nucleic Acids Research, 35, 2719–2733.

Ranjha, L., Howard, S.M. and Cejka, P. (2018) Main steps in DNA double-strand break repair: an introduction to homologous recombination and related processes. Chromosoma, 127, 187–214.

Ravi, M., Marimuthu, M.P.A., Tan, E.H., Maheshwari, S., Henry, I.M., Marin-Rodriguez, B., Urtecho, G., Tan, J., Thornhill, K., Zhu, F., Panoli, A., Sundaresan, V., Britt, A.B., Comai, L. and Chan, S.W.L. (2014) A haploid genetics toolbox for Arabidopsis thaliana. Nature Communications, 5.

Sanchez-Moran, E., Santos, J.L., Jones, G.H. and Franklin, F.C.H. (2007) ASY1 mediates AtDMC1-dependent interhomolog recombination during meiosis in Arabidopsis. Genes & Development, 21, 2220–2233.

Sansam, C.L. and Pezza, R.J. (2015) Connecting by breaking and repairing: mechanisms of DNA strand exchange in meiotic recombination. The FEBS journal, 282, 2444–2457.

Schommer, C., Beven, A., Lawrenson, T., Shaw, P. and Sablowski, R. (2003) AHP2 is required for bivalent formation and for segregation of homologous chromosomes in Arabidopsis meiosis. Plant Journal, 36, 1–11.

Sepsi, A. and Schwarzacher, T. (2020) Chromosome-nuclear envelope tethering – a process that orchestrates homologue pairing during plant meiosis? Journal of Cell Science, 133.

Shi, W.Q., Tang, D., Shen, Y., Xue, Z.H., Zhang, F.F., Zhang, C., Ren, L.J., Liu, C.Z., Du, G.J., Li, Y.F., Yan, C.J. and Cheng, Z.K. (2019) OsHOP2 regulates the maturation of crossovers by promoting homologous pairing and synapsis in rice meiosis. New Phytologist, 222, 805–819.

Shinohara, M., Gasior, S.L., Bishop, D.K. and Shinohara, A. (2000) Tid1/Rdh54 promotes colocalization of Rad51 and Dmc1 during meiotic recombination. Proc. Natl. Acad. Sci. U. S. A., 97, 10814–10819.

Smyth, D.R., Bowman, J.L. and Meyerowitz, E.M. (1990) Early flower development in Arabidopsis. Plant Cell, 2, 755–767.

Stronghill, P., Pathan, N., Ha, H., Supijono, E. and Hasenkampf, C. (2010) Ahp2 (Hop2) function in Arabidopsis thaliana (Ler) is required for stabilization of close alignment and synaptonemal complex formation except for the two short arms that contain nucleolus organizer regions. Chromosoma, 119, 443–458.

Stronghill, P.E., Azimi, W. and Hasenkampf, C.A. (2014) A novel method to follow meiotic progression in Arabidopsis using confocal microscopy and 5-ethynyl-2 '-deoxyuridine labeling. Plant Methods, 10.

Su, H., Cheng, Z.H., Huang, J.Y., Lin, J., Copenhaver, G.P., Ma, H. and Wang, Y.X. (2017) Arabidopsis RAD51, RAD51C and XRCC3 proteins form a complex and facilitate RAD51 localization on chromosomes for meiotic recombination. Plos Genetics, 13.

Tsubouchi, H. and Roeder, G.S. (2003) The Importance of Genetic Recombination for Fidelity of Chromosome Pairing in Meiosis. Developmental Cell, 5, 915–925.

Uanschou, C., Ronceret, A., Von Harder, M., De Muyt, A., Vezon, D., Pereira, L., Chelysheva, L., Kobayashi, W., Kurumizaka, H., Schlogelhofer, P. and Grelon, M. (2013) Sufficient Amounts of Functional HOP2/MND1 Complex Promote Interhomolog DNA Repair but Are Dispensable for Intersister DNA Repair during Meiosis in Arabidopsis. Plant Cell, 25, 4924–4940.

Vignard, J., Siwiec, T., Chelysheva, L., Vrielynck, N., Gonord, F., Armstrong, S.J., Schlogelhofer, P. and Mercier, R. (2007) The interplay of RecA-related proteins and the MND1-HOP2 complex during meiosis in Arabidopsis thaliana. Plos Genetics, 3, 1894–1906.

Yao, Y., Li, X., Chen, W., Liu, H., Mi, L., Ren, D., Mo, A. and Lu, P. (2020) ATM Promotes RAD51-Mediated Meiotic DSB Repair by Inter-Sister-Chromatid Recombination in Arabidopsis. Frontiers in Plant Science, 11.

Zhao, W.X., Saro, D., Hammel, M., Kwon, Y., Xu, Y.Y., Rambo, R.P., Williams, G.J., Chi, P., Lu, L., Pezza, R.J., Camerini-Otero, R.D., Tainer, J.A., Wang, H.W. and Sung, P. (2014) Mechanistic insights into the role of Hop2-Mnd1 in meiotic homologous DNA pairing. Nucleic Acids Research, 42, 906–917.

