## Supplemental Table S1. for "The Arabidopsis *HOP2* Gene Has a Role in Preventing illegitimate Exchanges between Nonhomologous Chromosomes"

### Supplemental Table 1. EdU Incorporation and γH2AX Immunostaining for each Filament containing Meiotic Cells

**Guide to Table S1**

Intact inflorescences were placed in media with EdU for two hours, transferred to media without EdU, then individual inflorescences were removed at specific time points. Once removed from the media each inflorescence was fixed and processed. For cytology individual buds were removed, then the individual anthers (ant) of each bud had their filaments (fil) of ~30 meiotic cells extruded onto glass slides for analysis. Time point zero is the time EdU media was replaced with media with no EdU; this was two hours from the start of the experiment.

Table S1 is first organized by genotype with all L*er* data appearing before that of *hop2-1*, which appears before Col which appears before *mnd1*. Next Table S1 is organized by the timepoint when the inflorescences were removed from the media and processed for analysis.

For each anther collected, additional data is provided for whether or not the meiotic and tapetal cells had incorporated EdU, level and type of γH2AX foci immunostaining, meiotic stage, tapetum division status, nucleolus shape and position, and callose cell wall thickness.

For the level of γH2AX foci, level 0 indicates no foci; level 2 indicates 1-50 foci, level 3 indicates significantly more than 50 foci. The notation 'fbf' is used to indicate nuclei that had only 2-5 large round signals; this was seen only in early pachytene of L*er* and Col filaments.

For each genotype, the first buds seen in a timepoint for which EdU incorporation occurred, for that stage, are shown in bold and framed in a box.

The table is color coded to show which data was used to generate Figure 3 for each meiotic stage for γH2AX foci, with lightest yellow for G2, and each subsequent stage being a darker color, culminating with brown for mid - to- late pachytene.

Each filament of ~30 meiotic cells was considered as one data point entry; this file contains data for 2086 filaments.

### Ler data begins here

| Geno-type | Time point (hr) | Bud, Anther, Filament | # meiocyte with EdU signal/total | Level γH2AX signal | Meiotic stage. | Nucleolus shape & location | Amount Callose | % bi-nucleate. tapetum. | Tapetum labeled w/EdU |
|---|---|---|---|---|---|---|---|---|---|
| LER | 0 | **bud 1,** ant1, fil1 | 0/30 | 2 | zygotene | not visible | 1 | 1 to 25 | yes |
| LER | 0 | fil 2 | 0/30 | 1 | leptotene | circular , peripheral | 1 | 0 | no |
| LER | 0 | fil 3 | 0/30 | 1 | leptotene | circular , peripheral | 1 | 0 | no |
| LER | 0 | fil 4 | 0/30 | 0 | leptotene | circular , pericentric | 1 | 0 | no |
| LER | 0 | **bud 2,** ant1, fil1 | 0/30 | 2 | zygotene | not visible | 1 | 1 to 25 | yes |
| **LER** | **0** | **bud 3, ant1, fil1** | **10/30** | **0** | **G2** | **circular , centric** | **1** | **0** | **no** |
| **LER** | **0** | **fil 2** | **20/30** | **0** | **G2** | **circular , centric** | **1** | **0** | **no** |
| **LER** | **0** | **fil 3** | **20/30** | **0** | **G2** | **circular , centric** | **1** | **0** | **no** |
| LER | 0 | **bud 4,** ant1, fil1 | 0/30 | 0 | leptotene | circular , pericentric | 1 | 0 | no |
| LER | 0 | fil 2 | 0/30 | 0 | leptotene | circular , pericentric | 1 | 0 | no |
| LER | 0 | ant 2, fil 1 | 0/30 | 0 | G2 | circular , centric | 1 | 0 | no |
| LER | 0 | fil2 | 0/30 | 0 | G2 | circular , centric | 1 | 0 | no |
| LER | 0 | fil 3 | 0/30 | 0 | G2 | circular , centric | 1 | 0 | no |
| LER | 0 | ant 3, fil 1 | 0/30 | 0 | G2 | circular , centric | 1 | 0 | no |
| LER | 0 | ant 4, fil 1 | 0/30 | 0 | leptotene | circular , pericentric | 1 | 0 | no |
| LER | 0 | ant 5 fil 1 | 0/30 | 0 | leptotene | circular , pericentric | 1 | 0 | no |
| LER | 0 | fil 2 | 0/30 | 0 | leptotene | circular , pericentric | 1 | 0 | no |
| LER | 0 | **bud 5,** ant1, fil1 | 0/30 | fbf | eP | circular,peripheral | 1 | 100 | yes |
| LER | 0 | fil 2 | 0/30 | fbf | eP | circular,peripheral | 1 | 100 | yes |
| LER | 0 | fil 3 | 0/30 | fbf | eP | circular,peripheral | 1 | 100 | yes |
| LER | 0 | **bud 6,** ant1, fil1 | 0/30 | 0 | diplotene | not visible | N/A | 100 | no |
| LER | 0 | fil 2 | 0/30 | 0 | diplotene | not visible | N/A | 100 | no |

| | | | | | | | | | |
|---|---|---|---|---|---|---|---|---|---|
| LER | 0 | fil 3 | 0/30 | 0 | diplotene | not visible | N/A | 100 | no |
| LER | 0 | ant 2, fil 1 | 0/30 | 0 | diplotene | not visible | N/A | 100 | no |
| LER | 0 | fil2 | 0/30 | 0 | diplotene | not visible | N/A | 100 | no |
| LER | 0 | ant 3, fil 1 | 0/30 | 0 | diplotene | not visible | N/A | 100 | no |
| LER | 0 | ant 4, fil 1 | 0/30 | 0 | diplotene | not visible | N/A | 100 | no |
| LER | 0 | fil2 | 0/30 | 0 | diplotene | not visible | N/A | 100 | no |
| LER | 2 | **bud 1,** ant1, fil1 | 0/30 | 0 | G2 | circular , centric | 1 | 0 | no |
| LER | 2 | fil 2 | 0/30 | 0 | G2 | circular , centric | 1 | 0 | no |
| LER | 2 | ant 2, fil 1 | 0/30 | 0 | leptotene | circular , centric | 1 | 0 | no |
| LER | 2 | fil 2 | 0/30 | 0 | leptotene | circular , centric | 1 | 0 | no |
| LER | 2 | fil 3 | 0/30 | 0 | leptotene | circular , centric | 1 | 0 | no |
| LER | 2 | ant 4, fil 1 | 0/30 | 2 | zygotene | not visible | 1 | 1 to 25 | yes |
| LER | 2 | fil 2 | 0/30 | 2 | zygotene | not visible | 1 | 1 to 25 | yes |
| LER | 2 | ant 5, fil 1 | 0/30 | 2 | zygotene | not visible | 1 | 1 to 25 | yes |
| LER | 2 | fil 2 | 0/30 | 2 | zygotene | not visible | 1 | 1 to 25 | yes |
| LER | 2 | fil 3 | 0/30 | 2 | zygotene | not visible | 1 | 1 to 25 | yes |
| LER | 2 | fil 4 | 0/30 | 2 | zygotene | not visible | 1 | 1 to 25 | yes |
| LER | 2 | **bud 2,** ant1, fil1 | 0/30 | fbf | eP | circular,peripheral | 1 | 100 | yes |
| LER | 2 | fil 2 | 0/30 | 0 | tetrad | not visible | N/A | 100 | no |
| LER | 2 | fil 3 | 0/30 | 0 | tetrad | not visible | N/A | 100 | no |
| LER | 2 | **bud 3,** ant1, fil1 | 0/30 | 0 | leptotene | circular , centric | 1 | 0 | no |
| LER | 2 | fil 2 | 0/30 | 0 | leptotene | circular , centric | 1 | 0 | no |
| LER | 2 | fil 3 | 0/30 | 0 | leptotene | circular , centric | 1 | 0 | no |
| LER | 2 | ant 2, fil 1 | 0/30 | 0 | G2 | circular , centric | 1 | 0 | no |
| LER | 2 | ant 3, fil 1 | 0/30 | 0 | leptotene | circular , centric | 1 | 0 | no |
| LER | 2 | fil2 | 0/30 | 0 | leptotene | circular , centric | 1 | 0 | no |
| LER | 2 | ant 4, fil 1 | 0/30 | 0 | leptotene | circular , centric | 1 | 0 | no |
| LER | 2 | fil2 | 0/30 | 0 | leptotene | circular , centric | 1 | 0 | no |

| | | | | | | | | | |
|---|---|---|---|---|---|---|---|---|---|
| LER | 2 | fil 3 | 0/30 | 0 | leptotene | circular , centric | 1 | 0 | no |
| LER | 4 | **bud 1,** ant1, fil1 | 0/30 | 0 | leptotene | circular , centric | 1 | 0 | no |
| LER | 4 | fil 2 | 0/30 | 0 | leptotene | circular , centric | 1 | 0 | no |
| LER | 4 | ant 2, fil 1 | 0/30 | 2 | zygotene | not visible | 1 | 1 to 25 | yes |
| LER | 4 | fil2 | 0/30 | 2 | zygotene | not visible | 1 | 1 to 25 | yes |
| LER | 4 | ant 3, fil 1 | 0/30 | 2 | zygotene | not visible | 1 | 26-75 | yes |
| LER | 4 | fil2 | 0/30 | 2 | zygotene | not visible | 1 | 26-75 | yes |
| LER | 4 | ant 4, fil 1 | 0/30 | 2 | zygotene | not visible | 1 | 1 to 25 | yes |
| LER | 4 | **bud 2,** ant1, fil1 | 0/30 | 0 | leptotene | circular , centric | 1 | 0 | no |
| LER | 4 | fil 2 | 0/30 | 0 | leptotene | circular , centric | 1 | 0 | no |
| LER | 4 | **bud 3,** ant1, fil1 | 0/30 | 0 | G2 | circular , centric | 1 | 0 | no |
| LER | 4 | fil 2 | 0/30 | 0 | G2 | circular , centric | 1 | 0 | no |
| LER | 4 | **bud 4,** ant1, fil1 | 10/30 | 0 | G2 | circular , centric | 1 | 0 | no |
| LER | 4 | fil 2 | 20/30 | 0 | G2 | circular , centric | 1 | 0 | no |
| LER | 4 | ant 2, fil 1 | 0/30 | 2 | zygotene | not visible | 1 | 1 to 25 | yes |
| LER | 4 | fil2 | 0/30 | 2 | zygotene | not visible | 1 | 1 to 25 | yes |
| LER | 4 | **bud 5,** ant1, fil1 | 0/30 | 0 | G2 | circular , centric | 1 | 0 | no |
| LER | 4 | fil 2 | 0/30 | 0 | G2 | circular , centric | 1 | 0 | no |
| LER | 4 | ant 2, fil 1 | 0/30 | 0 | G2 | circular , centric | 1 | 0 | no |
| LER | 4 | fil2 | 0/30 | 0 | G2 | circular , centric | 1 | 0 | no |
| LER | 4 | ant 3, fil 1 | 0/30 | 0 | G2 | circular , centric | 1 | 0 | no |
| LER | 4 | fil2 | 0/30 | 0 | G2 | circular , centric | 1 | 0 | no |
| LER | 4 | fil 3 | 0/30 | 0 | G2 | circular , centric | 1 | 0 | no |
| LER | 4 | **bud 6,** ant1, fil1 | 0/30 | 0 | m-l P | circular,peripheral | 3 | 100 | yes |
| LER | 4 | fil 2 | 0/30 | 0 | m-l P | circular,peripheral | 3 | 100 | yes |
| LER | 4 | fil 3 | 0/30 | 0 | m-l P | circular,peripheral | 3 | 100 | yes |
| LER | 4 | ant 2, fil 1 | 0/30 | 0 | m-l P | circular,peripheral | 3 | 100 | yes |
| LER | 4 | fil 2 | 0/30 | 0 | m-l P | circular,peripheral | 3 | 100 | yes |

| | | | | | | | | | |
|---|---|---|---|---|---|---|---|---|---|
| LER | 4 | fil 3 | 0/30 | 0 | m-l P | circular,peripheral | 3 | 100 | yes |
| LER | 4 | ant 3, fil 1 | 0/30 | 0 | m-l P | circular,peripheral | 3 | 100 | yes |
| LER | 4 | fil2 | 0/30 | 0 | m-l P | circular,peripheral | 3 | 100 | yes |
| LER | 4 | **bud 7,** ant1, fil1 | 10/30 | 0 | G2 | circular , centric | 1 | 0 | no |
| LER | 4 | fil 2 | 5/30 | 0 | G2 | circular , centric | 1 | 0 | no |
| LER | 4 | ant 2, fil 1 | 0/30 | 0 | G2 | circular , centric | 1 | 0 | no |
| LER | 4 | fil2 | 0/30 | 0 | G2 | circular , centric | 1 | 0 | no |
| LER | 4 | **bud 8,** ant1, fil1 | 0/30 | 0 | m-l P | circular,peripheral | 3 | 100 | yes |
| LER | 4 | fil 2 | 0/30 | 0 | m-l P | circular,peripheral | 3 | 100 | yes |
| LER | 4 | ant 2, fil 1 | 0/30 | 0 | diakenesis | not visible | N/A | 100 | no |
| LER | 4 | fil 2 | 0/30 | 0 | diakenesis | not visible | N/A | 100 | no |
| LER | 4 | fil 3 | 0/30 | 0 | m-l P | circular,peripheral | 3 | 100 | yes |
| LER | 4 | fil 4 | 0/30 | 0 | m-l P | circular,peripheral | 3 | 100 | yes |
| LER | 6 | **bud 1,** ant1, fil1 | 0/30 | 0 | metaphase I | not visible | N/A | 100 | no |
| LER | 6 | fil 2 | 0/30 | 0 | metaphase I | not visible | N/A | 100 | no |
| LER | 6 | fil 3 | 0/30 | 0 | tetrad | not visible | N/A | 100 | no |
| LER | 6 | **bud 2,** ant1, fil1 | 0/30 | 0 | G2 | circular , centric | 1 | 0 | no |
| LER | 6 | fil 2 | 0/30 | 0 | G2 | circular , centric | 1 | 0 | no |
| LER | 6 | fil 3 | 0/30 | 0 | G2 | circular , centric | 1 | 0 | no |
| LER | 6 | **bud 3,** ant1, fil1 | 0/30 | 0 | leptotene | circular , centric | 1 | 0 | no |
| LER | 6 | fil 2 | 0/30 | 0 | leptotene | circular , centric | 1 | 0 | no |
| LER | 6 | ant 2, fil 1 | 0/30 | 0 | leptotene | circular , centric | 1 | 0 | no |
| LER | 6 | fil2 | 0/30 | 0 | leptotene | circular , centric | 1 | 0 | no |
| LER | 6 | fil 3 | 5/30 | 0 | G2 | circular , centric | 1 | 0 | no |
| LER | 6 | ant 3, fil 1 | 10/30 | 0 | G2 | circular , centric | 1 | 0 | no |
| LER | 6 | fil2 | 5/30 | 0 | G2 | circular , centric | 1 | 0 | no |
| LER | 6 | ant 4, fil 1 | 10/30 | 0 | G2 | circular , centric | 1 | 0 | no |
| LER | 6 | **bud 4,** ant1, fil1 | 0/30 | 0 | metaphase I | not visible | N/A | 100 | no |

| | | | | | | | | | |
|---|---|---|---|---|---|---|---|---|---|
| LER | 6 | fil 2 | 0/30 | 0 | metaphase I | not visible | N/A | 100 | no |
| **LER** | **6** | **bud 5, ant1, fil1** | **10/30** | **0** | **leptotene** | **circular , centric** | **1** | **0** | **no** |
| **LER** | **6** | **fil 2** | **20/30** | **0** | **leptotene** | **circular , centric** | **1** | **0** | **no** |
| **LER** | **6** | **fil 3** | **10/30** | **0** | **leptotene** | **circular , centric** | **1** | **0** | **no** |
| **LER** | **6** | **fil 4** | **5/30** | **0** | **leptotene** | **circular , centric** | **1** | **0** | **no** |
| LER | 8 | **bud 1,** ant1, fil1 | 0/30 | 0 | G2 | circular , centric | 1 | 0 | no |
| LER | 8 | fil 2 | 0/30 | 0 | G2 | circular , centric | 1 | 0 | no |
| LER | 8 | ant 2, fil 1 | 0/30 | 0 | G2 | circular , centric | 1 | 0 | no |
| LER | 8 | **bud 2,** ant1, fil1 | 0/30 | 0 | m-l P | circular,peripheral | 3 | 100 | yes |
| LER | 8 | fil 2 | 0/30 | 0 | m-l P | circular,peripheral | 3 | 100 | yes |
| LER | 8 | ant 2, fil 1 | 0/30 | 0 | m-l P | circular,peripheral | 3 | 100 | yes |
| LER | 8 | fil 2 | 0/30 | 0 | m-l P | circular,peripheral | 3 | 100 | yes |
| LER | 8 | **bud 3,** ant1, fil1 | 0/30 | 0 | G2 | circular , centric | 1 | 0 | no |
| LER | 8 | fil 2 | 0/30 | 0 | G2 | circular , centric | 1 | 0 | no |
| LER | 8 | ant 2, fil 1 | 0/30 | 2 | zygotene | not visible | 1 | 1 to 25 | yes |
| LER | 8 | fil2 | 0/30 | 2 | zygotene | not visible | 1 | 1 to 25 | yes |
| LER | 8 | fil 3 | 0/30 | 2 | zygotene | not visible | 1 | 1 to 25 | yes |
| LER | 8 | fil 4 | 0/30 | 2 | zygotene | not visible | 1 | 1 to 25 | yes |
| LER | 8 | ant 3, fil 1 | 0/30 | 0 | m-l P | circular,peripheral | 2 | 100 | no |
| LER | 8 | fil2 | 0/30 | 0 | m-l P | circular,peripheral | 3 | 100 | yes |
| LER | 8 | fil 3 | 0/30 | fbf | eP | circular,peripheral | 1 | 100 | yes |
| LER | 8 | fil 4 | 0/30 | fbf | eP | circular,peripheral | 1 | 100 | yes |
| LER | 8 | **bud 4,** ant1, fil1 | 0/30 | 0 | G2 | circular , centric | 1 | 0 | no |
| LER | 8 | fil 2 | 0/30 | 0 | G2 | circular , centric | 1 | 0 | no |
| LER | 8 | **bud 5,** ant1, fil1 | 0/30 | 0 | m-l P | circular,peripheral | 3 | 100 | yes |
| LER | 8 | fil 2 | 0/30 | 0 | m-l P | circular,peripheral | 3 | 100 | yes |
| LER | 8 | fil 3 | 0/30 | 0 | m-l P | circular,peripheral | 3 | 100 | yes |
| LER | 8 | fil 4 | 0/30 | 0 | m-l P | circular,peripheral | 3 | 100 | yes |

| | | | | | | | | | |
|---|---|---|---|---|---|---|---|---|---|
| LER | 8 | ant 2, fil 1 | 30/30 | 0 | leptotene | circular, pericentric | 1 | 0 | no |
| LER | 8 | fil 2 | 30/30 | 0 | leptotene | circular, pericentric | 1 | 0 | no |
| LER | 8 | ant 3, fil 1 | 0/30 | 0 | m-l P | circular,peripheral | 3 | 100 | no |
| LER | 8 | fil 2 | 0/30 | 0 | m-l P | circular,peripheral | 3 | 100 | no |
| LER | 8 | fil 3 | 0/30 | 0 | m-l P | circular,peripheral | 3 | 100 | no |
| LER | 8 | fil 4 | 0/30 | 0 | m-l P | circular,peripheral | 3 | 100 | no |
| LER | 8 | ant 4, fil 1 | 0/30 | 1 | zygotene | not visible | 1 | 76-99 | yes |
| LER | 8 | fil2 | 0/30 | 0 | m-l P | circular,peripheral | 2 | 100 | yes |
| LER | 8 | fil 3 | 0/30 | 0 | m-l P | circular,peripheral | 2 | 100 | yes |
| LER | 10 | **bud 1,** ant1, fil1 | 0/30 | 0 | metaphase I | not visible | N/A | 100 | no |
| LER | 10 | fil 2 | 0/30 | 0 | metaphase I | not visible | N/A | 100 | no |
| LER | 10 | ant 2, fil 1 | 0/30 | 0 | diakenesis | not visible | N/A | 100 | no |
| LER | 10 | fil2 | 0/30 | 0 | diakenesis | not visible | N/A | 100 | no |
| LER | 10 | **bud 2,** ant1, fil1 | 0/30 | 0 | G2 | circular , centric | 1 | 0 | no |
| LER | 10 | fil 2 | 0/30 | 0 | G2 | circular , centric | 1 | 0 | no |
| LER | 10 | ant 2, fil 1 | 0/30 | 0 | G2 | circular , centric | 1 | 0 | no |
| LER | 10 | fil2 | 5/30 | 0 | G2 | circular , centric | 1 | 0 | no |
| LER | 10 | **bud 3,** ant1, fil1 | 30/30 | 0 | leptotene | circular, pericentric | 1 | 0 | no |
| LER | 10 | fil 2 | 30/30 | 0 | leptotene | circular, pericentric | 1 | 0 | no |
| LER | 10 | ant2, fil 1 | 30/30 | 0 | leptotene | circular, pericentric | 1 | 0 | no |
| LER | 10 | fil 2 | 30/30 | 0 | leptotene | circular, pericentric | 1 | 0 | no |
| LER | 10 | ant 3, fil 1 | 15/30 | 0 | leptotene | circular, pericentric | 1 | 0 | no |
| LER | 10 | fil2 | 15/30 | 0 | leptotene | circular, pericentric | 1 | 0 | no |
| LER | 10 | **bud 4,** ant1, fil1 | 0/30 | 0 | leptotene | circular , pericentric | 1 | 0 | no |
| LER | 10 | fil 2 | 0/30 | 0 | leptotene | circular , pericentric | 1 | 0 | no |
| LER | 10 | **bud 5,** ant1, fil1 | 0/30 | 0 | diakenesis | not visible | N/A | 100 | no |
| LER | 10 | fil 2 | 0/30 | 0 | diakenesis | not visible | N/A | 100 | no |
| LER | 10 | fil 3 | 0/30 | 0 | diakenesis | not visible | N/A | 100 | no |

| | | | | | | | | | |
|---|---|---|---|---|---|---|---|---|---|
| LER | 10 | fil 4 | 0/30 | 0 | diakenesis | not visible | N/A | 100 | no |
| LER | 10 | **bud 6,** ant1, fil1 | 5/30 | 0 | leptotene | circular, pericentric | 1 | 0 | no |
| LER | 10 | fil 2 | 10/30 | 0 | leptotene | circular, pericentric | 1 | 0 | no |
| LER | 10 | ant2, fil 1 | 10/30 | 0 | leptotene | circular, pericentric | 1 | 0 | no |
| LER | 10 | fil 2 | 10/30 | 0 | leptotene | circular, pericentric | 1 | 0 | no |
| LER | 10 | ant 3, fil 1 | 10/30 | 1 | leptotene | circular , peripheral | 1 | 0 | no |
| LER | 10 | fil2 | 10/30 | 1 | leptotene | circular , peripheral | 1 | 0 | no |
| LER | 10 | fil 3 | 10/30 | 1 | leptotene | circular , peripheral | 1 | 0 | no |
| LER | 10 | fil 4 | 10/30 | 1 | leptotene | circular , peripheral | 1 | 0 | no |
| LER | 10 | ant 4, fil 1 | 20/30 | 1 | leptotene | circular , peripheral | 1 | 0 | no |
| LER | 10 | fil2 | 20/30 | 1 | leptotene | circular , peripheral | 1 | 0 | no |
| LER | 10 | **bud 7,** ant1, fil1 | 0/30 | 0 | m-l P | circular,peripheral | 3 | 100 | no |
| LER | 10 | fil 2 | 0/30 | 0 | m-l P | circular,peripheral | 3 | 100 | no |
| LER | 10 | **bud 8,** ant1, fil1 | 0/30 | 2 | eP | circular,peripheral | 1 | 100 | yes |
| LER | 10 | fil 2 | 0/30 | fbf | eP | circular,peripheral | 1 | 100 | yes |
| LER | 10 | fil 3 | 0/30 | 2 | zygotene | not visible | 1 | 26-75 | yes |
| LER | 10 | fil 4 | 0/30 | 2 | zygotene | not visible | 1 | 26-75 | yes |
| LER | 10 | ant 2, fil 1 | 0/30 | fbf | eP | circular,peripheral | 1 | 100 | yes |
| LER | 10 | fil2 | 0/30 | fbf | eP | circular,peripheral | 1 | 100 | yes |
| LER | 10 | fil 3 | 0/30 | fbf | eP | circular,peripheral | 1 | 100 | yes |
| LER | 10 | fil 4 | 0/30 | fbf | eP | circular,peripheral | 1 | 100 | yes |
| LER | 12 | **bud 1,** ant1, fil1 | 0/30 | 0 | m-l P | circular,peripheral | 3 | 100 | no |
| LER | 12 | fil 2 | 0/30 | 0 | m-l P | circular,peripheral | 3 | 100 | no |
| LER | 12 | ant 2, fil 1 | 0/30 | 0 | m-l P | circular,peripheral | 2 | 100 | yes |
| LER | 12 | fil2 | 0/30 | 0 | m-l P | circular,peripheral | 3 | 100 | no |
| LER | 12 | fil 3 | 0/30 | 0 | m-l P | circular,peripheral | 3 | 100 | no |
| LER | 12 | fil 4 | 0/30 | fbf | eP | circular,peripheral | 1 | 100 | yes |
| LER | 12 | ant 3, fil 1 | 0/30 | 0 | m-l P | circular,peripheral | 2 | 100 | yes |

| | | | | | | | | | |
|---|---|---|---|---|---|---|---|---|---|
| LER | 12 | **bud 2,** ant1, fil1 | 0/30 | 0 | m-l P | circular,peripheral | 3 | 100 | no |
| LER | 12 | fil 2 | 0/30 | 0 | m-l P | circular,peripheral | 3 | 100 | no |
| LER | 12 | ant 2, fil 1 | 0/30 | 0 | tetrad | not visible | N/A | 100 | no |
| LER | 12 | fil2 | 0/30 | 0 | tetrad | not visible | N/A | 100 | no |
| **LER** | **12** | **bud 3, ant1, fil1** | **10/30** | **2** | **zygotene** | **not visible** | **1** | **1 to 25** | **yes** |
| LER | 12 | ant 2, fil 1 | 0/30 | fbf | eP | circular,peripheral | 1 | 100 | yes |
| LER | 12 | fil2 | 0/30 | fbf | eP | circular,peripheral | 1 | 100 | yes |
| LER | 12 | ant 3, fil 1 | 30/30 | 1 | leptotene | circular , peripheral | 1 | 0 | no |
| LER | 12 | fil2 | 30/30 | 1 | leptotene | circular , peripheral | 1 | 0 | no |
| LER | 12 | fil 3 | 30/30 | 1 | leptotene | circular , peripheral | 1 | 0 | no |
| LER | 12 | **bud 4,** ant1, fil1 | 0/30 | 0 | G2 | circular , centric | 1 | 0 | no |
| LER | 12 | fil 2 | 0/30 | 0 | G2 | circular , centric | 1 | 0 | no |
| LER | 12 | fil 3 | 0/30 | 0 | G2 | circular , centric | 1 | 0 | no |
| LER | 14 | **bud 1,** ant1, fil1 | 15/30 | 0 | leptotene | circular, peripheral | 1 | 0 | no |
| LER | 14 | **bud 2,** ant1, fil1 | 30/30 | 1 | leptotene | circular , peripheral | 1 | 0 | no |
| LER | 14 | fil 2 | 30/30 | 1 | leptotene | circular, peripheral | 1 | 0 | no |
| LER | 14 | fil 3 | 30/30 | 1 | leptotene | circular, peripheral | 1 | 0 | no |
| LER | 14 | ant 2, fil 1 | 30/30 | 0 | leptotene | circular, pericentric | 1 | 0 | no |
| LER | 14 | fil2 | 30/30 | 0 | leptotene | circular, pericentric | 1 | 0 | no |
| LER | 14 | ant 3, fil 1 | 0/30 | 2 | zygotene | not visible | 1 | 26-75 | yes |
| LER | 14 | ant 4, fil 1 | 30/30 | 0 | leptotene | circular, pericentric | 1 | 0 | no |
| LER | 14 | fil2 | 30/30 | 0 | leptotene | circular, pericentric | 1 | 0 | no |
| LER | 14 | **bud 3,** ant1, fil1 | 10/30 | 0 | leptotene | circular, pericentric | 1 | 0 | no |
| LER | 14 | fil 2 | 10/30 | 0 | leptotene | circular, pericentric | 1 | 0 | no |
| LER | 14 | fil 3 | 10/30 | 0 | leptotene | circular, pericentric | 1 | 0 | no |
| LER | 14 | ant 2, fil 1 | 30/30 | 0 | leptotene | circular, pericentric | 1 | 0 | no |
| LER | 14 | ant3, fil 1 | 30/30 | 0 | leptotene | circular, pericentric | 1 | 0 | no |
| LER | 14 | ant4, fil 1 | 30/30 | 0 | leptotene | circular, pericentric | 1 | 0 | no |

| | | | | | | | | | |
|---|---|---|---|---|---|---|---|---|---|
| LER | 14 | fil 2 | 30/30 | 0 | leptotene | circular, pericentric | 1 | 0 | no |
| LER | 14 | ant 5, fil 1 | 0/30 | 2 | zygotene | not visible | 1 | 26-75 | yes |
| LER | 14 | fil2 | 0/30 | 2 | zygotene | not visible | 1 | 26-75 | yes |
| LER | 14 | **bud 4,** ant1, fil1 | 0/30 | 0 | metaphase I | not visible | N/A | 100 | no |
| LER | 14 | fil 2 | 0/30 | 0 | metaphase I | not visible | N/A | 100 | no |
| LER | 14 | ant 2, fil 1 | 0/30 | 0 | metaphase I | not visible | N/A | 100 | no |
| LER | 14 | fil2 | 0/30 | 0 | metaphase I | not visible | N/A | 100 | no |
| LER | 14 | **bud 5,** ant1, fil1 | 30/30 | 2 | zygotene | not visible | 1 | 1 to 25 | yes |
| LER | 14 | fil 2 | 20/30 | 2 | zygotene | not visible | 1 | 1 to 25 | yes |
| LER | 14 | **bud 6,** ant1, fil1 | 20/30 | 0 | leptotene | circular, pericentric | 1 | 0 | no |
| LER | 14 | fil 2 | 20/30 | 0 | leptotene | circular, pericentric | 1 | 0 | no |
| LER | 14 | fil 3 | 20/30 | 0 | leptotene | circular, pericentric | 1 | 0 | no |
| LER | 14 | ant 2, fil 1 | 20/30 | 0 | leptotene | circular, pericentric | 1 | 0 | no |
| LER | 14 | fil 2 | 20/30 | 0 | leptotene | circular, pericentric | 1 | 0 | no |
| LER | 14 | fil 3 | 10/30 | 0 | leptotene | circular, pericentric | 1 | 0 | no |
| LER | 14 | **bud 7,** ant1, fil1 | 20/30 | 0 | leptotene | circular, pericentric | 1 | 0 | no |
| LER | 14 | fil 2 | 20/30 | 0 | leptotene | circular, pericentric | 1 | 0 | no |
| LER | 14 | fil 3 | 10/30 | 0 | leptotene | circular, pericentric | 1 | 0 | no |
| LER | 14 | fil 4 | 5/30 | 0 | leptotene | circular, pericentric | 1 | 0 | no |
| LER | 14 | **bud 8,** ant1, fil1 | 30/30 | 0 | leptotene | circular, pericentric | 1 | 0 | no |
| LER | 14 | fil 2 | 30/30 | 0 | leptotene | circular, pericentric | 1 | 0 | no |
| LER | 14 | fil 3 | 30/30 | 0 | leptotene | circular, pericentric | 1 | 0 | no |
| LER | 14 | fil 4 | 30/30 | 0 | leptotene | circular, pericentric | 1 | 0 | no |
| LER | 14 | ant 2, fil 1 | 30/30 | 0 | leptotene | circular, pericentric | 1 | 0 | no |
| LER | 14 | fil 2 | 30/30 | 0 | leptotene | circular, pericentric | 1 | 0 | no |
| LER | 14 | fil 3 | 30/30 | 0 | leptotene | circular, pericentric | 1 | 0 | no |
| LER | 14 | ant 3, fil1 | 30/30 | 0 | leptotene | circular, pericentric | 1 | 0 | no |
| LER | 14 | fil 2 | 30/30 | 0 | leptotene | circular, pericentric | 1 | 0 | no |
| LER | 14 | **bud 9,** ant1, fil1 | 30/30 | 0 | leptotene | circular , centric | 1 | 0 | no |

| | | | | | | | | | |
|---|---|---|---|---|---|---|---|---|---|
| LER | 14 | fil 2 | 30/30 | 0 | leptotene | circular , centric | 1 | 0 | no |
| LER | 14 | fil 3 | 30/30 | 0 | leptotene | circular , centric | 1 | 0 | no |
| LER | 14 | fil 4 | 30/30 | 0 | leptotene | circular , centric | 1 | 0 | no |
| LER | 14 | ant 2, fil 1 | 30/30 | 0 | leptotene | circular , centric | 1 | 0 | no |
| LER | 14 | fil 2 | 30/30 | 0 | leptotene | circular , centric | 1 | 0 | no |
| LER | 14 | ant 3, fil 1 | 0/30 | 0 | G2 | circular , centric | 1 | 0 | no |
| LER | 14 | ant 4, fil 1 | 30/30 | 0 | leptotene | circular, pericentric | 1 | 0 | no |
| LER | 14 | fil2 | 30/30 | 0 | leptotene | circular, pericentric | 1 | 0 | no |
| LER | 14 | fil 3 | 30/30 | 0 | leptotene | circular, pericentric | 1 | 0 | no |
| LER | 14 | fil 4 | 30/30 | 0 | leptotene | circular, pericentric | 1 | 0 | no |
| LER | 16 | **bud 1,** ant1, fil1 | 0/30 | 0 | m-l P | circular,peripheral | 2 | 100 | yes |
| LER | 16 | fil 2 | 0/30 | 0 | m-l P | circular,peripheral | 2 | 100 | yes |
| LER | 16 | ant 2, fil 1 | 0/30 | 0 | m-l P | circular,peripheral | 2 | 100 | yes |
| LER | 16 | fil2 | 0/30 | 0 | m-l P | circular,peripheral | 2 | 100 | yes |
| LER | 16 | **bud 2,** ant1, fil1 | 20/30 | 2 | zygotene | not visible | 1 | 26-75 | yes |
| LER | 16 | fil 2 | 0/30 | fbf | eP | circular,peripheral | 1 | 100 | yes |
| LER | 16 | fil 3 | 20/30 | 2 | zygotene | not visible | 1 | 26-75 | yes |
| LER | 16 | ant 2, fil 1 | 20/30 | 2 | zygotene | not visible | 1 | 1 to 25 | yes |
| LER | 16 | fil2 | 15/30 | 2 | zygotene | not visible | 1 | 1 to 25 | yes |
| LER | 16 | ant 3, fil 1 | 0/30 | fbf | eP | circular,peripheral | 1 | 100 | yes |
| LER | 16 | fil2 | 0/30 | fbf | eP | circular,peripheral | 1 | 100 | yes |
| LER | 16 | fil 3 | 0/30 | fbf | eP | circular,peripheral | 1 | 100 | yes |
| LER | 16 | fil 4 | 0/30 | fbf | eP | circular,peripheral | 1 | 100 | yes |
| LER | 16 | **bud 3,** ant1, fil1 | 25/30 | 2 | zygotene | not visible | 1 | 26-75 | yes |
| LER | 16 | **bud 4,** ant1, fil1 | 0/30 | 0 | G2 | circular , centric | 1 | 0 | no |
| LER | 16 | fil 2 | 0/30 | 0 | G2 | circular , centric | 1 | 0 | no |
| LER | 16 | fil 3 | 0/30 | 0 | G2 | circular , centric | 1 | 0 | no |
| LER | 16 | ant 2, fil 1 | 0/30 | 0 | G2 | circular , centric | 1 | 0 | no |

| | | | | | | | | | |
|---|---|---|---|---|---|---|---|---|---|
| LER | 16 | fil2 | 0/30 | 0 | G2 | circular , centric | 1 | 0 | no |
| LER | 16 | **bud 5,** ant1, fil1 | 0/30 | 0 | m-l P | circular,peripheral | 3 | 100 | no |
| LER | 16 | fil 2 | 0/30 | 0 | m-l P | circular,peripheral | 3 | 100 | no |
| LER | 16 | **bud 6,** ant1, fil1 | 0/30 | 0 | metaphase I | not visible | N/A | 100 | no |
| LER | 16 | ant 2, fil 1 | 0/30 | 0 | metaphase I | not visible | N/A | 100 | no |
| LER | 16 | **bud 7,** ant1, fil1 | 20/30 | 0 | leptotene | circular , centric | 1 | 0 | no |
| LER | 16 | fil 2 | 25/30 | 0 | leptotene | circular , centric | 1 | 0 | no |
| LER | 16 | ant 2, fil 1 | 30/30 | 0 | leptotene | circular, pericentric | 1 | 0 | no |
| LER | 16 | fil2 | 20/30 | 0 | leptotene | circular, pericentric | 1 | 0 | no |
| LER | 16 | fil 3 | 30/30 | 0 | leptotene | circular, pericentric | 1 | 0 | no |
| LER | 16 | **bud 8,** ant1, fil1 | 0/30 | 0 | tetrad | not visible | N/A | 100 | no |
| LER | 16 | **bud 9,** ant1, fil1 | 0/30 | 0 | G2 | circular , centric | 1 | 0 | no |
| LER | 16 | fil 2 | 0/30 | 0 | G2 | circular , centric | 1 | 0 | no |
| LER | 18 | **bud 1,** ant1, fil1 | 20/30 | 0 | leptotene | circular, pericentric | 1 | 0 | no |
| LER | 18 | fil 2 | 15/30 | 0 | leptotene | circular, pericentric | 1 | 0 | no |
| LER | 18 | ant2, fil 1 | 30/30 | 0 | leptotene | circular, pericentric | 1 | 0 | no |
| LER | 18 | fil 2 | 30/30 | 0 | leptotene | circular, pericentric | 1 | 0 | no |
| LER | 18 | ant 3, fil 1 | 0/30 | 0 | G2 | circular , centric | 1 | 0 | no |
| LER | 18 | fil2 | 0/30 | 0 | G2 | circular , centric | 1 | 0 | no |
| LER | 18 | ant 4, fil 1 | 0/30 | 0 | G2 | circular , centric | 1 | 0 | no |
| LER | 18 | fil2 | 0/30 | 0 | G2 | circular , centric | 1 | 0 | no |
| LER | 18 | ant 5, fil 1 | 5/30 | 0 | leptotene | circular , centric | 1 | 0 | no |
| LER | 18 | fil2 | 5/30 | 0 | leptotene | circular , centric | 1 | 0 | no |
| LER | 18 | **bud 2,** ant1, fil1 | 0/30 | 0 | G2 | circular , centric | 1 | 0 | no |
| LER | 18 | fil 2 | 0/30 | 0 | G2 | circular , centric | 1 | 0 | no |
| LER | 18 | fil 3 | 0/30 | 0 | G2 | circular , centric | 1 | 0 | no |
| LER | 18 | ant 2, fil 1 | 0/30 | 0 | G2 | circular , centric | 1 | 0 | no |
| LER | 18 | ant 3, fil 1 | 0/30 | 0 | G2 | circular , centric | 1 | 0 | no |

| | | | | | | | | | |
|---|---|---|---|---|---|---|---|---|---|
| LER | 18 | fil2 | 0/30 | 0 | G2 | circular , centric | 1 | 0 | no |
| LER | 18 | ant 4, fil 1 | 0/30 | 0 | G2 | circular , centric | 1 | 0 | no |
| LER | 18 | fil2 | 0/30 | 0 | G2 | circular , centric | 1 | 0 | no |
| LER | 18 | **bud 3,** ant1, fil1 | 0/30 | 0 | tetrad | not visible | N/A | 100 | no |
| LER | 18 | ant 2, fil 1 | 0/30 | 0 | m-l P | circular,peripheral | 2 | 100 | yes |
| LER | 18 | fil2 | 0/30 | 0 | m-l P | circular,peripheral | 2 | 100 | yes |
| LER | 18 | fil 3 | 0/30 | 0 | m-l P | circular,peripheral | 2 | 100 | yes |
| LER | 18 | **bud 4,** ant1, fil1 | 30/30 | 0 | leptotene | circular, pericentric | 1 | 0 | no |
| LER | 18 | fil 2 | 30/30 | 0 | leptotene | circular, pericentric | 1 | 0 | no |
| LER | 18 | fil 3 | 30/30 | 0 | leptotene | circular, pericentric | 1 | 0 | no |
| LER | 18 | **bud 5,** ant1, fil1 | 0/30 | 0 | m-l P | circular,peripheral | 2 | 100 | yes |
| LER | 18 | fil 2 | 0/30 | 0 | m-l P | circular,peripheral | 2 | 100 | yes |
| LER | 18 | **bud 6,** ant1, fil1 | 0/30 | 0 | metaphase I | not visible | N/A | 100 | no |
| LER | 18 | fil 2 | 0/30 | 0 | metaphase I | not visible | N/A | 100 | no |
| LER | 18 | fil 3 | 0/30 | 0 | metaphase I | not visible | N/A | 100 | no |
| LER | 18 | **bud 7,** ant1, fil1 | 0/30 | 0 | G2 | circular , centric | 1 | 0 | no |
| LER | 18 | fil 2 | 0/30 | 0 | G2 | circular , centric | 1 | 0 | no |
| LER | 18 | fil 3 | 0/30 | 0 | G2 | circular , centric | 1 | 0 | no |
| LER | 18 | ant 2, fil 1 | 0/30 | 0 | G2 | circular , centric | 1 | 0 | no |
| LER | 18 | fil2 | 0/30 | 0 | G2 | circular , centric | 1 | 0 | no |
| LER | 18 | fil 3 | 0/30 | 0 | G2 | circular , centric | 1 | 0 | no |
| LER | 18 | ant 3, fil 1 | 30/30 | 0 | leptotene | circular , centric | 1 | 0 | no |
| LER | 18 | fil2 | 30/30 | 0 | leptotene | circular , centric | 1 | 0 | no |
| LER | 18 | **bud 8,** ant1, fil1 | 30/30 | 1 | zygotene | not visible | 1 | 76-99 | yes |
| LER | 18 | fil 2 | 30/30 | 2 | zygotene | not visible | 1 | 26-75 | yes |
| LER | 18 | fil 3 | 30/30 | 2 | zygotene | not visible | 1 | 26-75 | yes |
| LER | 18 | fil 4 | 30/30 | 2 | zygotene | not visible | 1 | 26-75 | yes |
| **LER** | **18** | **ant 2, fil 1** | **10/30** | **fbf** | **eP** | **circular,peripheral** | **1** | **100** | **yes** |
| **LER** | **18** | **fil2** | **5/30** | **fbf** | **eP** | **circular,peripheral** | **1** | **100** | **yes** |

| | | | | | | | | | |
|---|---|---|---|---|---|---|---|---|---|
| **LER** | **18** | **fil 3** | **5/30** | **fbf** | **eP** | **circular,peripheral** | **1** | **100** | **yes** |
| LER | 18 | ant 3, fil 1 | 30/30 | 2 | zygotene | not visible | 1 | 26-75 | yes |
| LER | 18 | fil2 | 30/30 | 2 | zygotene | not visible | 1 | 26-75 | yes |
| LER | 18 | fil 3 | 30/30 | 2 | zygotene | not visible | 1 | 26-75 | yes |
| LER | 18 | **bud 9,** ant1, fil1 | 30/30 | 1 | leptotene | circular , peripheral | 1 | 0 | no |
| LER | 18 | fil 2 | 30/30 | 1 | leptotene | circular , peripheral | 1 | 0 | no |
| LER | 18 | fil 3 | 30/30 | 1 | leptotene | circular , peripheral | 1 | 0 | no |
| LER | 18 | **bud 10,** ant1, fil1 | 0/30 | 0 | G2 | circular , centric | 1 | 0 | no |
| LER | 18 | fil 2 | 0/30 | 0 | G2 | circular , centric | 1 | 0 | no |
| LER | 18 | fil 3 | 0/30 | 0 | G2 | circular , centric | 1 | 0 | no |
| LER | 18 | ant 2, fil 1 | 0/30 | 0 | G2 | circular , centric | 1 | 0 | no |
| LER | 18 | fil2 | 0/30 | 0 | G2 | circular , centric | 1 | 0 | no |
| LER | 18 | ant 3, fil 1 | 0/30 | 0 | G2 | circular , centric | 1 | 0 | no |
| LER | 18 | fil2 | 0/30 | 0 | G2 | circular , centric | 1 | 0 | no |
| LER | 20 | **bud 1,** ant1, fil1 | 0/30 | 0 | metaphase I | not visible | N/A | 100 | no |
| LER | 20 | fil 2 | 0/30 | 0 | metaphase I | not visible | N/A | 100 | no |
| LER | 20 | ant 2, fil 1 | 0/30 | 0 | metaphase I | not visible | N/A | 100 | no |
| LER | 20 | **bud 2,** ant1, fil1 | 15/30 | 2 | zygotene | not visible | 1 | 1 to 25 | no |
| LER | 20 | fil 2 | 30/30 | 0 | leptotene | circular, pericentric | 1 | 0 | no |
| LER | 20 | ant 2, fil 1 | 10/30 | 2 | zygotene | not visible | 1 | 1 to 25 | no |
| LER | 20 | fil2 | 10/30 | 2 | zygotene | not visible | 1 | 1 to 25 | no |
| LER | 20 | fil 3 | 10/30 | 2 | zygotene | not visible | 1 | 1 to 25 | no |
| LER | 20 | **bud 3,** ant1, fil1 | 0/30 | 0 | anaphase I | not visible | N/A | 100 | no |
| LER | 20 | fil 2 | 0/30 | 0 | anaphase I | not visible | N/A | 100 | no |
| LER | 20 | ant 2, fil 1 | 0/30 | 0 | diakenesis | not visible | N/A | 100 | no |
| LER | 20 | fil2 | 0/30 | 0 | metaphase I | not visible | N/A | 100 | no |
| LER | 20 | ant 3, fil 1 | 0/30 | 0 | anaphase I | not visible | N/A | 100 | no |
| LER | 20 | ant 4, fil 1 | 0/30 | 0 | tetrad | not visible | N/A | 100 | no |

| | | | | | | | | | |
|---|---|---|---|---|---|---|---|---|---|
| LER | 20 | ant 5, fil 1 | 10/30 | fbf | eP | circular,peripheral | 1 | 100 | yes |
| LER | 20 | fil2 | 10/30 | fbf | eP | circular,peripheral | 1 | 100 | yes |
| LER | 20 | **bud 4,** ant1, fil1 | 30/30 | 0 | leptotene | circular, pericentric | 1 | 0 | no |
| LER | 20 | fil 2 | 30/30 | 0 | leptotene | circular, pericentric | 1 | 0 | no |
| LER | 20 | fil 3 | 30/30 | 0 | leptotene | circular, pericentric | 1 | 0 | no |
| LER | 20 | fil 4 | 30/30 | 0 | leptotene | circular, pericentric | 1 | 0 | no |
| LER | 20 | ant 2, fil 1 | 30/30 | 0 | leptotene | circular, pericentric | 1 | 0 | no |
| LER | 20 | ant 3, fil 1 | 30/30 | 0 | leptotene | circular, pericentric | 1 | 0 | no |
| LER | 20 | fil 2 | 30/30 | 0 | leptotene | circular, pericentric | 1 | 0 | no |
| LER | 20 | **bud 5,** ant1, fil1 | 30/30 | 0 | leptotene | circular, pericentric | 1 | 0 | no |
| LER | 20 | fil 2 | 30/30 | 0 | leptotene | circular, pericentric | 1 | 0 | no |
| LER | 20 | ant 2, fil 1 | 30/30 | 0 | leptotene | circular, pericentric | 1 | 0 | no |
| LER | 20 | fil 2 | 30/30 | 0 | leptotene | circular, pericentric | 1 | 0 | no |
| LER | 20 | ant 2, fil 1 | 30/30 | 0 | leptotene | circular, pericentric | 1 | 0 | no |
| LER | 20 | fil2 | 30/30 | 0 | leptotene | circular, pericentric | 1 | 0 | no |
| LER | 22 | **bud 1,** ant1, fil1 | 0/30 | 0 | leptotene | circular , centric | 1 | 0 | no |
| LER | 22 | fil 2 | 0/30 | 0 | leptotene | circular , centric | 1 | 0 | no |
| LER | 22 | ant2, fil 3 | 0/30 | 0 | leptotene | circular , centric | 1 | 0 | no |
| LER | 22 | ant 3, fil 1 | 0/30 | 0 | leptotene | circular , centric | 1 | 0 | no |
| LER | 22 | fil2 | 0/30 | 0 | leptotene | circular , centric | 1 | 0 | no |
| LER | 22 | ant 4, fil 1 | 0/30 | 0 | leptotene | circular , centric | 1 | 0 | no |
| LER | 22 | fil2 | 0/30 | 0 | leptotene | circular , centric | 1 | 0 | no |
| LER | 22 | fil 3 | 0/30 | 0 | leptotene | circular , centric | 1 | 0 | no |
| LER | 22 | **bud 2,** ant1, fil1 | 30/30 | 2 | zygotene | not visible | 1 | 26-75 | yes |
| LER | 22 | fil 2 | 30/30 | 2 | zygotene | not visible | 1 | 26-75 | yes |
| LER | 22 | fil 3 | 30/30 | 2 | zygotene | not visible | 1 | 26-75 | yes |
| LER | 22 | ant 2, fil 1 | 25/30 | 0 | leptotene | circular, pericentric | 1 | 0 | no |
| LER | 22 | fil2 | 20/30 | 0 | leptotene | circular, pericentric | 1 | 0 | no |

| | | | | | | | | | |
|---|---|---|---|---|---|---|---|---|---|
| LER | 22 | **bud 3,** ant1, fil1 | 0/30 | 0 | anaphase I | not visible | N/A | 100 | no |
| LER | 22 | fil 2 | 0/30 | 0 | anaphase I | not visible | N/A | 100 | no |
| LER | 22 | fil 3 | 0/30 | 0 | anaphase I | not visible | N/A | 100 | no |
| LER | 24 | **bud 1,** ant1, fil1 | 0/30 | 0 | m-l P | circular,peripheral | 3 | 100 | yes |
| LER | 24 | fil 2 | 0/30 | 0 | m-l P | circular,peripheral | 3 | 100 | yes |
| LER | 24 | ant 2, fil 1 | 25/30 | 2 | zygotene | not visible | 1 | 1 to 25 | no |
| LER | 24 | fil2 | 20/30 | 2 | zygotene | not visible | 1 | 1 to 25 | no |
| LER | 24 | fil 3 | 20/30 | 2 | zygotene | not visible | 1 | 1 to 25 | no |
| LER | 24 | **bud 2,** ant1, fil1 | 0/30 | 0 | m-l P | circular,peripheral | 3 | 100 | yes |
| LER | 24 | fil 2 | 0/30 | 0 | m-l P | circular,peripheral | 3 | 100 | yes |
| LER | 24 | fil 3 | 0/30 | 0 | m-l P | circular,peripheral | 3 | 100 | yes |
| LER | 24 | fil 4 | 0/30 | 0 | m-l P | circular,peripheral | 3 | 100 | yes |
| LER | 24 | **bud 3,** ant1, fil1 | 0/30 | 0 | anaphase I | not visible | N/A | 100 | no |
| LER | 24 | fil 2 | 0/30 | 0 | anaphase I | not visible | N/A | 100 | no |
| LER | 26 | **bud 1,** ant1, fil1 | 0/30 | 0 | G2 | circular , centric | 1 | 0 | no |
| LER | 26 | ant 2, fil 1 | 0/30 | 0 | G2 | circular , centric | 1 | 0 | no |
| LER | 26 | fil2 | 0/30 | 0 | G2 | circular , centric | 1 | 0 | no |
| LER | 26 | fil 3 | 0/30 | 0 | G2 | circular , centric | 1 | 0 | no |
| LER | 26 | **bud 2,** ant1, fil1 | 20/30 | 0 | leptotene | circular, pericentric | 1 | 0 | no |
| LER | 26 | fil 2 | 20/30 | 0 | leptotene | circular, pericentric | 1 | 0 | no |
| LER | 26 | ant 2, fil 1 | 15/30 | 0 | leptotene | circular, pericentric | 1 | 0 | no |
| LER | 26 | fil2 | 20/30 | 0 | leptotene | circular, pericentric | 1 | 0 | no |
| LER | 26 | fil 3 | 25/30 | 0 | leptotene | circular, pericentric | 1 | 0 | no |
| LER | 26 | ant 3, fil 1 | 30/30 | 0 | leptotene | circular, pericentric | 1 | 0 | no |
| LER | 26 | fil2 | 20/30 | 2 | zygotene | not visible | 1 | 1 to 25 | no |
| LER | 26 | fil 3 | 20/30 | 2 | zygotene | not visible | 1 | 1 to 25 | no |
| LER | 26 | fil 4 | 30/30 | 0 | leptotene | circular, pericentric | 1 | 0 | no |
| LER | 26 | ant 4, fil 1 | 10/30 | 0 | leptotene | circular, pericentric | 1 | 0 | no |

| | | | | | | | | | |
|---|---|---|---|---|---|---|---|---|---|
| LER | 26 | fil2 | 10/30 | 0 | leptotene | circular, pericentric | 1 | 0 | no |
| LER | 26 | **bud 3,** ant1, fil1 | 0/30 | 0 | G2 | circular , centric | 1 | 0 | no |
| LER | 26 | fil 2 | 0/30 | 0 | G2 | circular , centric | 1 | 0 | no |
| LER | 26 | ant 2, fil 1 | 0/30 | 0 | G2 | circular , centric | 1 | 0 | no |
| LER | 26 | fil2 | 0/30 | 0 | G2 | circular , centric | 1 | 0 | no |
| LER | 26 | ant 3, fil 1 | 0/30 | 0 | G2 | circular , centric | 1 | 0 | no |
| LER | 26 | fil2 | 0/30 | 0 | G2 | circular , centric | 1 | 0 | no |
| LER | 26 | **bud 4,** ant1, fil1 | 30/30 | 0 | leptotene | circular, pericentric | 1 | 0 | no |
| LER | 26 | fil 2 | 30/30 | 0 | leptotene | circular, pericentric | 1 | 0 | no |
| LER | 26 | ant 2, fil 1 | 20/30 | 0 | leptotene | circular, pericentric | 1 | 0 | no |
| LER | 26 | fil2 | 25/30 | 0 | leptotene | circular, pericentric | 1 | 0 | no |
| LER | 26 | fil 3 | 20/30 | 1 | leptotene | circular , peripheral | 1 | 0 | no |
| LER | 26 | fil 4 | 20/30 | 1 | leptotene | circular , peripheral | 1 | 0 | no |
| LER | 26 | **bud 5,** ant1, fil1 | 30/30 | fbf | eP | circular,peripheral | 1 | 100 | yes |
| LER | 26 | fil 2 | 30/30 | fbf | eP | circular,peripheral | 1 | 100 | yes |
| LER | 26 | ant 2, fil 1 | 0/30 | 0 | m-l P | circular,peripheral | 3 | 100 | yes |
| LER | 26 | fil2 | 0/30 | 0 | m-l P | circular,peripheral | 3 | 100 | yes |
| LER | 26 | **bud 6,** ant1, fil1 | 30/30 | 1 | leptotene | circular , peripheral | 1 | 0 | no |
| LER | 26 | fil 2 | 30/30 | 1 | leptotene | circular , peripheral | 1 | 0 | no |
| LER | 26 | ant 2, fil 1 | 30/30 | 1 | leptotene | circular , peripheral | 1 | 0 | no |
| LER | 26 | fil2 | 30/30 | 1 | leptotene | circular , peripheral | 1 | 0 | no |
| LER | 26 | ant 3, fil 1 | 0/30 | 0 | leptotene | circular , centric | 1 | 0 | no |
| LER | 26 | fil2 | 0/30 | 0 | leptotene | circular , centric | 1 | 0 | no |
| LER | 26 | fil 3 | 0/30 | 0 | leptotene | circular , centric | 1 | 0 | no |
| LER | 26 | ant 4, fil 1 | 0/30 | 0 | leptotene | circular , centric | 1 | 0 | no |
| LER | 26 | fil2 | 0/30 | 0 | leptotene | circular , centric | 1 | 0 | no |
| LER | 26 | **bud 7,** ant1, fil1 | 30/30 | 0 | leptotene | circular, pericentric | 1 | 0 | no |
| LER | 26 | fil 2 | 30/30 | 0 | leptotene | circular, pericentric | 1 | 0 | no |
| LER | 26 | fil 3 | 30/30 | 0 | leptotene | circular, pericentric | 1 | 0 | no |

| | | | | | | | | | |
|---|---|---|---|---|---|---|---|---|---|
| LER | 26 | fil 4 | 30/30 | 0 | leptotene | circular, pericentric | 1 | 0 | no |
| LER | 28 | **bud 1,** ant1, fil1 | 0/30 | 0 | leptotene | circular , centric | 1 | 0 | no |
| LER | 28 | fil 2 | 0/30 | 0 | leptotene | circular , centric | 1 | 0 | no |
| LER | 28 | ant 2, fil 1 | 0/30 | 0 | leptotene | circular , centric | 1 | 0 | no |
| LER | 28 | fil2 | 0/30 | 0 | leptotene | circular , centric | 1 | 0 | no |
| LER | 28 | ant 3, fil 1 | 0/30 | 0 | leptotene | circular , centric | 1 | 0 | no |
| LER | 28 | fil2 | 0/30 | 0 | leptotene | circular , centric | 1 | 0 | no |
| LER | 28 | **bud 2,** ant1, fil1 | 0/30 | 0 | metaphase I | not visible | N/A | 100 | no |
| LER | 28 | fil 2 | 0/30 | 0 | metaphase I | not visible | N/A | 100 | no |
| LER | 28 | fil 3 | 0/30 | 0 | metaphase I | not visible | N/A | 100 | no |
| LER | 28 | bud 3, ant1, fil1 | 30/30 | 0 | m-l P | circular,peripheral | 3 | 100 | yes |
| LER | 28 | fil 2 | 30/30 | 0 | m-l P | circular,peripheral | 3 | 100 | yes |
| LER | 28 | fil 3 | 30/30 | 0 | m-l P | circular,peripheral | 3 | 100 | yes |
| LER | 28 | ant 2, fil 1 | 30/30 | 0 | m-l P | circular,peripheral | 3 | 100 | yes |
| LER | 28 | fil2 | 30/30 | 0 | m-l P | circular,peripheral | 3 | 100 | yes |
| LER | 28 | fil 3 | 30/30 | 0 | m-l P | circular,peripheral | 3 | 100 | yes |
| LER | 28 | **bud 4,** ant1, fil1 | 0/30 | 0 | metaphase I | not visible | N/A | 100 | no |
| LER | 28 | fil 2 | 0/30 | 0 | metaphase I | not visible | N/A | 100 | no |
| LER | 28 | fil 3 | 0/30 | 0 | metaphase I | not visible | N/A | 100 | no |
| LER | 30 | **bud 1,** ant1, fil1 | 30/30 | 2 | zygotene | not visible | 1 | 26-75 | yes |
| LER | 30 | fil 2 | 30/30 | 2 | zygotene | not visible | 1 | 26-75 | yes |
| LER | 30 | ant 2, fil 1 | 30/30 | 2 | zygotene | not visible | 1 | 26-75 | yes |
| LER | 30 | fil2 | 30/30 | 2 | zygotene | not visible | 1 | 26-75 | yes |
| LER | 30 | ant 3, fil 1 | 30/30 | 2 | zygotene | not visible | 1 | 26-75 | yes |
| LER | 30 | fil2 | 30/30 | 2 | zygotene | not visible | 1 | 26-75 | yes |
| LER | 30 | fil 3 | 30/30 | 2 | zygotene | not visible | 1 | 26-75 | yes |
| LER | 30 | **bud 2,** ant1, fil1 | 0/30 | 0 | G2 | circular , centric | 1 | 0 | no |
| LER | 30 | fil 2 | 0/30 | 0 | G2 | circular , centric | 1 | 0 | no |

| | | | | | | | | | |
|---|---|---|---|---|---|---|---|---|---|
| LER | 30 | fil 3 | 0/30 | 0 | G2 | circular , centric | 1 | 0 | no |
| LER | 30 | fil 4 | 0/30 | 0 | G2 | circular , centric | 1 | 0 | no |
| LER | 30 | ant 2, fil 1 | 0/30 | 0 | tetrad | not visible | N/A | 100 | no |
| LER | 30 | fil2 | 0/30 | 0 | tetrad | not visible | N/A | 100 | no |
| LER | 30 | fil 3 | 0/30 | 0 | tetrad | not visible | N/A | 100 | no |
| LER | 30 | **bud 3,** ant1, fil1 | 30/30 | 0 | m-l P | circular,peripheral | 3 | 100 | yes |
| LER | 30 | fil 2 | 30/30 | 0 | m-l P | circular,peripheral | 3 | 100 | yes |
| LER | 30 | fil 3 | 30/30 | 0 | m-l P | circular,peripheral | 3 | 100 | yes |

### hop2-1 data begins here

| Geno-type | Time point (hr) | Bud, Anther, Filament | # meiocyte with EdU signal/total | Level γH2AX signal | Meiotic stage. | Nucleolus shape & location | Amount Callose | % bi-nucleate. tapetum. | Tapetum labeled w/EdU |
|---|---|---|---|---|---|---|---|---|---|
| hop2-1 | 0 | **bud 1**, ant1, fil1 | 0/30 | 1 | zygotene | not visible | 1 | 1 to 25 | yes |
| hop2-1 | 0 | fil 2 | 0/30 | 1 | zygotene | not visible | 1 | 1 to 25 | yes |
| hop2-1 | 0 | fil 3 | 0/30 | 1 | zygotene | not visible | 1 | 1 to 25 | yes |
| hop2-1 | 0 | fil 4 | 0/30 | 1 | zygotene | not visible | 1 | 1 to 25 | yes |
| hop2-1 | 0 | **bud 2**, ant1, fil1 | 0/30 | 1 | leptotene | circular , peripheral | 1 | 0 | no |
| hop2-1 | 0 | fil 2 | 0/30 | 1 | zygotene | not visible | 1 | 1 to 25 | yes |
| hop2-1 | 0 | ant 2, fil 1 | 0/30 | 1 | zygotene | not visible | 1 | 1 to 25 | yes |
| hop2-1 | 0 | ant 3, fil 1 | 0/30 | 1 | zygotene | not visible | 1 | 1 to 25 | yes |
| hop2-1 | 0 | fil2 | 0/30 | 1 | zygotene | not visible | 1 | 1 to 25 | yes |
| hop2-1 | 0 | fil 3 | 0/30 | 1 | zygotene | not visible | 1 | 1 to 25 | yes |
| hop2-1 | 0 | **bud 3**, ant1, fil1 | 0/30 | 0 | G2 | circular , centric | 1 | 0 | no |
| hop2-1 | 0 | fil 2 | 0/30 | 0 | G2 | circular , centric | 1 | 0 | no |
| hop2-1 | 0 | fil 3 | 0/30 | 0 | G2 | circular , centric | 1 | 0 | no |

| | | | | | | | | | |
|---|---|---|---|---|---|---|---|---|---|
| hop2-1 | 0 | ant 2, fil 1 | 0/30 | 0 | G2 | circular , centric | 1 | 0 | no |
| hop2-1 | 0 | fil2 | 0/30 | 0 | G2 | circular , centric | 1 | 0 | no |
| hop2-1 | 0 | fil 3 | 0/30 | 0 | G2 | circular , centric | 1 | 0 | no |
| hop2-1 | 0 | fil 4 | 0/30 | 0 | G2 | circular , centric | 1 | 0 | no |
| hop2-1 | 0 | **bud 4**, ant1, fil1 | 0/30 | 0 | anaphase I I | not visible | N/A | 100 | no |
| hop2-1 | 0 | ant 2, fil 1 | 0/30 | 0 | m-l P | circular,peripheral | 3 | 100 | no |
| hop2-1 | 0 | fil2 | 0/30 | 0 | m-l P | circular,peripheral | 3 | 100 | no |
| hop2-1 | 0 | fil 3 | 0/30 | 0 | m-l P | circular,peripheral | 3 | 100 | no |
| hop2-1 | 0 | fil 4 | 0/30 | 0 | m-l P | circular,peripheral | 3 | 100 | no |
| hop2-1 | 0 | ant 3, fil 1 | 0/30 | 1 | eP | circular,peripheral | 1 | 100 | no |
| hop2-1 | 0 | fil2 | 0/30 | 1 | eP | circular,peripheral | 1 | 100 | no |
| hop2-1 | 0 | **bud 5**, ant1, fil1 | 0/30 | 0 | m-l P | circular,peripheral | 2 | 100 | no |
| hop2-1 | 0 | fil 2 | 0/30 | 0 | m-l P | circular,peripheral | 2 | 100 | no |
| hop2-1 | 0 | fil 3 | 0/30 | 0 | m-l P | circular,peripheral | 2 | 100 | no |
| hop2-1 | 0 | fil 4 | 0/30 | 0 | m-l P | circular,peripheral | 2 | 100 | no |
| hop2-1 | 0 | ant 2, fil 1 | 0/30 | 1 | zygotene | not visible | 1 | 76-99 | yes |
| hop2-1 | 0 | fil2 | 0/30 | 1 | zygotene | not visible | 1 | 76-99 | yes |
| hop2-1 | 0 | fil 3 | 0/30 | 1 | zygotene | not visible | 1 | 76-99 | yes |
| hop2-1 | 0 | fil 4 | 0/30 | 1 | zygotene | not visible | 1 | 76-99 | yes |
| hop2-1 | 2 | **bud 1**, ant1, fil1 | 0/30 | 0 | tetrad | not visible | N/A | 100 | no |
| hop2-1 | 2 | fil 2 | 0/30 | 0 | tetrad | not visible | N/A | 100 | no |
| hop2-1 | 2 | ant 2, fil 1 | 0/30 | 0 | tetrad | not visible | N/A | 100 | no |
| hop2-1 | 2 | fil2 | 0/30 | 0 | tetrad | not visible | N/A | 100 | no |
| hop2-1 | 2 | ant 3, fil 1 | 0/30 | 0 | tetrad | not visible | N/A | 100 | no |
| hop2-1 | 2 | fil2 | 0/30 | 0 | tetrad | not visible | N/A | 100 | no |
| hop2-1 | 2 | **bud 2**, ant1, fil1 | 0/30 | 1 | eP | circular,peripheral | 1 | 100 | no |
| hop2-1 | 2 | fil 2 | 0/30 | 1 | eP | circular,peripheral | 1 | 100 | no |
| hop2-1 | 2 | ant 2, fil 1 | 0/30 | 2 | zygotene | not visible | 1 | 1 to 25 | yes |

| | | | | | | | | | |
|---|---|---|---|---|---|---|---|---|---|
| hop2-1 | 2 | fil2 | 0/30 | 2 | zygotene | not visible | 1 | 1 to 25 | yes |
| hop2-1 | 2 | ant 3, fil 1 | 0/30 | 0 | m-l P | circular,peripheral | 3 | 100 | no |
| hop2-1 | 2 | ant 4, fil 1 | 0/30 | 2 | zygotene | not visible | 1 | 1 to 25 | yes |
| hop2-1 | 2 | fil2 | 0/30 | 2 | zygotene | not visible | 1 | 1 to 25 | yes |
| hop2-1 | 2 | fil 3 | 0/30 | 2 | zygotene | not visible | 1 | 1 to 25 | yes |
| hop2-1 | 2 | ant 5, fil 1 | 0/30 | 2 | zygotene | not visible | 1 | 1 to 25 | yes |
| hop2-1 | 2 | fil2 | 0/30 | 2 | zygotene | not visible | 1 | 1 to 25 | yes |
| hop2-1 | 2 | ant 6, fil 1 | 0/30 | 0 | m-l P | circular,peripheral | 3 | 100 | no |
| hop2-1 | 2 | fil2 | 0/30 | 0 | m-l P | circular,peripheral | 3 | 100 | no |
| **hop2-1** | **2** | **bud 3, ant1, fil1** | **30/30** | **0** | **G2** | **circular , centric** | **1** | **0** | **no** |
| **hop2-1** | **2** | **fil 2** | **30/30** | **0** | **G2** | **circular , centric** | **1** | **0** | **no** |
| **hop2-1** | **2** | **fil 3** | **30/30** | **0** | **G2** | **circular , centric** | **1** | **0** | **no** |
| **hop2-1** | **2** | **ant 2, fil 1** | **30/30** | **0** | **G2** | **circular , centric** | **1** | **0** | **no** |
| **hop2-1** | **2** | **fil2** | **30/30** | **0** | **G2** | **circular , centric** | **1** | **0** | **no** |
| hop2-1 | 2 | **bud 4**, ant1, fil1 | 0/30 | 1 | eP | circular,peripheral | 1 | 100 | no |
| hop2-1 | 2 | fil 2 | 0/30 | 1 | eP | circular,peripheral | 1 | 100 | no |
| hop2-1 | 2 | **bud 5**, ant1, fil1 | 0/30 | 0 | leptotene | circular , centric | 1 | 0 | no |
| hop2-1 | 2 | fil 2 | 0/30 | 0 | leptotene | circular , centric | 1 | 0 | no |
| hop2-1 | 2 | fil 3 | 0/30 | 0 | leptotene | circular , centric | 1 | 0 | no |
| hop2-1 | 2 | **bud 6**, ant1, fil1 | 0/30 | 1 | zygotene | not visible | 1 | 1 to 25 | yes |
| hop2-1 | 2 | fil 2 | 0/30 | 1 | zygotene | not visible | 1 | 1 to 25 | yes |
| hop2-1 | 2 | fil 3 | 0/30 | 1 | zygotene | not visible | 1 | 1 to 25 | yes |
| hop2-1 | 2 | fil 4 | 0/30 | 1 | zygotene | not visible | 1 | 1 to 25 | yes |
| hop2-1 | 2 | ant 2, fil 1 | 0/30 | 1 | zygotene | not visible | 1 | 1 to 25 | yes |
| hop2-1 | 2 | fil2 | 0/30 | 1 | zygotene | not visible | 1 | 1 to 25 | yes |
| hop2-1 | 2 | fil 3 | 0/30 | 1 | zygotene | not visible | 1 | 1 to 25 | yes |
| hop2-1 | 2 | fil 4 | 0/30 | 1 | zygotene | not visible | 1 | 1 to 25 | yes |
| hop2-1 | 2 | **bud 7**, ant1, fil1 | 0/30 | 0 | anaphase I | not visible | N/A | 100 | no |

| | | | | | | | | | |
|---|---|---|---|---|---|---|---|---|---|
| hop2-1 | 2 | fil 2 | 0/30 | 0 | anaphase I | not visible | N/A | 100 | no |
| hop2-1 | 4 | **bud 1**, ant1, fil1 | 0/30 | 1 | zygotene | not visible | 1 | 1 to 25 | yes |
| hop2-1 | 4 | fil 2 | 0/30 | 1 | zygotene | not visible | 1 | 1 to 25 | yes |
| hop2-1 | 4 | fil 3 | 0/30 | 1 | zygotene | not visible | 1 | 1 to 25 | yes |
| hop2-1 | 4 | ant 2, fil 1 | 30/30 | 0 | G2 | circular , centric | 1 | 0 | no |
| hop2-1 | 4 | fil2 | 30/30 | 0 | G2 | circular , centric | 1 | 0 | no |
| hop2-1 | 4 | **bud 2**, ant1, fil1 | 0/30 | 1 | zygotene | not visible | 1 | 1 to 25 | yes |
| hop2-1 | 4 | fil 2 | 0/30 | 1 | zygotene | not visible | 1 | 1 to 25 | yes |
| hop2-1 | 4 | fil 3 | 0/30 | 1 | zygotene | not visible | 1 | 1 to 25 | yes |
| hop2-1 | 4 | **bud 3**, ant1, fil1 | 0/30 | 2 | zygotene | not visible | 1 | 26-75 | no |
| hop2-1 | 4 | fil 2 | 0/30 | 2 | zygotene | not visible | 1 | 26-75 | no |
| hop2-1 | 4 | ant 2, fil 1 | 0/30 | 0 | m-l P | circular,peripheral | 3 | 100 | no |
| hop2-1 | 4 | fil2 | 0/30 | 0 | m-l P | circular,peripheral | 3 | 100 | no |
| hop2-1 | 4 | ant 3, fil 1 | 0/30 | 0 | leptotene | circular , centric | 1 | 0 | yes |
| hop2-1 | 4 | fil2 | 0/30 | 0 | leptotene | circular , centric | 1 | 0 | yes |
| hop2-1 | 4 | ant 4, fil 1 | 0/30 | 1 | eP | circular,peripheral | 1 | 100 | no |
| hop2-1 | 4 | fil2 | 0/30 | 1 | eP | circular,peripheral | 1 | 100 | no |
| hop2-1 | 4 | fil 3 | 0/30 | 1 | eP | circular,peripheral | 1 | 100 | no |
| hop2-1 | 4 | **bud 4**, ant1, fil1 | 0/30 | 0 | leptotene | circular, pericentric | 1 | 0 | yes |
| hop2-1 | 4 | fil 2 | 0/30 | 0 | leptotene | circular, pericentric | 1 | 0 | yes |
| hop2-1 | 4 | fil 3 | 0/30 | 0 | leptotene | circular, pericentric | 1 | 0 | yes |
| hop2-1 | 4 | ant 2, fil 1 | 0/30 | 2 | zygotene | not visible | 1 | 26-75 | no |
| hop2-1 | 4 | fil2 | 0/30 | 1 | zygotene | not visible | 1 | 1 to 25 | yes |
| hop2-1 | 4 | fil 3 | 0/30 | 1 | zygotene | not visible | 1 | 1 to 25 | yes |
| hop2-1 | 4 | fil 4 | 0/30 | 1 | zygotene | not visible | 1 | 1 to 25 | yes |
| hop2-1 | 4 | **bud 5**, ant1, fil1 | 0/30 | 2 | zygotene | not visible | 1 | 76-99 | no |
| hop2-1 | 4 | fil 2 | 0/30 | 2 | zygotene | not visible | 1 | 26-75 | no |
| hop2-1 | 4 | ant 2, fil 1 | 0/30 | 1 | zygotene | not visible | 1 | 1 to 25 | yes |

| | | | | | | | | | |
|---|---|---|---|---|---|---|---|---|---|
| hop2-1 | 4 | fil2 | 0/30 | 2 | zygotene | not visible | 1 | 1 to 25 | yes |
| hop2-1 | 4 | **bud 6**, ant1, fil1 | 0/30 | 2 | zygotene | not visible | 1 | 1 to 25 | yes |
| hop2-1 | 4 | fil 2 | 0/30 | 2 | zygotene | not visible | 1 | 1 to 25 | yes |
| hop2-1 | 4 | ant 2, fil 1 | 0/30 | 2 | zygotene | not visible | 1 | 1 to 25 | yes |
| hop2-1 | 4 | fil2 | 0/30 | 2 | zygotene | not visible | 1 | 1 to 25 | yes |
| hop2-1 | 4 | fil 3 | 0/30 | 2 | zygotene | not visible | 1 | 1 to 25 | yes |
| hop2-1 | 4 | **bud 7**, ant1, fil1 | 0/30 | 0 | anaphase I | not visible | N/A | 100 | no |
| hop2-1 | 4 | fil 2 | 0/30 | 0 | diplotene | not visible | N/A | 100 | no |
| hop2-1 | 4 | ant 2, fil 1 | 0/30 | 0 | diplotene | not visible | N/A | 100 | no |
| hop2-1 | 4 | fil2 | 0/30 | 0 | m-l P | circular,peripheral | 3 | 100 | no |
| hop2-1 | 4 | fil 3 | 0/30 | 0 | m-l P | circular,peripheral | 3 | 100 | no |
| hop2-1 | 6 | **bud 1**, ant1, fil1 | 0/30 | 1 | zygotene | not visible | 1 | 1 to 25 | yes |
| hop2-1 | 6 | fil 2 | 0/30 | 1 | zygotene | not visible | 1 | 1 to 25 | yes |
| hop2-1 | 6 | fil 3 | 0/30 | 1 | zygotene | not visible | 1 | 1 to 25 | yes |
| hop2-1 | 6 | fil 4 | 0/30 | 1 | zygotene | not visible | 1 | 1 to 25 | yes |
| hop2-1 | 6 | ant 2, fil 1 | 0/30 | 1 | zygotene | not visible | 1 | 1 to 25 | yes |
| hop2-1 | 6 | fil2 | 0/30 | 2 | zygotene | not visible | 1 | 1 to 25 | yes |
| hop2-1 | 6 | fil 3 | 0/30 | 2 | zygotene | not visible | 1 | 1 to 25 | yes |
| hop2-1 | 6 | fil 4 | 0/30 | 2 | zygotene | not visible | 1 | 1 to 25 | yes |
| **hop2-1** | **6** | **ant 3, fil 1** | **30/30** | **0** | **leptotene** | **circular, centric** | **1** | **0** | **no** |
| hop2-1 | 6 | ant 4, fil 1 | 0/30 | 1 | zygotene | not visible | 1 | 1 to 25 | yes |
| hop2-1 | 6 | fil2 | 0/30 | 1 | zygotene | not visible | 1 | 1 to 25 | yes |
| hop2-1 | 6 | fil 3 | 0/30 | 1 | zygotene | not visible | 1 | 1 to 25 | yes |
| hop2-1 | 6 | fil 4 | 0/30 | 2 | zygotene | not visible | 1 | 1 to 25 | yes |
| hop2-1 | 6 | bud 2, ant1, fil1 | 30/30 | 0 | leptotene | circular, centric | 1 | 0 | no |
| hop2-1 | 6 | fil 2 | 30/30 | 0 | leptotene | circular, centric | 1 | 0 | no |
| hop2-1 | 6 | ant 2, fil 1 | 0/30 | 1 | zygotene | not visible | 1 | 1 to 25 | yes |
| hop2-1 | 6 | fil2 | 0/30 | 1 | zygotene | not visible | 1 | 1 to 25 | yes |

| | | | | | | | | | |
|---|---|---|---|---|---|---|---|---|---|
| hop2-1 | 6 | fil 3 | 0/30 | 1 | zygotene | not visible | 1 | 1 to 25 | yes |
| hop2-1 | 6 | ant 3, fil 1 | 0/30 | 1 | zygotene | not visible | 1 | 1 to 25 | yes |
| hop2-1 | 6 | fil2 | 0/30 | 1 | zygotene | not visible | 1 | 1 to 25 | yes |
| hop2-1 | 6 | ant 4, fil 1 | 30/30 | 0 | leptotene | circular, centric | 1 | 0 | no |
| hop2-1 | 6 | ant 5 fil 1 | 0/30 | 0 | G2 | circular , centric | 1 | 0 | no |
| hop2-1 | 6 | fil2 | 0/30 | 0 | G2 | circular , centric | 1 | 0 | no |
| hop2-1 | 6 | **bud 3**, ant1, fil1 | 0/30 | 0 | m-l P | circular,peripheral | 3 | 100 | no |
| hop2-1 | 6 | fil 2 | 0/30 | 0 | m-l P | circular,peripheral | 3 | 100 | no |
| hop2-1 | 6 | ant 2, fil 1 | 0/30 | 1 | eP | circular,peripheral | 1 | 100 | no |
| hop2-1 | 6 | fil2 | 0/30 | 1 | eP | circular,peripheral | 1 | 100 | no |
| hop2-1 | 6 | **bud 4**, ant1, fil1 | 0/30 | 1 | eP | circular,peripheral | 1 | 100 | no |
| hop2-1 | 6 | fil 2 | 0/30 | 1 | eP | circular,peripheral | 1 | 100 | no |
| hop2-1 | 6 | fil 3 | 0/30 | 1 | eP | circular,peripheral | 1 | 100 | no |
| hop2-1 | 8 | **bud 1**, ant1, fil1 | 0/30 | 0 | diakenesis | not visible | N/A | 100 | no |
| hop2-1 | 8 | fil 2 | 0/30 | 0 | diakenesis | not visible | N/A | 100 | no |
| hop2-1 | 8 | ant 2, fil 1 | 0/30 | 0 | diakenesis | not visible | N/A | 100 | no |
| hop2-1 | 8 | fil2 | 0/30 | 0 | diakenesis | not visible | N/A | 100 | no |
| hop2-1 | 8 | fil 3 | 0/30 | 0 | anaphase I | not visible | N/A | 100 | no |
| hop2-1 | 8 | fil 4 | 0/30 | 0 | anaphase I | not visible | N/A | 100 | no |
| hop2-1 | 8 | ant 3, fil 1 | 0/30 | 0 | diakenesis | not visible | N/A | 100 | no |
| hop2-1 | 8 | ant 4, fil 1 | 0/30 | 2 | zygotene | not visible | 1 | 26-75 | no |
| hop2-1 | 8 | fil 2 | 0/30 | 2 | zygotene | not visible | 1 | 26-75 | no |
| hop2-1 | 8 | **bud 2**, ant1, fil1 | 0/30 | 0 | leptotene | circular, peripheral | 1 | 0 | no |
| hop2-1 | 8 | fil 2 | 0/30 | 0 | leptotene | circular, peripheral | 1 | 0 | no |
| hop2-1 | 8 | fil 3 | 0/30 | 0 | leptotene | circular, peripheral | 1 | 0 | no |
| hop2-1 | 8 | fil 4 | 0/30 | 0 | leptotene | circular, peripheral | 1 | 0 | no |
| hop2-1 | 8 | **bud 3**, ant1, fil1 | 0/30 | 2 | zygotene | not visible | 1 | 26-75 | no |
| hop2-1 | 8 | fil 2 | 0/30 | 2 | zygotene | not visible | 1 | 26-75 | no |

| | | | | | | | | | |
|---|---|---|---|---|---|---|---|---|---|
| hop2-1 | 8 | **bud 4**, ant1, fil1 | 30/30 | 0 | leptotene | circular, pericentric | 1 | 0 | no |
| hop2-1 | 8 | fil 2 | 30/30 | 0 | leptotene | circular, pericentric | 1 | 0 | no |
| hop2-1 | 8 | fil 3 | 30/30 | 0 | leptotene | circular, pericentric | 1 | 0 | no |
| hop2-1 | 8 | fil 4 | 30/30 | 0 | leptotene | circular, pericentric | 1 | 0 | no |
| hop2-1 | 10 | **bud 1**, ant1, fil1 | 30/30 | 0 | leptotene | circular , peripheral | 1 | 0 | no |
| hop2-1 | 10 | fil 2 | 30/30 | 0 | leptotene | circular , peripheral | 1 | 0 | no |
| hop2-1 | 10 | fil 3 | 30/30 | 0 | leptotene | circular , peripheral | 1 | 0 | no |
| hop2-1 | 10 | fil 4 | 30/30 | 0 | leptotene | circular , peripheral | 1 | 0 | no |
| hop2-1 | 10 | ant 2, fil 1 | 30/30 | 0 | leptotene | circular , peripheral | 1 | 0 | no |
| hop2-1 | 10 | fil2 | 30/30 | 0 | leptotene | circular , peripheral | 1 | 0 | no |
| hop2-1 | 10 | ant 3, fil 1 | 30/30 | 0 | leptotene | circular , peripheral | 1 | 0 | no |
| hop2-1 | 10 | fil2 | 30/30 | 0 | leptotene | circular , peripheral | 1 | 0 | no |
| hop2-1 | 10 | fil 3 | 30/30 | 0 | leptotene | circular , peripheral | 1 | 0 | no |
| hop2-1 | 10 | **bud 2**, ant1, fil1 | 0/30 | 0 | diakenesis | not visible | N/A | 100 | yes |
| hop2-1 | 10 | fil 2 | 0/30 | 0 | diakenesis | not visible | N/A | 100 | yes |
| hop2-1 | 10 | ant 2, fil 1 | 0/30 | 0 | diakenesis | not visible | N/A | 100 | yes |
| hop2-1 | 10 | fil2 | 0/30 | 0 | diakenesis | not visible | N/A | 100 | yes |
| hop2-1 | 10 | fil 3 | 0/30 | 0 | diakenesis | not visible | N/A | 100 | yes |
| hop2-1 | 10 | ant 3, fil 1 | 0/30 | 0 | m-l P | circular,peripheral | 3 | 100 | yes |
| hop2-1 | 10 | fil2 | 0/30 | 0 | m-l P | circular,peripheral | 3 | 100 | yes |
| hop2-1 | 10 | **bud 3**, ant1, fil1 | 0/30 | 1 | zygotene | not visible | 1 | 76-99 | yes |
| hop2-1 | 10 | fil 2 | 0/30 | 1 | zygotene | not visible | 1 | 76-99 | yes |
| **hop2-1** | **10** | **ant 2, fil 1** | **5/30** | **2** | **zygotene** | **not visible** | **1** | **1 to 25** | **yes** |
| **hop2-1** | **10** | **fil2** | **10/30** | **1** | **zygotene** | **not visible** | **1** | **1 to 25** | **yes** |
| **hop2-1** | **10** | **fil 3** | **30/30** | **2** | **zygotene** | **not visible** | **1** | **1 to 25** | **yes** |
| **hop2-1** | **10** | **fil 4** | **30/30** | **1** | **zygotene** | **not visible** | **1** | **1 to 25** | **yes** |
| hop2-1 | 10 | **bud 4**, ant1, fil1 | 0/30 | 0 | tetrad | not visible | N/A | 100 | no |
| hop2-1 | 10 | fil 2 | 0/30 | 0 | tetrad | not visible | N/A | 100 | no |

| | | | | | | | | | |
|---|---|---|---|---|---|---|---|---|---|
| hop2-1 | 12 | **bud 1**, ant1, fil1 | 30/30 | 2 | zygotene | not visible | 1 | 1 to 25 | yes |
| hop2-1 | 12 | fil 2 | 30/30 | 0 | leptotene | circular , peripheral | 1 | 0 | no |
| hop2-1 | 12 | fil 3 | 30/30 | 2 | zygotene | not visible | 1 | 1 to 25 | yes |
| hop2-1 | 12 | fil 4 | 30/30 | 0 | leptotene | circular , peripheral | 1 | 0 | no |
| hop2-1 | 12 | **bud 2**, ant1, fil1 | 10/30 | 0 | G2 | circular , centric | 1 | 0 | no |
| hop2-1 | 12 | fil 2 | 10/30 | 0 | G2 | circular , centric | 1 | 0 | no |
| hop2-1 | 12 | fil 3 | 10/30 | 0 | G2 | circular , centric | 1 | 0 | no |
| hop2-1 | 12 | **bud 3**, ant1, fil1 | 10/30 | 0 | leptotene | circular , peripheral | 1 | 0 | no |
| hop2-1 | 12 | fil 2 | 10/30 | 0 | leptotene | circular , peripheral | 1 | 0 | no |
| hop2-1 | 12 | fil 3 | 10/30 | 0 | leptotene | circular , peripheral | 1 | 0 | no |
| hop2-1 | 12 | fil 4 | 5/30 | 0 | leptotene | circular , peripheral | 1 | 0 | no |
| hop2-1 | 12 | **bud 4**, ant1, fil1 | 0/30 | 0 | m-l P | circular,peripheral | 2 | 100 | no |
| hop2-1 | 12 | fil 2 | 0/30 | 0 | m-l P | circular,peripheral | 2 | 100 | no |
| hop2-1 | 12 | **bud 5**, ant1, fil1 | 30/30 | 1 | zygotene | not visible | 1 | 1 to 25 | yes |
| hop2-1 | 12 | fil 2 | 30/30 | 1 | zygotene | not visible | 1 | 1 to 25 | yes |
| hop2-1 | 12 | ant 2, fil 1 | 30/30 | 0 | leptotene | circular , peripheral | 1 | 0 | no |
| hop2-1 | 12 | fil2 | 30/30 | 0 | leptotene | circular , peripheral | 1 | 0 | no |
| hop2-1 | 14 | **bud 1**, ant1, fil1 | 10/30 | 0 | leptotene | circular , peripheral | 1 | 0 | no |
| hop2-1 | 14 | fil 2 | 10/30 | 0 | leptotene | circular , peripheral | 1 | 0 | no |
| hop2-1 | 14 | fil 3 | 5/30 | 0 | leptotene | circular , peripheral | 1 | 0 | no |
| hop2-1 | 14 | ant 2, fil 1 | 30/30 | 0 | leptotene | circular, pericentric | 1 | 0 | no |
| hop2-1 | 14 | fil2 | 30/30 | 0 | leptotene | circular, pericentric | 1 | 0 | no |
| hop2-1 | 14 | fil 3 | 30/30 | 0 | leptotene | circular, pericentric | 1 | 0 | no |
| hop2-1 | 14 | **bud 2**, ant1, fil1 | 0/30 | 0 | m-l P | circular,peripheral | 2 | 100 | no |
| hop2-1 | 14 | fil 2 | 0/30 | 0 | m-l P | circular,peripheral | 2 | 100 | no |
| hop2-1 | 14 | **bud 3**, ant1, fil1 | 0/30 | 0 | metaphase I | not visible | N/A | 100 | no |
| hop2-1 | 14 | fil 2 | 0/30 | 0 | metaphase I | not visible | N/A | 100 | no |
| hop2-1 | 14 | **bud 4**, ant1, fil1 | 0/30 | 0 | diakenesis | not visible | N/A | 100 | no |

| | | | | | | | | | |
|---|---|---|---|---|---|---|---|---|---|
| hop2-1 | 14 | fil 2 | 0/30 | 0 | diakenesis | not visible | N/A | 100 | no |
| hop2-1 | 14 | fil 3 | 0/30 | 0 | diakenesis | not visible | N/A | 100 | no |
| hop2-1 | 14 | ant 2, fil 1 | 0/30 | 0 | diakenesis | not visible | N/A | 100 | no |
| hop2-1 | 14 | fil2 | 0/30 | 0 | diakenesis | not visible | N/A | 100 | no |
| hop2-1 | 14 | fil 3 | 0/30 | 0 | diakenesis | not visible | N/A | 100 | no |
| hop2-1 | 14 | **bud 5**, ant1, fil1 | 30/30 | 0 | leptotene | circular, pericentric | 1 | 0 | no |
| hop2-1 | 14 | fil 2 | 30/30 | 0 | leptotene | circular, pericentric | 1 | 0 | no |
| hop2-1 | 14 | fil 3 | 30/30 | 0 | leptotene | circular, pericentric | 1 | 0 | no |
| hop2-1 | 14 | **bud 6**, ant1, fil1 | 0/30 | 0 | anaphase I | not visible | N/A | 100 | no |
| hop2-1 | 14 | fil 2 | 0/30 | 0 | anaphase I | not visible | N/A | 100 | no |
| hop2-1 | 16 | **bud 1**, ant1, fil1 | 30/30 | 1 | zygotene | not visible | 1 | 1 to 25 | yes |
| hop2-1 | 16 | fil 2 | 30/30 | 1 | zygotene | not visible | 1 | 1 to 25 | yes |
| hop2-1 | 16 | fil 3 | 30/30 | 1 | zygotene | not visible | 1 | 1 to 25 | yes |
| hop2-1 | 16 | ant 2, fil 1 | 0/30 | 1 | eP | circular,peripheral | 1 | 100 | yes |
| hop2-1 | 16 | fil2 | 0/30 | 1 | eP | circular,peripheral | 1 | 100 | yes |
| hop2-1 | 16 | ant 3, fil 1 | 30/30 | 2 | zygotene | not visible | 1 | 26-75 | yes |
| hop2-1 | 16 | fil2 | 30/30 | 2 | zygotene | not visible | 1 | 26-75 | yes |
| hop2-1 | 16 | fil 3 | 30/30 | 2 | zygotene | not visible | 1 | 26-75 | yes |
| hop2-1 | 16 | ant 4, fil 1 | 30/30 | 1 | zygotene | not visible | 1 | 1 to 25 | yes |
| hop2-1 | 16 | fil2 | 30/30 | 1 | zygotene | not visible | 1 | 1 to 25 | yes |
| hop2-1 | 16 | fil 3 | 30/30 | 1 | zygotene | not visible | 1 | 1 to 25 | yes |
| hop2-1 | 16 | **bud 2**, ant1, fil1 | 10/30 | 0 | leptotene | circular, pericentric | 1 | 0 | no |
| hop2-1 | 16 | fil 2 | 15/30 | 0 | leptotene | circular, pericentric | 1 | 0 | no |
| hop2-1 | 16 | fil 3 | 30/30 | 0 | leptotene | circular, pericentric | 1 | 0 | no |
| hop2-1 | 16 | ant 2, fil 1 | 0/30 | 0 | m-l P | circular,peripheral | 2 | 100 | no |
| hop2-1 | 16 | fil2 | 0/30 | 0 | m-l P | circular,peripheral | 2 | 100 | no |
| hop2-1 | 16 | **bud 3**, ant1, fil1 | 0/30 | 0 | anaphase I | not visible | N/A | 100 | no |
| hop2-1 | 16 | fil 2 | 0/30 | 0 | anaphase I | not visible | N/A | 100 | no |

| | | | | | | | | | |
|---|---|---|---|---|---|---|---|---|---|
| hop2-1 | 16 | **bud 4**, ant1, fil1 | 0/30 | 1 | eP | circular,peripheral | 1 | 100 | yes |
| hop2-1 | 16 | fil 2 | 0/30 | 1 | eP | circular,peripheral | 1 | 100 | yes |
| hop2-1 | 16 | ant 2, fil 1 | 0/30 | 1 | zygotene | not visible | 1 | 76-99 | yes |
| hop2-1 | 16 | fil2 | 0/30 | 1 | zygotene | not visible | 1 | 76-99 | yes |
| hop2-1 | 16 | ant 3, fil 1 | 30/30 | 0 | leptotene | circular , peripheral | 1 | 0 | no |
| hop2-1 | 16 | fil2 | 30/30 | 0 | leptotene | circular , peripheral | 1 | 0 | no |
| hop2-1 | 18 | **bud 1**, ant1, fil1 | 0/30 | 1 | eP | circular,peripheral | 1 | 100 | yes |
| hop2-1 | 18 | fil 2 | 0/30 | 1 | eP | circular,peripheral | 1 | 100 | yes |
| hop2-1 | 18 | fil 3 | 0/30 | 1 | eP | circular,peripheral | 1 | 100 | yes |
| hop2-1 | 18 | ant 2, fil 1 | 0/30 | 1 | eP | circular,peripheral | 1 | 100 | yes |
| hop2-1 | 18 | fil2 | 0/30 | 1 | eP | circular,peripheral | 1 | 100 | yes |
| hop2-1 | 18 | **bud 2**, ant1, fil1 | 0/30 | 0 | metaphase I | not visible | N/A | 100 | no |
| hop2-1 | 18 | ant 2, fil 1 | 30/30 | 1 | zygotene | not visible | 1 | 76-99 | yes |
| hop2-1 | 18 | fil2 | 30/30 | 1 | zygotene | not visible | 1 | 76-99 | yes |
| hop2-1 | 18 | ant 3, fil 1 | 30/30 | 1 | zygotene | not visible | 1 | 76-99 | yes |
| hop2-1 | 18 | fil2 | 30/30 | 1 | zygotene | not visible | 1 | 76-99 | yes |
| hop2-1 | 18 | ant 4, fil 1 | 0/30 | 1 | eP | circular,peripheral | 1 | 100 | yes |
| hop2-1 | 18 | fil2 | 0/30 | 1 | eP | circular,peripheral | 1 | 100 | yes |
| hop2-1 | 18 | **bud 3**, ant1, fil1 | 30/30 | 2 | zygotene | not visible | 1 | 26-75 | yes |
| hop2-1 | 18 | fil 2 | 30/30 | 2 | zygotene | not visible | 1 | 26-75 | yes |
| hop2-1 | 18 | fil 3 | 30/30 | 2 | zygotene | not visible | 1 | 26-75 | yes |
| hop2-1 | 20 | **bud 1**, ant1, fil1 | 30/30 | 1 | zygotene | not visible | 1 | 76-99 | yes |
| hop2-1 | 20 | fil 2 | 30/30 | 1 | zygotene | not visible | 1 | 76-99 | yes |
| hop2-1 | 20 | fil 3 | 30/30 | 1 | zygotene | not visible | 1 | 76-99 | yes |
| hop2-1 | 20 | fil4 | 30/30 | 1 | zygotene | not visible | 1 | 76-99 | yes |
| hop2-1 | 20 | **bud 2**, ant1, fil1 | 30/30 | 2 | zygotene | not visible | 1 | 76-99 | yes |
| hop2-1 | 20 | fil 2 | 30/30 | 2 | zygotene | not visible | 1 | 76-99 | yes |
| hop2-1 | 20 | fil 3 | 30/30 | 2 | zygotene | not visible | 1 | 76-99 | yes |

| | | | | | | | | | |
|---|---|---|---|---|---|---|---|---|---|
| hop2-1 | 20 | ant 2, fil 1 | 30/30 | 2 | zygotene | not visible | 1 | 76-99 | yes |
| hop2-1 | 20 | fil2 | 30/30 | 2 | zygotene | not visible | 1 | 76-99 | yes |
| hop2-1 | 20 | fil 3 | 30/30 | 2 | zygotene | not visible | 1 | 76-99 | yes |
| hop2-1 | 20 | fil 4 | 30/30 | 2 | zygotene | not visible | 1 | 76-99 | yes |
| hop2-1 | 20 | ant 3, fil 1 | 30/30 | 2 | zygotene | not visible | 1 | 76-99 | yes |
| hop2-1 | 20 | fil2 | 30/30 | 2 | zygotene | not visible | 1 | 76-99 | yes |
| hop2-1 | 20 | fil 3 | 30/30 | 2 | zygotene | not visible | 1 | 76-99 | yes |
| hop2-1 | 20 | fil 4 | 30/30 | 2 | zygotene | not visible | 1 | 76-99 | yes |
| hop2-1 | 20 | ant 4, fil 1 | 30/30 | 2 | zygotene | not visible | 1 | 76-99 | yes |
| hop2-1 | 20 | fil2 | 30/30 | 2 | zygotene | not visible | 1 | 76-99 | yes |
| hop2-1 | 20 | fil 3 | 30/30 | 2 | zygotene | not visible | 1 | 76-99 | yes |
| hop2-1 | 20 | fil 4 | 30/30 | 2 | zygotene | not visible | 1 | 76-99 | yes |
| hop2-1 | 20 | **bud 3**, ant1, fil1 | 0/30 | 0 | G2 | circular , centric | 1 | 0 | no |
| hop2-1 | 20 | fil 2 | 0/30 | 0 | G2 | circular , centric | 1 | 0 | no |
| hop2-1 | 20 | **bud 4**, ant1, fil1 | 30/30 | 0 | eP | circular,peripheral | 1 | 100 | no |
| hop2-1 | 20 | fil 2 | 30/30 | 0 | eP | circular,peripheral | 1 | 100 | no |
| hop2-1 | 20 | ant 2, fil 1 | 0/30 | 0 | m-l P | circular,peripheral | 3 | 100 | yes |
| hop2-1 | 20 | fil2 | 0/30 | 0 | m-l P | circular,peripheral | 3 | 100 | yes |
| hop2-1 | 20 | **bud 5**, ant1, fil1 | 30/30 | 2 | zygotene | not visible | 1 | 26-75 | yes |
| hop2-1 | 20 | fil 2 | 30/30 | 2 | zygotene | not visible | 1 | 26-75 | yes |
| hop2-1 | 20 | fil 3 | 30/30 | 2 | zygotene | not visible | 1 | 26-75 | yes |
| hop2-1 | 20 | fil4 | 30/30 | 2 | zygotene | not visible | 1 | 26-75 | yes |
| hop2-1 | 22 | **bud 1**, ant1, fil1 | 30/30 | 0 | leptotene | circular , centric | 1 | 0 | yes |
| hop2-1 | 22 | ant 2, fil 1 | 30/30 | 0 | leptotene | circular , centric | 1 | 0 | yes |
| hop2-1 | 22 | fil2 | 30/30 | 0 | leptotene | circular , centric | 1 | 0 | yes |
| hop2-1 | 22 | fil 3 | 30/30 | 0 | leptotene | circular , centric | 1 | 0 | yes |
| hop2-1 | 22 | fil 4 | 30/30 | 0 | leptotene | circular , centric | 1 | 0 | yes |
| hop2-1 | 22 | **bud 2**, ant1, fil1 | 30/30 | 0 | leptotene | circular, pericentric | 1 | 0 | no |

| | | | | | | | | | |
|---|---|---|---|---|---|---|---|---|---|
| hop2-1 | 22 | fil 2 | 30/30 | 0 | leptotene | circular, pericentric | 1 | 0 | no |
| hop2-1 | 22 | fil 3 | 30/30 | 0 | leptotene | circular, pericentric | 1 | 0 | no |
| hop2-1 | 22 | ant 2, fil 1 | 30/30 | 2 | zygotene | not visible | 1 | 26-75 | yes |
| hop2-1 | 22 | fil2 | 30/30 | 2 | zygotene | not visible | 1 | 26-75 | yes |
| hop2-1 | 22 | fil 3 | 30/30 | 2 | zygotene | not visible | 1 | 26-75 | yes |
| hop2-1 | 22 | ant 3, fil 1 | 30/30 | 2 | zygotene | not visible | 1 | 26-75 | yes |
| hop2-1 | 22 | fil2 | 30/30 | 2 | zygotene | not visible | 1 | 26-75 | yes |
| hop2-1 | 22 | fil 3 | 30/30 | 2 | zygotene | not visible | 1 | 26-75 | yes |
| hop2-1 | 22 | ant 4, fil 1 | 30/30 | 2 | zygotene | not visible | 1 | 26-75 | yes |
| hop2-1 | 22 | fil2 | 30/30 | 2 | zygotene | not visible | 1 | 26-75 | yes |
| hop2-1 | 22 | fil 3 | 30/30 | 2 | zygotene | not visible | 1 | 26-75 | yes |
| hop2-1 | 22 | ant 5, fil 1 | 30/30 | 2 | zygotene | not visible | 1 | 1 to 25 | yes |
| hop2-1 | 22 | fil2 | 30/30 | 2 | zygotene | not visible | 1 | 1 to 25 | yes |
| hop2-1 | 22 | fil 3 | 30/30 | 2 | zygotene | not visible | 1 | 1 to 25 | yes |
| hop2-1 | 22 | **bud 3**, ant1, fil1 | 30/30 | 1 | zygotene | not visible | 1 | 76-99 | yes |
| hop2-1 | 22 | fil 2 | 30/30 | 1 | zygotene | not visible | 1 | 76-99 | yes |
| hop2-1 | 22 | ant 2, fil 1 | 30/30 | 1 | zygotene | not visible | 1 | 1 to 25 | yes |
| hop2-1 | 22 | fil2 | 30/30 | 1 | zygotene | not visible | 1 | 1 to 25 | yes |
| hop2-1 | 22 | ant 3, fil 1 | 30/30 | 2 | zygotene | not visible | 1 | 1 to 25 | yes |
| hop2-1 | 22 | fil2 | 30/30 | 2 | zygotene | not visible | 1 | 1 to 25 | yes |
| hop2-1 | 22 | fil 3 | 30/30 | 2 | zygotene | not visible | 1 | 1 to 25 | yes |
| hop2-1 | 22 | fil 4 | 30/30 | 2 | zygotene | not visible | 1 | 1 to 25 | yes |
| hop2-1 | 22 | ant 4, fil 1 | 30/30 | 2 | zygotene | not visible | 1 | 1 to 25 | yes |
| hop2-1 | 22 | fil2 | 30/30 | 2 | zygotene | not visible | 1 | 1 to 25 | yes |
| hop2-1 | 22 | fil 3 | 30/30 | 2 | zygotene | not visible | 1 | 1 to 25 | yes |
| hop2-1 | 22 | fil 4 | 30/30 | 2 | zygotene | not visible | 1 | 1 to 25 | yes |
| hop2-1 | 22 | ant 5, fil 1 | 30/30 | 2 | zygotene | not visible | 1 | 26-75 | yes |
| hop2-1 | 22 | fil2 | 30/30 | 2 | zygotene | not visible | 1 | 26-75 | yes |
| hop2-1 | 22 | ant 6, fil 1 | 30/30 | 1 | zygotene | not visible | 1 | 76-99 | yes |

| | | | | | | | | | |
|---|---|---|---|---|---|---|---|---|---|
| hop2-1 | 22 | fil2 | 30/30 | 1 | zygotene | not visible | 1 | 76-99 | yes |
| hop2-1 | 22 | **bud 4**, ant1, fil1 | 30/30 | 2 | zygotene | not visible | 1 | 1 to 25 | yes |
| hop2-1 | 22 | fil 2 | 30/30 | 1 | zygotene | not visible | 1 | 1 to 25 | yes |
| hop2-1 | 22 | fil 3 | 30/30 | 2 | zygotene | not visible | 1 | 1 to 25 | yes |
| hop2-1 | 22 | ant 2, fil 1 | 30/30 | 0 | leptotene | circular, pericentric | 1 | 0 | no |
| hop2-1 | 22 | fil2 | 30/30 | 0 | leptotene | circular, pericentric | 1 | 0 | no |
| hop2-1 | 22 | ant 3, fil 1 | 30/30 | 2 | zygotene | not visible | 1 | 1 to 25 | yes |
| hop2-1 | 22 | fil2 | 30/30 | 2 | zygotene | not visible | 1 | 1 to 25 | yes |
| hop2-1 | 22 | fil 3 | 30/30 | 2 | zygotene | not visible | 1 | 1 to 25 | yes |
| hop2-1 | 22 | ant 4, fil 1 | 30/30 | 1 | zygotene | not visible | 1 | 76-99 | yes |
| hop2-1 | 22 | fil2 | 30/30 | 1 | zygotene | not visible | 1 | 76-99 | yes |
| hop2-1 | 22 | fil 3 | 30/30 | 1 | zygotene | not visible | 1 | 76-99 | yes |
| hop2-1 | 24 | **bud 1**, ant1, fil1 | 30/30 | 2 | zygotene | not visible | 1 | 1 to 25 | yes |
| hop2-1 | 24 | fil 2 | 30/30 | 1 | zygotene | not visible | 1 | 1 to 25 | yes |
| hop2-1 | 24 | fil 3 | 30/30 | 1 | zygotene | not visible | 1 | 1 to 25 | yes |
| hop2-1 | 24 | fil4 | 30/30 | 1 | zygotene | not visible | 1 | 1 to 25 | yes |
| hop2-1 | 24 | ant 2, fil 1 | 30/30 | 0 | eP | circular,peripheral | 1 | 100 | yes |
| hop2-1 | 24 | fil2 | 30/30 | 0 | eP | circular,peripheral | 1 | 100 | yes |
| hop2-1 | 24 | **bud 2**, ant1, fil1 | 0/30 | 0 | m-l P | circular,peripheral | 2 | 100 | no |
| hop2-1 | 24 | fil 2 | 0/30 | 0 | m-l P | circular,peripheral | 2 | 100 | no |
| hop2-1 | 24 | ant 2, fil 1 | 0/30 | 0 | m-l P | circular,peripheral | 3 | 100 | yes |
| hop2-1 | 24 | fil2 | 0/30 | 0 | m-l P | circular,peripheral | 3 | 100 | yes |
| hop2-1 | 24 | fil 3 | 30/30 | 0 | m-l P | circular,peripheral | 2 | 100 | no |
| hop2-1 | 24 | fil 4 | 30/30 | 0 | m-l P | circular,peripheral | 2 | 100 | no |
| hop2-1 | 24 | ant 3 fil 1 | 0/30 | 0 | diplotene | not visible | N/A | 100 | no |
| hop2-1 | 24 | fil2 | 0/30 | 0 | diplotene | not visible | N/A | 100 | no |
| hop2-1 | 24 | fil 3 | 0/30 | 0 | diplotene | not visible | N/A | 100 | no |
| hop2-1 | 24 | fil 4 | 0/30 | 0 | diplotene | not visible | N/A | 100 | no |

| | | | | | | | | | |
|---|---|---|---|---|---|---|---|---|---|
| hop2-1 | 26 | **bud 1**, ant1, fil1 | 30/30 | 1 | zygotene | not visible | 1 | 1 to 25 | yes |
| hop2-1 | 26 | fil 2 | 30/30 | 1 | zygotene | not visible | 1 | 1 to 25 | yes |
| hop2-1 | 26 | fil 3 | 30/30 | 0 | leptotene | circular , peripheral | 1 | 0 | yes |
| hop2-1 | 26 | fil4 | 30/30 | 0 | leptotene | circular , peripheral | 1 | 0 | yes |
| hop2-1 | 26 | ant 2, fil 1 | 30/30 | 2 | zygotene | not visible | 1 | 1 to 25 | yes |
| hop2-1 | 26 | fil2 | 30/30 | 1 | zygotene | not visible | 1 | 1 to 25 | yes |
| hop2-1 | 26 | fil 3 | 30/30 | 1 | zygotene | not visible | 1 | 1 to 25 | yes |
| hop2-1 | 26 | fil 4 | 30/30 | 1 | zygotene | not visible | 1 | 1 to 25 | yes |
| hop2-1 | 26 | ant 3, fil 1 | 30/30 | 0 | leptotene | circular , peripheral | 1 | 0 | yes |
| hop2-1 | 26 | fil2 | 30/30 | 0 | leptotene | circular , peripheral | 1 | 0 | yes |
| hop2-1 | 26 | ant 4, fil 1 | 30/30 | 1 | zygotene | not visible | 1 | 1 to 25 | yes |
| hop2-1 | 26 | fil2 | 30/30 | 0 | leptotene | circular , peripheral | 1 | 0 | yes |
| hop2-1 | 26 | **bud 2**, ant1, fil1 | 30/30 | 2 | zygotene | not visible | 1 | 26-75 | yes |
| hop2-1 | 26 | fil 2 | 30/30 | 2 | zygotene | not visible | 1 | 26-75 | yes |
| hop2-1 | 26 | **bud 3**, ant1, fil1 | 30/30 | 0 | leptotene | circular , peripheral | 1 | 0 | yes |
| hop2-1 | 26 | fil 2 | 30/30 | 0 | leptotene | circular , peripheral | 1 | 0 | yes |
| hop2-1 | 26 | ant 2, fil 1 | 30/30 | 0 | m-l P | circular,peripheral | 2 | 100 | yes |
| hop2-1 | 26 | fil2 | 30/30 | 0 | m-l P | circular,peripheral | 2 | 100 | yes |
| hop2-1 | 26 | **bud 4**, ant1, fil1 | 30/30 | 2 | zygotene | not visible | 1 | 1 to 25 | yes |
| hop2-1 | 26 | fil 2 | 30/30 | 0 | leptotene | circular, pericentric | 1 | 0 | no |
| hop2-1 | 26 | fil 3 | 30/30 | 0 | leptotene | circular , peripheral | 1 | 0 | yes |
| hop2-1 | 26 | fil4 | 30/30 | 0 | leptotene | circular , peripheral | 1 | 0 | yes |
| hop2-1 | 26 | ant 2, fil 1 | 30/30 | 0 | leptotene | circular, pericentric | 1 | 0 | no |
| hop2-1 | 26 | fil2 | 30/30 | 0 | leptotene | circular, pericentric | 1 | 0 | no |
| hop2-1 | 26 | fil 3 | 30/30 | 0 | leptotene | circular , peripheral | 1 | 0 | yes |
| hop2-1 | 26 | fil 4 | 30/30 | 0 | leptotene | circular , peripheral | 1 | 0 | yes |
| hop2-1 | 26 | ant 3, fil 1 | 30/30 | 0 | leptotene | circular , peripheral | 1 | 0 | yes |
| hop2-1 | 26 | fil2 | 30/30 | 0 | leptotene | circular , peripheral | 1 | 0 | yes |

| | | | | | | | | | |
|---|---|---|---|---|---|---|---|---|---|
| hop2-1 | 26 | fil 3 | 30/30 | 0 | leptotene | circular , peripheral | 1 | 0 | yes |
| hop2-1 | 28 | **bud 1**, ant1, fil1 | 30/30 | 0 | leptotene | circular, pericentric | 1 | 0 | no |
| hop2-1 | 28 | fil 2 | 30/30 | 0 | leptotene | circular, pericentric | 1 | 0 | no |
| hop2-1 | 28 | fil 3 | 30/30 | 0 | leptotene | circular, pericentric | 1 | 0 | no |
| hop2-1 | 28 | **bud 2**, ant1, fil1 | 0/30 | 0 | metaphase I | not visible | N/A | 100 | no |
| hop2-1 | 28 | fil 2 | 0/30 | 0 | metaphase I | not visible | N/A | 100 | no |
| hop2-1 | 28 | fil 3 | 0/30 | 0 | metaphase I | not visible | N/A | 100 | no |
| hop2-1 | 28 | ant 2, fil 1 | 30/30 | 2 | zygotene | not visible | 1 | 26-75 | yes |
| hop2-1 | 28 | fil2 | 30/30 | 1 | zygotene | not visible | 1 | 1 to 25 | yes |
| hop2-1 | 28 | fil 3 | 30/30 | 1 | zygotene | not visible | 1 | 1 to 25 | yes |
| hop2-1 | 28 | **bud 3**, ant1, fil1 | 30/30 | 1 | zygotene | not visible | 1 | 26-75 | yes |
| hop2-1 | 28 | fil 2 | 30/30 | 1 | zygotene | not visible | 1 | 1 to 25 | yes |
| hop2-1 | 28 | fil 3 | 30/30 | 1 | zygotene | not visible | 1 | 1 to 25 | yes |
| hop2-1 | 28 | **bud 4**, ant1, fil1 | 0/30 | 0 | diakenesis | not visible | N/A | 100 | no |
| hop2-1 | 28 | fil 2 | 0/30 | 0 | diakenesis | not visible | N/A | 100 | no |
| hop2-1 | 28 | fil 3 | 0/30 | 0 | diakenesis | not visible | N/A | 100 | no |
| hop2-1 | 28 | **bud 5**, ant1, fil1 | 0/30 | 0 | G2 | circular , centric | 1 | 0 | no |
| hop2-1 | 28 | fil 2 | 0/30 | 0 | G2 | circular , centric | 1 | 0 | no |
| hop2-1 | 28 | fil 3 | 0/30 | 0 | G2 | circular , centric | 1 | 0 | no |
| hop2-1 | 28 | **bud 6**, ant1, fil1 | 30/30 | 2 | zygotene | not visible | 1 | 26-75 | yes |
| hop2-1 | 28 | fil 2 | 30/30 | 2 | zygotene | not visible | 1 | 26-75 | yes |
| hop2-1 | 28 | fil 3 | 30/30 | 2 | zygotene | not visible | 1 | 26-75 | yes |
| hop2-1 | 30 | **bud 1**, ant1, fil1 | 30/30 | 0 | m-l P | circular,peripheral | 2 | 100 | yes |
| hop2-1 | 30 | fil 2 | 30/30 | 0 | m-l P | circular,peripheral | 2 | 100 | yes |
| hop2-1 | 30 | fil 3 | 30/30 | 0 | m-l P | circular,peripheral | 2 | 100 | yes |
| hop2-1 | 30 | ant 2, fil 1 | 30/30 | 0 | m-l P | circular,peripheral | 2 | 100 | yes |
| hop2-1 | 30 | fil2 | 30/30 | 0 | m-l P | circular,peripheral | 2 | 100 | yes |
| hop2-1 | 30 | fil 3 | 30/30 | 0 | m-l P | circular,peripheral | 2 | 100 | yes |

| | | | | | | | | | |
|---|---|---|---|---|---|---|---|---|---|
| hop2-1 | 30 | ant 3, fil 1 | 30/30 | 0 | m-l P | circular,peripheral | 2 | 100 | yes |
| hop2-1 | 30 | fil2 | 30/30 | 0 | m-l P | circular,peripheral | 2 | 100 | yes |
| hop2-1 | 30 | fil 3 | 30/30 | 0 | m-l P | circular,peripheral | 2 | 100 | yes |
| hop2-1 | 30 | ant 4, fil 1 | 30/30 | 1 | eP | circular,peripheral | 1 | 100 | yes |
| hop2-1 | 30 | fil2 | 30/30 | 1 | eP | circular,peripheral | 1 | 100 | yes |
| hop2-1 | 30 | fil 3 | 30/30 | 1 | eP | circular,peripheral | 1 | 100 | yes |
| hop2-1 | 30 | **bud 2**, ant1, fil1 | 30/30 | 1 | eP | circular,peripheral | 1 | 100 | yes |
| hop2-1 | 30 | fil 2 | 30/30 | 1 | eP | circular,peripheral | 1 | 100 | yes |
| hop2-1 | 30 | fil 3 | 30/30 | 1 | eP | circular,peripheral | 1 | 100 | yes |
| hop2-1 | 32 | **bud 1**, ant1, fil1 | 30/30 | 1 | zygotene | not visible | 1 | 76-99 | yes |
| hop2-1 | 32 | fil 2 | 30/30 | 1 | zygotene | not visible | 1 | 76-99 | yes |
| hop2-1 | 32 | fil 3 | 30/30 | 1 | zygotene | not visible | 1 | 76-99 | yes |
| hop2-1 | 32 | ant 2, fil 1 | 30/30 | 1 | zygotene | not visible | 1 | 1 to 25 | yes |
| hop2-1 | 32 | fil2 | 30/30 | 1 | zygotene | not visible | 1 | 1 to 25 | yes |
| hop2-1 | 32 | ant 3, fil 1 | 30/30 | 2 | zygotene | not visible | 1 | 26-75 | yes |
| hop2-1 | 32 | fil2 | 30/30 | 2 | zygotene | not visible | 1 | 26-75 | yes |
| hop2-1 | 32 | **bud 2**, ant1, fil1 | 30/30 | 1 | eP | circular,peripheral | 1 | 100 | yes |
| hop2-1 | 32 | fil 2 | 30/30 | 1 | eP | circular,peripheral | 1 | 100 | yes |
| hop2-1 | 32 | **bud 3**, ant1, fil1 | 30/30 | 1 | eP | circular,peripheral | 1 | 100 | yes |
| hop2-1 | 32 | fil 2 | 30/30 | 1 | eP | circular,peripheral | 1 | 100 | yes |
| hop2-1 | 32 | **bud 4**, ant1, fil1 | 30/30 | 2 | zygotene | not visible | 1 | 76-99 | yes |
| hop2-1 | 32 | fil 2 | 30/30 | 2 | zygotene | not visible | 1 | 76-99 | yes |
| hop2-1 | 32 | fil 3 | 30/30 | 2 | zygotene | not visible | 1 | 76-99 | yes |
| hop2-1 | 32 | ant 2, fil 1 | 30/30 | 1 | zygotene | not visible | 1 | 76-99 | yes |
| hop2-1 | 32 | fil2 | 30/30 | 2 | zygotene | not visible | 1 | 76-99 | yes |
| hop2-1 | 32 | fil 3 | 30/30 | 1 | zygotene | not visible | 1 | 76-99 | yes |
| hop2-1 | 32 | **bud 5**, ant1, fil1 | 30/30 | 1 | eP | circular,peripheral | 1 | 100 | yes |
| hop2-1 | 32 | ant 2, fil 1 | 30/30 | 2 | zygotene | not visible | 1 | 1 to 25 | yes |

| | | | | | | | | | |
|---|---|---|---|---|---|---|---|---|---|
| hop2-1 | 32 | fil2 | 30/30 | 2 | zygotene | not visible | 1 | 1 to 25 | yes |
| hop2-1 | 32 | **bud 6**, ant1, fil1 | 30/30 | 0 | diplotene | not visible | N/A | 100 | no |
| hop2-1 | 32 | fil 2 | 30/30 | 0 | diplotene | not visible | N/A | 100 | no |
| hop2-1 | 32 | fil 3 | 30/30 | 0 | diplotene | not visible | N/A | 100 | no |
| hop2-1 | 32 | fil4 | 30/30 | 0 | diplotene | not visible | N/A | 100 | no |
| hop2-1 | 32 | ant 2, fil 1 | 0/30 | 0 | diakenesis | not visible | N/A | 100 | no |
| hop2-1 | 32 | fil2 | 0/30 | 0 | diakenesis | not visible | N/A | 100 | no |
| hop2-1 | 32 | **bud 7**, ant1, fil1 | 30/30 | 2 | zygotene | not visible | 1 | 1 to 25 | yes |
| hop2-1 | 32 | fil 2 | 30/30 | 2 | zygotene | not visible | 1 | 1 to 25 | yes |
| hop2-1 | 32 | fil 3 | 30/30 | 2 | zygotene | not visible | 1 | 1 to 25 | yes |

### Col data begins here

| Geno-type | Time point (hr) | Bud, Anther, Filament | # meiocyte with EdU signal/total | Level γH2AX signal | Meiotic stage. | Nucleolus shape & location | Amount Callose | % bi-nucleate. tapetum. | Tapetum labeled w/EdU |
|---|---|---|---|---|---|---|---|---|---|
| **COL** | **0** | **bud 1, ant1, fil1** | **4/30** | **0** | **G2** | **circular , centric** | **1** | **0** | **no** |
| COL | 0 | fil 2 | 0/30 | 0 | G2 | circular , centric | 1 | 0 | no |
| COL | 0 | ant 2, fil 1 | 30/30 | 0 | G2 | circular , centric | 1 | 0 | no |
| COL | 0 | fil2 | 20/30 | 0 | G2 | circular , centric | 1 | 0 | no |
| COL | 0 | ant 3, fil 1 | 30/30 | 0 | G2 | circular , centric | 1 | 0 | no |
| COL | 0 | fil2 | 30/30 | 0 | G2 | circular , centric | 1 | 0 | no |
| COL | 0 | ant 4, fil 1 | 30/30 | 0 | G2 | circular , centric | 1 | 0 | no |
| COL | 0 | fil2 | 30/30 | 0 | G2 | circular , centric | 1 | 0 | no |
| COL | 0 | **bud 2**, ant1, fil1 | 0/30 | 1 | leptotene | circular , peripheral | 1 | 0 | yes |
| COL | 0 | fil 2 | 0/30 | 1 | leptotene | circular , peripheral | 1 | 0 | yes |

| | | | | | | | | | |
|---|---|---|---|---|---|---|---|---|---|
| COL | 0 | ant 2, fil 1 | 0/30 | 1 | leptotene | circular , peripheral | 1 | 0 | yes |
| COL | 0 | fil2 | 0/30 | 1 | leptotene | circular , peripheral | 1 | 0 | yes |
| COL | 0 | ant 3, fil 1 | 0/30 | 0 | leptotene | circular , pericentric | 1 | 0 | no |
| COL | 0 | fil2 | 0/30 | 0 | leptotene | circular , pericentric | 1 | 0 | no |
| COL | 0 | fil 3 | 0/30 | 0 | leptotene | circular , pericentric | 1 | 0 | no |
| COL | 0 | ant 4, fil 1 | 0/30 | 2 | zygotene | not visible | 1 | 1 to 25 | yes |
| COL | 0 | fil2 | 0/30 | 2 | zygotene | not visible | 1 | 1 to 25 | yes |
| COL | 0 | **bud 3**, ant1, fil1 | 0/30 | 0 | zygotene | not visible | 1 | 76-99 | no |
| COL | 0 | fil 2 | 0/30 | 1 | zygotene | not visible | 1 | 76-99 | no |
| COL | 0 | fil 3 | 0/30 | 1 | zygotene | not visible | 1 | 76-99 | no |
| COL | 0 | **bud 4**, ant1, fil1 | 0/30 | 1 | zygotene | not visible | 1 | 76-99 | no |
| COL | 0 | fil 2 | 0/30 | 1 | zygotene | not visible | 1 | 76-99 | no |
| COL | 0 | fil 3 | 0/30 | 1 | zygotene | not visible | 1 | 76-99 | no |
| COL | 2 | **bud 1**, ant1, fil1 | 0/30 | 0 | leptotene | circular pericentric | 1 | 0 | no |
| COL | 2 | fil 2 | 0/30 | 2 | zygotene | not visible | 1 | 1 to 25 | yes |
| COL | 2 | fil 3 | 0/30 | 2 | zygotene | not visible | 1 | 1 to 25 | yes |
| COL | 2 | ant 2, fil 1 | 20/30 | 0 | G2 | circular , centric | 1 | 0 | no |
| COL | 2 | ant 3, fil 1 | 0/30 | 2 | zygotene | not visible | 1 | 1 to 25 | yes |
| COL | 2 | fil2 | 0/30 | 2 | zygotene | not visible | 1 | 1 to 25 | yes |
| COL | 2 | fil 3 | 0/30 | 2 | zygotene | not visible | 1 | 1 to 25 | yes |
| COL | 2 | **bud 2**, ant1, fil1 | 0/30 | 0 | m-l P | circular,peripheral | 3 | 100 | yes |
| COL | 2 | fil 2 | 0/30 | 0 | m-l P | circular,peripheral | 3 | 100 | yes |
| COL | 2 | fil 3 | 0/30 | 0 | m-l P | circular,peripheral | 3 | 100 | yes |
| COL | 2 | ant 2, fil 1 | 0/30 | 2 | zygotene | not visible | 1 | 26-75 | yes |
| COL | 2 | fil2 | 0/30 | 2 | zygotene | not visible | 1 | 26-75 | yes |
| COL | 2 | fil 3 | 0/30 | 1 | zygotene | not visible | 1 | 76-99 | yes |
| COL | 2 | fil 4 | 0/30 | 1 | zygotene | not visible | 1 | 76-99 | yes |
| COL | 2 | **bud 3**, ant1, fil1 | 0/30 | 1 | zygotene | not visible | 1 | 76-99 | yes |

| | | | | | | | | | |
|---|---|---|---|---|---|---|---|---|---|
| COL | 2 | fil 2 | 0/30 | 1 | zygotene | not visible | 1 | 76-99 | yes |
| COL | 2 | fil 3 | 0/30 | 1 | zygotene | not visible | 1 | 76-99 | yes |
| COL | 2 | ant 2, fil 1 | 0/30 | 1 | zygotene | not visible | 1 | 76-99 | yes |
| COL | 2 | fil2 | 0/30 | 1 | zygotene | not visible | 1 | 76-99 | yes |
| COL | 2 | fil 3 | 0/30 | 1 | zygotene | not visible | 1 | 76-99 | yes |
| COL | 2 | fil 4 | 0/30 | 1 | zygotene | not visible | 1 | 76-99 | yes |
| COL | 2 | **bud 4** ant 3, fil 1 | 0/30 | 2 | zygotene | not visible | 1 | 1 to 25 | yes |
| COL | 2 | fil2 | 0/30 | 2 | zygotene | not visible | 1 | 1 to 25 | yes |
| COL | 2 | fil 3 | 0/30 | 2 | zygotene | not visible | 1 | 1 to 25 | yes |
| COL | 2 | fil 4 | 0/30 | 2 | zygotene | not visible | 1 | 1 to 25 | yes |
| COL | 2 | **bud 5**, ant1, fil1 | 0/30 | 2 | zygotene | not visible | 1 | 26-75 | no |
| COL | 2 | fil 2 | 0/30 | fbf | eP | circular,peripheral | 1 | 100 | no |
| COL | 2 | ant 2, fil 1 | 0/30 | 0 | leptotene | circular , centric | 1 | 0 | no |
| COL | 2 | fil2 | 0/30 | 0 | leptotene | circular , centric | 1 | 0 | no |
| COL | 2 | fil 3 | 0/30 | 0 | leptotene | circular , centric | 1 | 0 | no |
| COL | 2 | fil 4 | 0/30 | 0 | leptotene | circular , centric | 1 | 0 | no |
| COL | 2 | **bud 6**, ant1, fil1 | 0/30 | fbf | eP | circular,peripheral | 1 | 100 | no |
| COL | 2 | fil 2 | 0/30 | fbf | eP | circular,peripheral | 1 | 100 | no |
| COL | 2 | fil 3 | 0/30 | fbf | eP | circular,peripheral | 1 | 100 | no |
| COL | 2 | **bud 7**, ant1, fil1 | 0/30 | 2 | zygotene | not visible | 1 | 1 to 25 | yes |
| COL | 2 | ant 2, fil 1 | 5/30 | 0 | G2 | circular , centric | 1 | 0 | no |
| COL | 2 | fil2 | 0/30 | 0 | G2 | circular , centric | 1 | 0 | no |
| COL | 2 | fil 3 | 0/30 | 0 | G2 | circular , centric | 1 | 0 | no |
| COL | 2 | **bud 8**, ant1, fil1 | 0/30 | 2 | zygotene | not visible | 1 | 26-75 | no |
| COL | 2 | fil 2 | 0/30 | 2 | zygotene | not visible | 1 | 26-75 | no |
| COL | 2 | fil 3 | 0/30 | 2 | zygotene | not visible | 1 | 26-75 | no |
| COL | 2 | fil 4 | 0/30 | 2 | zygotene | not visible | 1 | 26-75 | no |
| COL | 4 | **bud 1**, ant1, fil1 | 0/30 | 0 | diplotene | not visible | N/A | 100 | no |

| | | | | | | | | | |
|---|---|---|---|---|---|---|---|---|---|
| COL | 4 | fil 2 | 0/30 | 0 | diplotene | not visible | N/A | 100 | no |
| COL | 4 | fil 3 | 0/30 | 0 | diplotene | not visible | N/A | 100 | no |
| COL | 4 | fil 4 | 0/30 | fbf | eP | circular,peripheral | 1 | 100 | no |
| COL | 4 | ant 2, fil 1 | 0/30 | 2 | zygotene | not visible | 1 | 26-75 | yes |
| COL | 4 | fil2 | 0/30 | 2 | zygotene | not visible | 1 | 26-75 | yes |
| COL | 4 | fil 3 | 0/30 | 2 | zygotene | not visible | 1 | 26-75 | yes |
| COL | 4 | ant 3, fil 1 | 0/30 | 2 | zygotene | not visible | 1 | 1 to 25 | yes |
| COL | 4 | fil2 | 0/30 | 2 | zygotene | not visible | 1 | 1 to 25 | yes |
| COL | 4 | fil 3 | 0/30 | 2 | zygotene | not visible | 1 | 1 to 25 | yes |
| COL | 4 | fil 4 | 0/30 | 2 | zygotene | not visible | 1 | 1 to 25 | yes |
| **COL** | **4** | **ant 4, fil 1** | **5/30** | **0** | **leptotene** | **circular , centric** | **1** | **0** | **no** |
| **COL** | **4** | **fil2** | **10/30** | **0** | **leptotene** | **circular , centric** | **1** | **0** | **no** |
| COL | 4 | **bud 2**, ant1, fil1 | 0/30 | 2 | zygotene | not visible | 1 | 1 to 25 | no |
| COL | 4 | fil 2 | 0/30 | 1 | zygotene | not visible | 1 | 76-99 | no |
| COL | 4 | ant 2, fil 1 | 0/30 | 0 | m-l P | circular,peripheral | 2 | 100 | no |
| COL | 4 | fil2 | 0/30 | 0 | m-l P | circular,peripheral | 2 | 100 | no |
| COL | 4 | fil 3 | 0/30 | 0 | m-l P | circular,peripheral | 2 | 100 | no |
| COL | 4 | ant 3, fil 1 | 0/30 | 2 | zygotene | not visible | 1 | 1 to 25 | yes |
| COL | 4 | fil2 | 0/30 | 2 | zygotene | not visible | 1 | 1 to 25 | yes |
| COL | 4 | fil 3 | 0/30 | 2 | zygotene | not visible | 1 | 1 to 25 | yes |
| COL | 4 | fil 4 | 0/30 | 2 | zygotene | not visible | 1 | 1 to 25 | yes |
| COL | 4 | **bud 3**, ant1, fil1 | 0/30 | 0 | m-l P | circular,peripheral | 3 | 100 | no |
| COL | 4 | fil 2 | 0/30 | 0 | m-l P | circular,peripheral | 3 | 100 | no |
| COL | 4 | fil 3 | 0/30 | 0 | m-l P | circular,peripheral | 3 | 100 | no |
| COL | 4 | ant 2, fil 1 | 0/30 | 0 | m-l P | circular,peripheral | 3 | 100 | no |
| COL | 4 | fil2 | 0/30 | 0 | m-l P | circular,peripheral | 3 | 100 | no |
| COL | 4 | fil 3 | 0/30 | 0 | m-l P | circular,peripheral | 3 | 100 | no |
| COL | 4 | fil 4 | 0/30 | fbf | eP | circular,peripheral | 1 | 100 | no |
| COL | 4 | **bud 4**, ant1, fil1 | 0/30 | 2 | zygotene | not visible | 1 | 1 to 25 | yes |

| | | | | | | | | | |
|---|---|---|---|---|---|---|---|---|---|
| COL | 4 | fil 2 | 0/30 | 2 | zygotene | not visible | 1 | 1 to 25 | yes |
| COL | 4 | fil 3 | 0/30 | 2 | zygotene | not visible | 1 | 1 to 25 | yes |
| COL | 4 | fil 4 | 0/30 | 2 | zygotene | not visible | 1 | 1 to 25 | yes |
| COL | 4 | ant 2, fil 1 | 0/30 | fbf | eP | circular,peripheral | 1 | 100 | no |
| COL | 4 | fil2 | 0/30 | fbf | eP | circular,peripheral | 1 | 100 | no |
| COL | 4 | fil 3 | 0/30 | fbf | eP | circular,peripheral | 1 | 100 | no |
| COL | 4 | fil 4 | 0/30 | fbf | eP | circular,peripheral | 1 | 100 | no |
| COL | 4 | **bud 5**, ant1, fil1 | 0/30 | 0 | m-l P | circular,peripheral | 3 | 100 | no |
| COL | 4 | fil 2 | 0/30 | 0 | m-l P | circular,peripheral | 3 | 100 | no |
| COL | 4 | fil 3 | 0/30 | 0 | m-l P | circular,peripheral | 3 | 100 | no |
| COL | 4 | ant 2, fil 1 | 0/30 | 0 | m-l P | circular,peripheral | 3 | 100 | yes |
| COL | 4 | fil2 | 0/30 | 0 | m-l P | circular,peripheral | 3 | 100 | yes |
| COL | 4 | fil 3 | 0/30 | 0 | m-l P | circular,peripheral | 3 | 100 | yes |
| COL | 4 | fil 4 | 0/30 | 0 | m-l P | circular,peripheral | 3 | 100 | yes |
| COL | 4 | **bud 6**, ant1, fil1 | 0/30 | 0 | m-l P | circular,peripheral | 3 | 100 | no |
| COL | 4 | fil 2 | 0/30 | 0 | m-l P | circular,peripheral | 2 | 100 | no |
| COL | 4 | fil 3 | 0/30 | 0 | m-l P | circular,peripheral | 2 | 100 | no |
| COL | 4 | ant 2, fil 1 | 0/30 | 0 | prometaphase I | not visible | N/A | 100 | no |
| COL | 4 | ant 3, fil 1 | 0/30 | 0 | m-l P | circular,peripheral | 2 | 100 | yes |
| COL | 4 | fil2 | 0/30 | 0 | m-l P | circular,peripheral | 2 | 100 | yes |
| COL | 4 | **bud 7**, ant1, fil1 | 0/30 | 0 | leptotene | circular , centric | 1 | 0 | no |
| COL | 4 | fil 2 | 0/30 | 0 | leptotene | circular , centric | 1 | 0 | no |
| COL | 4 | ant 2, fil 1 | 5/30 | 0 | leptotene | circular , centric | 1 | 0 | no |
| COL | 4 | fil2 | 5/30 | 0 | leptotene | circular , centric | 1 | 0 | no |
| COL | 4 | **bud 8**, ant1, fil1 | 0/30 | 0 | diakenesis | not visible | N/A | 100 | no |
| COL | 4 | fil 2 | 0/30 | 0 | diakenesis | not visible | N/A | 100 | no |
| COL | 4 | ant 2, fil 1 | 0/30 | 0 | m-l P | circular,peripheral | 2 | 100 | no |
| COL | 4 | fil2 | 0/30 | fbf | eP | circular,peripheral | 1 | 100 | no |

| | | | | | | | | | |
|---|---|---|---|---|---|---|---|---|---|
| COL | 4 | ant 3, fil 1 | 0/30 | 0 | m-l P | circular,peripheral | 3 | 100 | yes |
| COL | 4 | fil2 | 0/30 | 0 | m-l P | circular,peripheral | 3 | 100 | yes |
| COL | 4 | fil 3 | 0/30 | 0 | m-l P | circular,peripheral | 3 | 100 | yes |
| COL | 6 | **bud 1**, ant1, fil1 | 0/30 | 2 | zygotene | not visible | 1 | 1 to 25 | yes |
| COL | 6 | fil 2 | 0/30 | 2 | zygotene | not visible | 1 | 1 to 25 | yes |
| COL | 6 | fil 3 | 0/30 | 2 | zygotene | not visible | 1 | 1 to 25 | yes |
| COL | 6 | fil 4 | 0/30 | 2 | zygotene | not visible | 1 | 1 to 25 | yes |
| COL | 6 | ant 2, fil 1 | 5/30 | 1 | leptotene | circular, peripheral | 1 | 0 | yes |
| COL | 6 | fil2 | 15/30 | 1 | leptotene | circular, peripheral | 1 | 0 | yes |
| COL | 6 | fil 3 | 10/30 | 1 | leptotene | circular, peripheral | 1 | 0 | yes |
| COL | 6 | fil 4 | 5/30 | 1 | leptotene | circular, peripheral | 1 | 0 | yes |
| COL | 6 | ant 3, fil 1 | 25/30 | 0 | leptotene | circular, pericentric | 1 | 0 | yes |
| COL | 6 | fil2 | 15/30 | 0 | leptotene | circular, pericentric | 1 | 0 | yes |
| COL | 6 | fil 3 | 30/30 | 0 | leptotene | circular, pericentric | 1 | 0 | yes |
| COL | 6 | fil 4 | 25/30 | 0 | leptotene | circular, pericentric | 1 | 0 | yes |
| COL | 6 | **bud 2**, ant1, fil1 | 0/30 | 0 | m-l P | circular,peripheral | 3 | 100 | yes |
| COL | 6 | fil 2 | 0/30 | 0 | diplotene | not visible | N/A | 100 | no |
| COL | 6 | fil 3 | 0/30 | 0 | diplotene | not visible | N/A | 100 | no |
| COL | 6 | **bud 3**, ant1, fil1 | 0/30 | 0 | diplotene | not visible | N/A | 100 | no |
| COL | 6 | fil 2 | 0/30 | 0 | diplotene | not visible | N/A | 100 | no |
| COL | 6 | fil 3 | 0/30 | 0 | diplotene | not visible | N/A | 100 | no |
| COL | 6 | ant 2, fil 1 | 0/30 | 0 | m-l P | circular,peripheral | 3 | 100 | no |
| COL | 6 | fil2 | 0/30 | 0 | m-l P | circular,peripheral | 3 | 100 | no |
| COL | 6 | fil 3 | 0/30 | 0 | m-l P | circular,peripheral | 3 | 100 | no |
| COL | 6 | ant 3, fil 1 | 0/30 | fbf | eP | circular,peripheral | 1 | 100 | no |
| COL | 6 | fil2 | 0/30 | fbf | eP | circular,peripheral | 1 | 100 | no |
| COL | 6 | fil 3 | 0/30 | fbf | eP | circular,peripheral | 1 | 100 | no |
| COL | 6 | fil 4 | 0/30 | fbf | eP | circular,peripheral | 1 | 100 | no |

| | | | | | | | | | |
|---|---|---|---|---|---|---|---|---|---|
| COL | 6 | **bud 4**, ant1, fil1 | 0/30 | fbf | eP | circular,peripheral | 1 | 100 | no |
| COL | 6 | fil 2 | 0/30 | fbf | eP | circular,peripheral | 1 | 100 | no |
| COL | 6 | fil 3 | 0/30 | fbf | eP | circular,peripheral | 1 | 100 | no |
| COL | 6 | ant 2, fil 1 | 0/30 | fbf | eP | circular,peripheral | 1 | 100 | no |
| COL | 6 | fil2 | 0/30 | fbf | eP | circular,peripheral | 1 | 100 | no |
| COL | 6 | fil 3 | 0/30 | fbf | eP | circular,peripheral | 1 | 100 | no |
| COL | 6 | fil 4 | 0/30 | fbf | eP | circular,peripheral | 1 | 100 | no |
| COL | 6 | **bud 5**, ant1, fil1 | 0/30 | fbf | eP | circular,peripheral | 1 | 100 | no |
| COL | 6 | fil 2 | 0/30 | fbf | eP | circular,peripheral | 1 | 100 | no |
| COL | 6 | fil 3 | 0/30 | fbf | eP | circular,peripheral | 1 | 100 | no |
| COL | 6 | ant 2, fil 1 | 0/30 | fbf | eP | circular,peripheral | 1 | 100 | no |
| COL | 6 | fil2 | 0/30 | fbf | eP | circular,peripheral | 1 | 100 | no |
| COL | 6 | fil 3 | 0/30 | fbf | eP | circular,peripheral | 1 | 100 | no |
| COL | 8 | **bud 1**, ant1, fil1 | 0/30 | 0 | G2 | circular , centric | 1 | 0 | no |
| COL | 8 | fil 2 | 0/30 | 0 | G2 | circular , centric | 1 | 0 | no |
| COL | 8 | ant 2, fil 1 | 0/30 | 0 | G2 | circular , centric | 1 | 0 | no |
| COL | 8 | fil2 | 0/30 | 0 | G2 | circular , centric | 1 | 0 | no |
| COL | 8 | **bud 2**, ant1, fil1 | 0/30 | 0 | G2 | circular , centric | 1 | 0 | no |
| COL | 8 | fil 2 | 0/30 | 0 | G2 | circular , centric | 1 | 0 | no |
| COL | 8 | fil 3 | 0/30 | 0 | G2 | circular , centric | 1 | 0 | no |
| COL | 8 | ant 2, fil 1 | 0/30 | 0 | G2 | circular , centric | 1 | 0 | no |
| COL | 8 | fil2 | 0/30 | 0 | G2 | circular , centric | 1 | 0 | no |
| COL | 8 | fil 3 | 0/30 | 0 | G2 | circular , centric | 1 | 0 | no |
| COL | 8 | ant 3, fil 1 | 0/30 | 0 | G2 | circular , centric | 1 | 0 | no |
| COL | 8 | fil2 | 0/30 | 0 | G2 | circular , centric | 1 | 0 | no |
| COL | 8 | fil 3 | 0/30 | 0 | G2 | circular , centric | 1 | 0 | no |
| COL | 8 | **bud 3**, ant1, fil1 | 0/30 | 0 | diplotene | not visible | N/A | 100 | no |
| COL | 8 | fil 2 | 0/30 | 0 | diplotene | not visible | N/A | 100 | no |

| | | | | | | | | | |
|---|---|---|---|---|---|---|---|---|---|
| COL | 8 | ant 2, fil 1 | 0/30 | 0 | m-l P | circular,peripheral | 3 | 100 | yes |
| COL | 8 | fil2 | 0/30 | 0 | m-l P | circular,peripheral | 3 | 100 | yes |
| COL | 8 | fil 3 | 0/30 | 0 | m-l P | circular,peripheral | 3 | 100 | yes |
| COL | 8 | **bud 4**, ant1, fil1 | 0/30 | fbf | eP | circular,peripheral | 1 | 100 | no |
| COL | 8 | fil 2 | 0/30 | fbf | eP | circular,peripheral | 1 | 100 | no |
| COL | 8 | ant 2, fil 1 | 10/30 | 0 | leptotene | circular, pericentric | 1 | 0 | yes |
| COL | 8 | fil2 | 15/30 | 0 | leptotene | circular, pericentric | 1 | 0 | yes |
| COL | 8 | fil 3 | 15/30 | 0 | leptotene | circular, pericentric | 1 | 0 | yes |
| COL | 8 | **bud 5**, ant1, fil1 | 0/30 | 2 | zygotene | not visible | 1 | 26-75 | yes |
| COL | 8 | fil 2 | 0/30 | 2 | zygotene | not visible | 1 | 26-75 | yes |
| COL | 8 | fil 3 | 0/30 | 2 | zygotene | not visible | 1 | 26-75 | yes |
| COL | 8 | ant 2, fil 1 | 0/30 | 2 | zygotene | not visible | 1 | 1 to 25 | yes |
| COL | 8 | fil2 | 0/30 | 2 | zygotene | not visible | 1 | 1 to 25 | yes |
| COL | 10 | **bud 1**, ant1, fil1 | 0/30 | 0 | tetrad | not visible | N/A | 100 | no |
| COL | 10 | fil 2 | 0/30 | 0 | anaphase II | not visible | N/A | 100 | no |
| COL | 10 | fil 3 | 0/30 | 0 | anaphase II | not visible | N/A | 100 | no |
| COL | 10 | ant 2, fil 1 | 0/30 | 0 | tetrad | not visible | N/A | 100 | no |
| COL | 10 | fil2 | 0/30 | 0 | tetrad | not visible | N/A | 100 | no |
| COL | 10 | fil 3 | 0/30 | 0 | tetrad | not visible | N/A | 100 | no |
| COL | 10 | ant 3, fil 1 | 0/30 | 0 | diakenesis | not visible | N/A | 100 | no |
| COL | 10 | fil2 | 0/30 | 0 | diakenesis | not visible | N/A | 100 | no |
| COL | 10 | fil 3 | 0/30 | 0 | diakenesis | not visible | N/A | 100 | no |
| COL | 10 | **bud 2**, ant1, fil1 | 0/30 | fbf | eP | circular,peripheral | 1 | 100 | no |
| COL | 10 | fil 2 | 0/30 | fbf | eP | circular,peripheral | 1 | 100 | no |
| COL | 10 | fil 3 | 0/30 | fbf | eP | circular,peripheral | 1 | 100 | no |
| **COL** | **10** | **ant 2, fil 1** | **10/30** | **2** | **zygotene** | **not visible** | **1** | **1 to 25** | **yes** |
| **COL** | **10** | **fil2** | **5/30** | **2** | **zygotene** | **not visible** | **1** | **1 to 25** | **yes** |
| **COL** | **10** | **fil 3** | **10/30** | **2** | **zygotene** | **not visible** | **1** | **1 to 25** | **yes** |

| | | | | | | | | | |
|---|---|---|---|---|---|---|---|---|---|
| **COL** | **10** | **ant 3, fil 1** | **10/30** | **2** | **zygotene** | **not visible** | **1** | **1 to 25** | **yes** |
| **COL** | **10** | **fil2** | **10/30** | **2** | **zygotene** | **not visible** | **1** | **1 to 25** | **yes** |
| **COL** | **10** | **fil 3** | **10/30** | **2** | **zygotene** | **not visible** | **1** | **1 to 25** | **yes** |
| COL | 10 | **bud 3**, ant1, fil1 | 0/30 | 0 | m-l P | circular,peripheral | 3 | 100 | no |
| COL | 10 | fil 2 | 0/30 | 0 | m-l P | circular,peripheral | 3 | 100 | no |
| COL | 10 | fil 3 | 0/30 | 0 | m-l P | circular,peripheral | 3 | 100 | no |
| COL | 10 | ant 2, fil 1 | 0/30 | 0 | diakenesis | not visible | N/A | 100 | no |
| COL | 10 | fil2 | 0/30 | 0 | diakenesis | not visible | N/A | 100 | no |
| COL | 10 | fil 3 | 0/30 | 0 | diakenesis | not visible | N/A | 100 | no |
| COL | 10 | bud 4, ant1, fil1 | 5/30 | 2 | zygotene | not visible | 1 | 1 to 25 | yes |
| COL | 10 | fil 2 | 5/30 | 2 | zygotene | not visible | 1 | 1 to 25 | yes |
| COL | 10 | fil 3 | 5/30 | 2 | zygotene | not visible | 1 | 1 to 25 | yes |
| COL | 10 | ant 2, fil 1 | 30/30 | 0 | leptotene | circular, pericentric | 1 | 0 | yes |
| COL | 10 | fil2 | 30/30 | 0 | leptotene | circular, pericentric | 1 | 0 | yes |
| COL | 10 | fil 3 | 30/30 | 0 | leptotene | circular, pericentric | 1 | 0 | yes |
| COL | 10 | ant 3, fil 1 | 5/30 | 2 | zygotene | not visible | 1 | 1 to 25 | yes |
| COL | 10 | fil2 | 5/30 | 2 | zygotene | not visible | 1 | 1 to 25 | yes |
| COL | 10 | fil 3 | 30/30 | 0 | leptotene | circular, pericentric | 1 | 0 | no |
| COL | 10 | fil 4 | 30/30 | 0 | leptotene | circular, pericentric | 1 | 0 | no |
| COL | 10 | **bud 5**, ant1, fil1 | 30/30 | 0 | leptotene | circular, pericentric | 1 | 0 | no |
| COL | 10 | fil 2 | 30/30 | 0 | leptotene | circular, pericentric | 1 | 0 | no |
| COL | 10 | fil 3 | 30/30 | 0 | leptotene | circular, pericentric | 1 | 0 | no |
| COL | 10 | fil 4 | 30/30 | 0 | leptotene | circular, pericentric | 1 | 0 | no |
| COL | 10 | **bud 6**, ant1, fil1 | 0/30 | 0 | leptotene | circular, pericentric | 1 | 0 | no |
| COL | 10 | fil 2 | 0/30 | 0 | leptotene | circular, pericentric | 1 | 0 | no |
| COL | 10 | fil 3 | 0/30 | 0 | leptotene | circular, pericentric | 1 | 0 | no |
| COL | 10 | fil 4 | 0/30 | 0 | leptotene | circular, pericentric | 1 | 0 | no |
| COL | 10 | ant 2, fil 1 | 0/30 | 0 | leptotene | circular, pericentric | 1 | 0 | no |
| COL | 10 | fil2 | 0/30 | 0 | leptotene | circular, pericentric | 1 | 0 | no |

| | | | | | | | | | |
|---|---|---|---|---|---|---|---|---|---|
| COL | 10 | fil 3 | 0/30 | 0 | leptotene | circular, pericentric | 1 | 0 | no |
| COL | 10 | ant 3, fil 1 | 0/30 | 0 | leptotene | circular, pericentric | 1 | 0 | no |
| COL | 10 | fil2 | 0/30 | 0 | leptotene | circular, pericentric | 1 | 0 | no |
| COL | 10 | fil 3 | 0/30 | 0 | leptotene | circular, pericentric | 1 | 0 | no |
| COL | 10 | fil 4 | 0/30 | 0 | leptotene | circular, pericentric | 1 | 0 | no |
| COL | 10 | **bud 7**, ant1, fil1 | 0/30 | 2 | zygotene | not visible | 1 | 1 to 25 | yes |
| COL | 10 | fil 2 | 0/30 | 2 | zygotene | not visible | 1 | 1 to 25 | yes |
| COL | 10 | fil 3 | 0/30 | 2 | zygotene | not visible | 1 | 1 to 25 | yes |
| COL | 10 | ant 2, fil 1 | 0/30 | fbf | eP | circular,peripheral | 1 | 100 | no |
| COL | 10 | fil2 | 0/30 | fbf | eP | circular,peripheral | 1 | 100 | no |
| COL | 10 | fil 3 | 0/30 | fbf | eP | circular,peripheral | 1 | 100 | no |
| COL | 10 | ant 3, fil 1 | 0/30 | 2 | zygotene | not visible | 1 | 26-75 | yes |
| COL | 10 | fil2 | 0/30 | 2 | zygotene | not visible | 1 | 26-75 | yes |
| COL | 10 | fil 3 | 0/30 | 2 | zygotene | not visible | 1 | 26-75 | yes |
| COL | 10 | fil 4 | 0/30 | 2 | zygotene | not visible | 1 | 26-75 | yes |
| COL | 10 | **bud 8**, ant1, fil1 | 0/30 | 0 | diakenesis | not visible | N/A | 100 | no |
| COL | 10 | fil 2 | 0/30 | 0 | diakenesis | not visible | N/A | 100 | no |
| COL | 10 | fil 3 | 0/30 | 0 | diakenesis | not visible | N/A | 100 | no |
| COL | 12 | **bud 1**, ant1, fil1 | 5/30 | 0 | G2 | circular , centric | 1 | 0 | no |
| COL | 12 | fil 2 | 5/30 | 0 | G2 | circular , centric | 1 | 0 | no |
| COL | 12 | fil 3 | 5/30 | 0 | G2 | circular , centric | 1 | 0 | no |
| COL | 12 | ant 2, fil 1 | 0/30 | 0 | G2 | circular , centric | 1 | 0 | no |
| COL | 12 | fil2 | 0/30 | 0 | G2 | circular , centric | 1 | 0 | no |
| COL | 12 | fil 3 | 0/30 | 0 | G2 | circular , centric | 1 | 0 | no |
| COL | 12 | **bud 2**, ant1, fil1 | 30/30 | 0 | leptotene | circular, pericentric | 1 | 0 | no |
| COL | 12 | fil 2 | 30/30 | 0 | leptotene | circular, pericentric | 1 | 0 | no |
| COL | 12 | fil 3 | 30/30 | 0 | leptotene | circular, pericentric | 1 | 0 | no |
| COL | 12 | ant 2, fil 1 | 15/30 | 1 | leptotene | circular, peripheral | 1 | 0 | yes |

| | | | | | | | | | |
|---|---|---|---|---|---|---|---|---|---|
| COL | 12 | fil2 | 15/30 | 1 | leptotene | circular, peripheral | 1 | 0 | yes |
| COL | 12 | fil 3 | 20/30 | 1 | leptotene | circular, peripheral | 1 | 0 | yes |
| COL | 12 | fil 4 | 25/30 | 1 | leptotene | circular, peripheral | 1 | 0 | yes |
| COL | 12 | ant 3, fil 1 | 20/30 | 1 | leptotene | circular, peripheral | 1 | 0 | yes |
| COL | 12 | fil2 | 20/30 | 1 | leptotene | circular, peripheral | 1 | 0 | yes |
| COL | 12 | fil 3 | 20/30 | 1 | leptotene | circular, peripheral | 1 | 0 | yes |
| COL | 12 | ant 4, fil 1 | 30/30 | 2 | zygotene | not visible | 1 | 1 to 25 | yes |
| COL | 12 | fil2 | 30/30 | 2 | zygotene | not visible | 1 | 1 to 25 | yes |
| COL | 12 | fil 3 | 30/30 | 2 | zygotene | not visible | 1 | 1 to 25 | yes |
| COL | 12 | **bud 3**, ant1, fil1 | 30/30 | 1 | leptotene | circular, peripheral | 1 | 0 | yes |
| COL | 12 | fil 2 | 30/30 | 1 | leptotene | circular, peripheral | 1 | 0 | yes |
| COL | 12 | fil 3 | 30/30 | 1 | leptotene | circular, peripheral | 1 | 0 | yes |
| COL | 12 | ant 2, fil 1 | 30/30 | 1 | leptotene | circular, peripheral | 1 | 0 | yes |
| COL | 12 | fil2 | 30/30 | 1 | leptotene | circular, peripheral | 1 | 0 | yes |
| COL | 12 | fil 3 | 30/30 | 1 | leptotene | circular, peripheral | 1 | 0 | yes |
| COL | 12 | ant 3, fil 1 | 30/30 | 1 | leptotene | circular, peripheral | 1 | 0 | yes |
| COL | 12 | fil2 | 30/30 | 1 | leptotene | circular, peripheral | 1 | 0 | yes |
| COL | 12 | fil 3 | 30/30 | 1 | leptotene | circular, peripheral | 1 | 0 | yes |
| COL | 12 | **bud 4**, ant1, fil1 | 0/30 | fbf | eP | circular,peripheral | 1 | 100 | no |
| COL | 12 | fil 2 | 0/30 | fbf | eP | circular,peripheral | 1 | 100 | no |
| COL | 12 | fil 3 | 0/30 | fbf | eP | circular,peripheral | 1 | 100 | no |
| COL | 12 | ant 2, fil 1 | 0/30 | 0 | diplotene | not visible | N/A | 100 | no |
| COL | 12 | fil2 | 0/30 | 0 | diplotene | not visible | N/A | 100 | no |
| COL | 12 | ant 3, fil 1 | 0/30 | fbf | eP | circular,peripheral | 1 | 100 | no |
| COL | 12 | fil2 | 0/30 | fbf | eP | circular,peripheral | 1 | 100 | no |
| COL | 12 | fil 3 | 0/30 | fbf | eP | circular,peripheral | 1 | 100 | no |
| COL | 12 | **bud 5**, ant1, fil1 | 0/30 | 0 | G2 | circular , centric | 1 | 0 | no |
| COL | 12 | fil 2 | 0/30 | 0 | G2 | circular , centric | 1 | 0 | no |
| COL | 12 | fil 3 | 0/30 | 0 | G2 | circular , centric | 1 | 0 | no |

| | | | | | | | | | |
|---|---|---|---|---|---|---|---|---|---|
| COL | 12 | **bud 6**, ant1, fil1 | 0/30 | 0 | m-l P | circular,peripheral | 2 | 100 | no |
| COL | 12 | fil 2 | 0/30 | 0 | m-l P | circular,peripheral | 2 | 100 | no |
| COL | 12 | fil 3 | 0/30 | 0 | m-l P | circular,peripheral | 2 | 100 | no |
| COL | 12 | fil 4 | 0/30 | 0 | m-l P | circular,peripheral | 2 | 100 | no |
| COL | 12 | ant 2, fil 1 | 0/30 | 1 | zygotene | not visible | 1 | 76-99 | no |
| COL | 12 | fil2 | 0/30 | 1 | zygotene | not visible | 1 | 76-99 | no |
| COL | 12 | fil 3 | 0/30 | 1 | zygotene | not visible | 1 | 76-99 | no |
| COL | 12 | fil 4 | 0/30 | 1 | zygotene | not visible | 1 | 76-99 | no |
| COL | 14 | **bud 1**, ant1, fil1 | 0/30 | 0 | G2 | circular , centric | 1 | 0 | no |
| COL | 14 | fil 2 | 0/30 | 0 | G2 | circular , centric | 1 | 0 | no |
| COL | 14 | ant 2, fil 1 | 0/30 | 0 | G2 | circular , centric | 1 | 0 | no |
| COL | 14 | fil2 | 0/30 | 0 | G2 | circular , centric | 1 | 0 | no |
| COL | 14 | ant 3, fil 1 | 0/30 | 0 | G2 | circular , centric | 1 | 0 | no |
| COL | 14 | fil2 | 0/30 | 0 | G2 | circular , centric | 1 | 0 | no |
| COL | 14 | **bud 2**, ant1, fil1 | 0/30 | 0 | G2 | circular , centric | 1 | 0 | no |
| COL | 14 | fil 2 | 0/30 | 0 | G2 | circular , centric | 1 | 0 | no |
| COL | 14 | fil 3 | 0/30 | 0 | G2 | circular , centric | 1 | 0 | no |
| COL | 14 | ant 2, fil 1 | 0/30 | 0 | G2 | circular , centric | 1 | 0 | no |
| COL | 14 | fil2 | 0/30 | 0 | G2 | circular , centric | 1 | 0 | no |
| COL | 14 | fil 3 | 0/30 | 0 | G2 | circular , centric | 1 | 0 | no |
| COL | 14 | **bud 3,** ant1, fil1 | 30/30 | 1 | leptotene | circular, peripheral | 1 | 0 | no |
| COL | 14 | fil 2 | 30/30 | 1 | leptotene | circular, peripheral | 1 | 0 | no |
| COL | 14 | fil 3 | 30/30 | 1 | leptotene | circular, peripheral | 1 | 0 | no |
| COL | 14 | fil 4 | 30/30 | 1 | leptotene | circular, peripheral | 1 | 0 | no |
| COL | 14 | ant 2, fil 1 | 10/30 | 1 | leptotene | circular, peripheral | 1 | 0 | no |
| COL | 14 | fil2 | 20/30 | 1 | leptotene | circular, peripheral | 1 | 0 | no |
| COL | 14 | fil 3 | 20/30 | 1 | leptotene | circular, peripheral | 1 | 0 | no |
| COL | 14 | fil 4 | 20/30 | 1 | leptotene | circular, peripheral | 1 | 0 | no |

| | | | | | | | | | |
|---|---|---|---|---|---|---|---|---|---|
| COL | 14 | **bud 4**, ant1, fil1 | 0/30 | 0 | metaphase I | not visible | N/A | 100 | no |
| COL | 14 | fil 2 | 0/30 | 0 | metaphase I | not visible | N/A | 100 | no |
| COL | 14 | ant 2, fil 1 | 0/30 | 0 | tetrad | not visible | N/A | 100 | no |
| COL | 14 | fil2 | 0/30 | 0 | tetrad | not visible | N/A | 100 | no |
| COL | 14 | fil 3 | 0/30 | 0 | tetrad | not visible | N/A | 100 | no |
| COL | 14 | **bud 5**, ant1, fil1 | 0/30 | 0 | G2 | circular , centric | 1 | 0 | no |
| COL | 14 | fil 2 | 0/30 | 0 | G2 | circular , centric | 1 | 0 | no |
| COL | 14 | fil 3 | 0/30 | 0 | G2 | circular , centric | 1 | 0 | no |
| COL | 14 | ant 2, fil 1 | 0/30 | 0 | G2 | circular , centric | 1 | 0 | no |
| COL | 14 | fil2 | 0/30 | 0 | G2 | circular , centric | 1 | 0 | no |
| COL | 14 | fil 3 | 0/30 | 0 | G2 | circular , centric | 1 | 0 | no |
| COL | 14 | **bud 6,** ant1, fil1 | 0/30 | 2 | zygotene | not visible | 1 | 26-75 | yes |
| COL | 14 | fil 2 | 0/30 | 2 | zygotene | not visible | 1 | 26-75 | yes |
| COL | 14 | fil 3 | 0/30 | 2 | zygotene | not visible | 1 | 26-75 | yes |
| COL | 14 | fil 4 | 0/30 | 2 | zygotene | not visible | 1 | 26-75 | yes |
| COL | 14 | ant 2, fil 1 | 0/30 | 2 | zygotene | not visible | 1 | 26-75 | yes |
| COL | 14 | fil2 | 0/30 | 2 | zygotene | not visible | 1 | 26-75 | yes |
| COL | 14 | fil 3 | 0/30 | 2 | zygotene | not visible | 1 | 26-75 | yes |
| COL | 14 | fil 4 | 0/30 | 2 | zygotene | not visible | 1 | 26-75 | yes |
| COL | 14 | **bud 7,** ant1, fil1 | 30/30 | 1 | leptotene | circular, peripheral | 1 | 0 | no |
| COL | 14 | fil 2 | 30/30 | 1 | leptotene | circular, peripheral | 1 | 0 | no |
| COL | 14 | fil 3 | 30/30 | 1 | leptotene | circular, peripheral | 1 | 0 | no |
| COL | 14 | fil 4 | 30/30 | 1 | leptotene | circular, peripheral | 1 | 0 | no |
| COL | 14 | ant 2, fil 1 | 0/30 | 0 | G2 | circular , centric | 1 | 0 | no |
| COL | 14 | fil2 | 0/30 | 0 | G2 | circular , centric | 1 | 0 | no |
| COL | 14 | fil 3 | 0/30 | 0 | G2 | circular , centric | 1 | 0 | no |
| COL | 16 | **bud 1,** ant1, fil1 | 0/30 | 0 | m-l P | circular,peripheral | 3 | 100 | no |
| COL | 16 | fil 2 | 0/30 | 0 | m-l P | circular,peripheral | 3 | 100 | no |

| | | | | | | | | | |
|---|---|---|---|---|---|---|---|---|---|
| COL | 16 | ant 2, fil 1 | 0/30 | 0 | m-l P | circular,peripheral | 3 | 100 | no |
| COL | 16 | fil2 | 0/30 | 0 | m-l P | circular,peripheral | 3 | 100 | no |
| COL | 16 | fil 3 | 0/30 | 0 | m-l P | circular,peripheral | 3 | 100 | no |
| COL | 16 | fil 4 | 0/30 | 0 | m-l P | circular,peripheral | 3 | 100 | no |
| COL | 16 | **bud 2,** ant1, fil1 | 0/30 | fbf | eP | circular,peripheral | 1 | 100 | no |
| COL | 16 | fil 2 | 0/30 | fbf | eP | circular,peripheral | 1 | 100 | no |
| COL | 16 | fil 3 | 0/30 | fbf | eP | circular,peripheral | 1 | 100 | no |
| COL | 16 | fil 4 | 0/30 | fbf | eP | circular,peripheral | 1 | 100 | no |
| COL | 16 | **bud 3,** ant1, fil1 | 30/30 | 1 | leptotene | circular, peripheral | 1 | 0 | yes |
| COL | 16 | fil 2 | 30/30 | 1 | leptotene | circular, peripheral | 1 | 0 | yes |
| COL | 16 | fil 3 | 30/30 | 0 | leptotene | circular, pericentric | 1 | 0 | no |
| COL | 16 | fil 4 | 30/30 | 0 | leptotene | circular, pericentric | 1 | 0 | no |
| COL | 16 | ant 2, fil 1 | 30/30 | 2 | zygotene | not visible | 1 | 1 to 25 | yes |
| COL | 16 | fil2 | 30/30 | 2 | zygotene | not visible | 1 | 1 to 25 | yes |
| COL | 16 | fil 3 | 30/30 | 2 | zygotene | not visible | 1 | 1 to 25 | yes |
| COL | 16 | **bud 4,** ant1, fil1 | 30/30 | 2 | zygotene | not visible | 1 | 1 to 25 | yes |
| COL | 16 | fil 2 | 30/30 | 2 | zygotene | not visible | 1 | 1 to 25 | yes |
| COL | 16 | fil 3 | 30/30 | 2 | zygotene | not visible | 1 | 1 to 25 | yes |
| COL | 16 | ant 2, fil 1 | 0/30 | 0 | m-l P | circular,peripheral | 3 | 100 | no |
| COL | 16 | fil2 | 0/30 | 0 | m-l P | circular,peripheral | 3 | 100 | no |
| COL | 16 | ant 3, fil 1 | 0/30 | 0 | m-l P | circular,peripheral | 3 | 100 | no |
| COL | 16 | fil2 | 0/30 | 0 | m-l P | circular,peripheral | 3 | 100 | no |
| COL | 16 | fil 3 | 0/30 | 0 | m-l P | circular,peripheral | 3 | 100 | no |
| COL | 16 | **bud 5,** ant1, fil1 | 0/30 | 0 | metaphase I | not visible | N/A | 100 | no |
| COL | 16 | fil 2 | 0/30 | 0 | metaphase I | not visible | N/A | 100 | no |
| COL | 16 | **bud 6,** ant1, fil1 | 20/30 | 2 | zygotene | not visible | 1 | 26-75 | yes |
| COL | 16 | fil 2 | 20/30 | 2 | zygotene | not visible | 1 | 26-75 | yes |
| COL | 18 | **bud 1,** ant1, fil1 | 30/30 | 1 | zygotene | not visible | 1 | 76-99 | no |

| | | | | | | | | | |
|---|---|---|---|---|---|---|---|---|---|
| **COL** | **18** | **fil 2** | **15/30** | **fbf** | **eP** | **circular,peripheral** | **1** | **100** | **yes** |
| COL | 18 | fil 3 | 30/30 | 1 | zygotene | not visible | 1 | 76-99 | no |
| COL | 18 | ant 2, fil 1 | 0/30 | 0 | m-l P | circular,peripheral | 3 | 100 | yes |
| COL | 18 | fil2 | 0/30 | 0 | diplotene | not visible | N/A | 100 | no |
| COL | 18 | ant 3, fil 1 | 0/30 | 0 | diplotene | not visible | N/A | 100 | no |
| COL | 18 | fil2 | 15/30 | fbf | eP | circular,peripheral | 1 | 100 | yes |
| COL | 18 | ant 4, fil 1 | 0/30 | fbf | eP | circular,peripheral | 1 | 100 | yes |
| COL | 18 | fil2 | 0/30 | 0 | m-l P | circular,peripheral | 3 | 100 | yes |
| COL | 18 | fil 3 | 0/30 | 0 | m-l P | circular,peripheral | 3 | 100 | yes |
| COL | 20 | **bud 1,** ant1, fil1 | 0/30 | 0 | anaphase I | not visible | N/A | 100 | no |
| COL | 20 | fil 2 | 0/30 | 0 | anaphase I | not visible | N/A | 100 | no |
| COL | 20 | fil 3 | 0/30 | 0 | anaphase I | not visible | N/A | 100 | no |
| COL | 20 | ant 2, fil 1 | 0/30 | 0 | tetrad | not visible | N/A | 100 | no |
| COL | 20 | fil2 | 0/30 | 0 | tetrad | not visible | N/A | 100 | no |
| COL | 20 | ant 3, fil 1 | 0/30 | 0 | m-l P | circular,peripheral | 3 | 100 | no |
| COL | 20 | fil2 | 0/30 | 0 | m-l P | circular,peripheral | 3 | 100 | no |
| COL | 20 | fil 3 | 0/30 | 0 | m-l P | circular,peripheral | 3 | 100 | no |
| COL | 20 | fil 4 | 0/30 | 0 | m-l P | circular,peripheral | 3 | 100 | no |
| COL | 20 | **bud 2,** ant1, fil1 | 0/30 | 0 | G2 | circular , centric | 1 | 0 | no |
| COL | 20 | fil 2 | 0/30 | 0 | G2 | circular , centric | 1 | 0 | no |
| COL | 20 | ant 2, fil 1 | 0/30 | 0 | G2 | circular , centric | 1 | 0 | no |
| COL | 20 | fil2 | 0/30 | 0 | G2 | circular , centric | 1 | 0 | no |
| COL | 20 | fil 3 | 0/30 | 0 | G2 | circular , centric | 1 | 0 | no |
| COL | 20 | **bud 3,** ant1, fil1 | 30/30 | 0 | leptotene | circular, pericentric | 1 | 0 | no |
| COL | 20 | fil 2 | 30/30 | 0 | leptotene | circular, pericentric | 1 | 0 | no |
| COL | 20 | **bud 4,** ant1, fil1 | 0/30 | 0 | metaphase I | not visible | N/A | 100 | no |
| COL | 20 | fil 2 | 0/30 | 0 | metaphase I | not visible | N/A | 100 | no |
| COL | 20 | fil 3 | 0/30 | 0 | metaphase I | not visible | N/A | 100 | no |

| | | | | | | | | | |
|---|---|---|---|---|---|---|---|---|---|
| COL | 22 | **bud 1,** ant1, fil1 | 30/30 | 2 | zygotene | not visible | 1 | 1 to 25 | yes |
| COL | 22 | fil 2 | 30/30 | 2 | zygotene | not visible | 1 | 1 to 25 | yes |
| COL | 22 | fil 3 | 30/30 | 2 | zygotene | not visible | 1 | 1 to 25 | yes |
| COL | 22 | ant 2, fil 1 | 30/30 | 1 | leptotene | circular, peripheral | 1 | 0 | yes |
| COL | 22 | fil2 | 30/30 | 1 | leptotene | circular, peripheral | 1 | 0 | yes |
| COL | 22 | fil 3 | 30/30 | 1 | leptotene | circular, peripheral | 1 | 0 | yes |
| COL | 22 | fil 4 | 30/30 | 1 | leptotene | circular, peripheral | 1 | 0 | yes |
| COL | 22 | **bud 2,** ant1, fil1 | 0/30 | fbf | eP | circular,peripheral | 1 | 100 | no |
| COL | 22 | fil 2 | 0/30 | fbf | eP | circular,peripheral | 1 | 100 | no |
| COL | 22 | fil 3 | 0/30 | fbf | eP | circular,peripheral | 1 | 100 | no |
| COL | 22 | **bud 3,** ant1, fil1 | 30/30 | 0 | leptotene | circular, pericentric | 1 | 0 | no |
| COL | 22 | fil 2 | 30/30 | 0 | leptotene | circular, pericentric | 1 | 0 | no |
| COL | 22 | fil 3 | 30/30 | 0 | leptotene | circular, pericentric | 1 | 0 | no |
| COL | 22 | ant 2, fil 1 | 30/30 | 2 | zygotene | not visible | 1 | 1 to 25 | yes |
| COL | 22 | fil2 | 30/30 | 2 | zygotene | not visible | 1 | 1 to 25 | yes |
| COL | 22 | **bud 4,** ant1, fil1 | 0/30 | 0 | m-l P | circular,peripheral | 3 | 100 | no |
| COL | 22 | fil 2 | 0/30 | 0 | m-l P | circular,peripheral | 3 | 100 | no |
| COL | 22 | fil 3 | 0/30 | 0 | m-l P | circular,peripheral | 3 | 100 | no |
| COL | 22 | **bud 5,** ant1, fil1 | 0/30 | 0 | metaphase I | not visible | N/A | 100 | no |
| COL | 22 | fil 2 | 0/30 | 0 | metaphase I | not visible | N/A | 100 | no |
| COL | 22 | fil 3 | 0/30 | 0 | metaphase I | not visible | N/A | 100 | no |
| COL | 24 | **bud 1,** ant1, fil1 | 0/30 | 0 | m-l P | circular,peripheral | 3 | 100 | no |
| COL | 24 | fil 2 | 0/30 | 0 | m-l P | circular,peripheral | 3 | 100 | no |
| COL | 24 | fil 3 | 0/30 | 0 | m-l P | circular,peripheral | 3 | 100 | no |
| COL | 24 | ant 2, fil 1 | 30/30 | 2 | zygotene | not visible | 1 | 1 to 25 | yes |
| COL | 24 | fil2 | 30/30 | 2 | zygotene | not visible | 1 | 1 to 25 | yes |
| COL | 24 | fil 3 | 30/30 | 2 | zygotene | not visible | 1 | 1 to 25 | yes |
| COL | 24 | ant 3, fil 1 | 30/30 | 0 | m-l P | circular,peripheral | 2 | 100 | yes |

| | | | | | | | | | |
|---|---|---|---|---|---|---|---|---|---|
| COL | 24 | fil2 | 30/30 | 0 | m-l P | circular,peripheral | 2 | 100 | yes |
| COL | 24 | fil 3 | 30/30 | 0 | m-l P | circular,peripheral | 2 | 100 | yes |
| COL | 24 | fil 4 | 30/30 | 0 | m-l P | circular,peripheral | 2 | 100 | yes |
| COL | 26 | **bud 1,** ant1, fil1 | 30/30 | fbf | eP | circular,peripheral | 1 | 100 | no |
| COL | 26 | fil 2 | 30/30 | fbf | eP | circular,peripheral | 1 | 100 | no |
| COL | 26 | fil 3 | 30/30 | fbf | eP | circular,peripheral | 1 | 100 | no |
| COL | 26 | ant 2, fil 1 | 30/30 | fbf | eP | circular,peripheral | 1 | 100 | no |
| COL | 26 | fil2 | 30/30 | fbf | eP | circular,peripheral | 1 | 100 | no |
| COL | 26 | fil 3 | 30/30 | fbf | eP | circular,peripheral | 1 | 100 | no |
| COL | 26 | **bud 2,** ant1, fil1 | 0/30 | 0 | leptotene | circular , centric | 1 | 0 | no |
| COL | 26 | fil 2 | 0/30 | 0 | leptotene | circular , centric | 1 | 0 | no |
| COL | 26 | fil 3 | 0/30 | 0 | leptotene | circular , centric | 1 | 0 | no |
| COL | 26 | ant 2, fil 1 | 0/30 | 0 | leptotene | circular , centric | 1 | 0 | no |
| COL | 26 | fil2 | 0/30 | 0 | leptotene | circular , centric | 1 | 0 | no |
| COL | 26 | fil 3 | 0/30 | 0 | leptotene | circular , centric | 1 | 0 | no |
| COL | 26 | ant 3, fil 1 | 0/30 | 0 | leptotene | circular , centric | 1 | 0 | no |
| COL | 26 | fil2 | 0/30 | 0 | leptotene | circular , centric | 1 | 0 | no |
| COL | 26 | fil 3 | 0/30 | 0 | leptotene | circular , centric | 1 | 0 | no |
| COL | 26 | ant 4, fil 1 | 0/30 | 0 | leptotene | circular , centric | 1 | 0 | no |
| COL | 26 | fil2 | 0/30 | 0 | leptotene | circular , centric | 1 | 0 | no |
| COL | 26 | fil 3 | 0/30 | 0 | leptotene | circular , centric | 1 | 0 | no |
| COL | 26 | **bud 3,** ant1, fil1 | 0/30 | 0 | m-l P | circular,peripheral | 3 | 100 | no |
| COL | 26 | fil 2 | 0/30 | 0 | m-l P | circular,peripheral | 3 | 100 | no |
| COL | 26 | fil 3 | 0/30 | 0 | m-l P | circular,peripheral | 3 | 100 | no |
| COL | 26 | **bud 4,** ant1, fil1 | 0/30 | 0 | G2 | circular , centric | 1 | 0 | no |
| COL | 26 | fil 2 | 0/30 | 0 | G2 | circular , centric | 1 | 0 | no |
| COL | 26 | fil 3 | 0/30 | 0 | G2 | circular , centric | 1 | 0 | no |
| COL | 26 | **bud 5,** ant1, fil1 | 0/30 | 0 | tetrad | not visible | N/A | 100 | no |

| | | | | | | | | | |
|---|---|---|---|---|---|---|---|---|---|
| COL | 26 | fil 2 | 0/30 | 0 | tetrad | not visible | N/A | 100 | no |
| COL | 26 | fil 3 | 0/30 | 0 | tetrad | not visible | N/A | 100 | no |
| COL | 26 | **bud 6,** ant1, fil1 | 30/30 | 2 | zygotene | not visible | 1 | 1 to 25 | yes |
| COL | 26 | fil 2 | 30/30 | 2 | zygotene | not visible | 1 | 1 to 25 | yes |
| COL | 26 | fil 3 | 30/30 | 2 | zygotene | not visible | 1 | 1 to 25 | yes |
| COL | 26 | **bud 7,** ant1, fil1 | 30/30 | 2 | zygotene | not visible | 1 | 1 to 25 | yes |
| COL | 26 | fil 2 | 30/30 | 2 | zygotene | not visible | 1 | 1 to 25 | yes |
| COL | 26 | fil 3 | 30/30 | 2 | zygotene | not visible | 1 | 1 to 25 | yes |
| COL | 28 | **bud 1,** ant1, fil1 | 0/30 | 0 | G2 | circular , centric | 1 | 0 | no |
| COL | 28 | fil 2 | 0/30 | 0 | G2 | circular , centric | 1 | 0 | no |
| COL | 28 | fil 3 | 0/30 | 0 | G2 | circular , centric | 1 | 0 | no |
| COL | 28 | ant 2, fil 1 | 0/30 | 0 | G2 | circular , centric | 1 | 0 | no |
| COL | 28 | fil2 | 0/30 | 0 | G2 | circular , centric | 1 | 0 | no |
| COL | 28 | fil 3 | 0/30 | 0 | G2 | circular , centric | 1 | 0 | no |
| COL | 28 | ant 3, fil 1 | 0/30 | 0 | G2 | circular , centric | 1 | 0 | no |
| COL | 28 | fil2 | 0/30 | 0 | G2 | circular , centric | 1 | 0 | no |
| COL | 28 | fil 3 | 0/30 | 0 | G2 | circular , centric | 1 | 0 | no |
| COL | 28 | **bud 2,** ant1, fil1 | 30/30 | 2 | zygotene | not visible | 1 | 76-99 | yes |
| COL | 28 | fil 2 | 30/30 | 2 | zygotene | not visible | 1 | 76-99 | yes |
| COL | 28 | fil 3 | 30/30 | 2 | zygotene | not visible | 1 | 76-99 | yes |
| COL | 30 | **bud 1,** ant1, fil1 | 0/30 | 0 | leptotene | circular , centric | 1 | 0 | no |
| COL | 30 | fil 2 | 0/30 | 0 | leptotene | circular , centric | 1 | 0 | no |
| COL | 30 | fil 3 | 0/30 | 0 | leptotene | circular , centric | 1 | 0 | no |
| COL | 30 | ant 2, fil 1 | 0/30 | 0 | leptotene | circular , centric | 1 | 0 | no |
| COL | 30 | fil2 | 0/30 | 0 | leptotene | circular , centric | 1 | 0 | no |
| COL | 30 | fil 3 | 0/30 | 0 | leptotene | circular , centric | 1 | 0 | no |
| COL | 30 | fil 4 | 0/30 | 0 | leptotene | circular , centric | 1 | 0 | no |
| COL | 30 | **bud 2,** ant1, fil1 | 30/30 | fbf | eP | circular,peripheral | 1 | 100 | no |

| | | | | | | | | | |
|---|---|---|---|---|---|---|---|---|---|
| COL | 30 | fil 2 | 30/30 | fbf | eP | circular,peripheral | 1 | 100 | no |
| COL | 30 | fil 3 | 30/30 | fbf | eP | circular,peripheral | 1 | 100 | no |
| COL | 32 | **bud 1,** ant1, fil1 | 30/30 | 0 | m-l P | circular,peripheral | 3 | 100 | no |
| COL | 32 | fil 2 | 30/30 | 0 | m-l P | circular,peripheral | 3 | 100 | no |
| COL | 32 | **bud 2,** ant1, fil1 | 0/30 | 0 | metaphase I | not visible | N/A | 100 | no |
| COL | 32 | fil 2 | 0/30 | 0 | metaphase I | not visible | N/A | 100 | no |

***mnd1* data begins here**

| Geno-type | Time point (hr) | Bud, Anther, Filament | # meiocyte with EdU signal/total | Level γH2AX signal | Meiotic stage. | Nucleolus shape & location | Amount Callose | % bi-nucleate. tapetum. | Tapetum labeled w/EdU |
|---|---|---|---|---|---|---|---|---|---|
| mnd1 | 0 | **bud 1**, ant1, fil1 | 0/30 | 0 | G2 | circular , centric | 1 | 0 | no |
| mnd1 | 0 | fil 2 | 0/30 | 0 | G2 | circular , centric | 1 | 0 | no |
| **mnd1** | **0** | **fil3** | **5/30** | **0** | **G2** | **circular , centric** | **1** | **0** | **no** |
| **mnd1** | **0** | **ant 2, fil 1** | **15/30** | **0** | **G2** | **circular , centric** | **1** | **0** | **no** |
| **mnd1** | **0** | **fil2** | **10/30** | **0** | **G2** | **circular , centric** | **1** | **0** | **no** |
| **mnd1** | **0** | **fil3** | **12/30** | **0** | **G2** | **circular , centric** | **1** | **0** | **no** |
| mnd1 | 0 | ant 3 fil1 | 0/30 | 0 | leptotene | circular , centric | 1 | 0 | no |
| mnd1 | 0 | | 0/30 | 0 | leptotene | circular , centric | 1 | 0 | no |
| mnd1 | 0 | | 0/30 | 0 | leptotene | circular , centric | 1 | 0 | no |
| mnd1 | 0 | bud 2 ant 1, fil 1 | 7/30 | 0 | G2 | circular , centric | 1 | 0 | no |
| mnd1 | 0 | fil2 | 0/30 | 0 | G2 | circular , centric | 1 | 0 | no |
| mnd1 | 0 | fil3 | 2/30 | 0 | G2 | circular , centric | 1 | 0 | no |
| mnd1 | 0 | ant 2 fil 1 | 2/30 | 0 | G2 | circular , centric | 1 | 0 | no |
| mnd1 | 0 | fil2 | 3/30 | 0 | G2 | circular , centric | 1 | 0 | no |
| mnd1 | 0 | fil3 | 2/30 | 0 | G2 | circular , centric | 1 | 0 | no |

| | | | | | | | | | |
|---|---|---|---|---|---|---|---|---|---|
| mnd1 | 0 | ant 3 fil 1 | 0/30 | 1 | leptotene | circular , centric | 1 | 0 | no |
| mnd1 | 0 | fil2 | 0/30 | 1 | leptotene | circular , centric | 1 | 0 | no |
| mnd1 | 0 | fil3 | 0/30 | 1 | leptotene | circular , centric | 1 | 0 | no |
| mnd1 | 0 | ant 4 fil 1 | 0/30 | 1 | leptotene | circular , centric | 1 | 0 | no |
| mnd1 | 0 | fil2 | 0/30 | 1 | leptotene | circular , centric | 1 | 0 | no |
| mnd1 | 0 | fil3 | 0/30 | 1 | leptotene | circular , centric | 1 | 0 | no |
| mnd1 | 2 | **bud 1**, ant1, fil1 | 0/30 | 1 | zygotene | not visible | 1 | 2575 | no |
| mnd1 | 2 | fil 2 | 0/30 | 1 | zygotene | not visible | 1 | 26-75 | no |
| mnd1 | 2 | fil3 | 0/30 | 1 | zygotene | not visible | 1 | 26-75 | no |
| mnd1 | 2 | ant2 fil 1 | 0/30 | 0 | m-l P | circular,peripheral | 2 | 100 | yes |
| mnd1 | 2 | fil 2 | 0/30 | 0 | m-l P | circular,peripheral | 2 | 100 | yes |
| mnd1 | 2 | fil 3 | 0/30 | 0 | m-l P | circular,peripheral | 2 | 100 | yes |
| mnd1 | 2 | **bud 2** ant 1 fil 1 | 0/30 | 0 | leptotene | circular , centric | 1 | 0 | no |
| mnd1 | 2 | fil 2 | 0/30 | 0 | leptotene | circular , centric | 1 | 0 | no |
| mnd1 | 2 | fil3 | 0/30 | 0 | leptotene | circular , centric | 1 | 0 | no |
| mnd1 | 2 | **bud 3** ant 1 fil 1 | 0/30 | 1 | leptotene | circular , centric | 1 | 0 | no |
| mnd1 | 2 | fil 2 | 0/30 | 1 | leptotene | circular , centric | 1 | 0 | no |
| mnd1 | 2 | fil 3 | 0/30 | 1 | leptotene | circular , centric | 1 | 0 | no |
| mnd1 | 2 | ant2 fil 1 | 0/30 | 0 | leptotene | circular , centric | 1 | 0 | no |
| mnd1 | 2 | fil 2 | 0/30 | 0 | leptotene | circular , centric | 1 | 0 | no |
| mnd1 | 2 | fil 3 | 0/30 | 0 | leptotene | circular , centric | 1 | 0 | no |
| mnd1 | 2 | **bud 4** ant 1 fil 1 | 0/30 | 0 | leptotene | circular , centric | 1 | 0 | no |
| mnd1 | 2 | fil 2 | 0/30 | 0 | leptotene | circular , centric | 1 | 0 | no |
| mnd1 | 2 | fil 3 | 0/30 | 0 | leptotene | circular , centric | 1 | 0 | no |
| mnd1 | 4 | **bud 1**, ant1, fil1 | 0/30 | 2 | zygotene | not visible | 1 | 1 to 25 | no |
| mnd1 | 4 | fil 2 | 0/30 | 2 | zygotene | not visible | 1 | 1 to 25 | no |
| mnd1 | 4 | fil3 | 0/30 | 2 | zygotene | not visible | 1 | 1 to 25 | no |
| mnd1 | 4 | ant2, fil1 | 15/30 | 0 | leptotene | circular , centric | 1 | 0 | no |

| | | | | | | | | | |
|---|---|---|---|---|---|---|---|---|---|
| mnd1 | 4 | fil 2 | 20/30 | 0 | leptotene | circular , centric | 1 | 0 | no |
| mnd1 | 4 | **bud 2**, ant1, fil1 | 0/30 | 2 | zygotene | not visible | 1 | 26-75 | no |
| mnd1 | 4 | fil2 | 0/30 | 2 | zygotene | not visible | 1 | 26-75 | no |
| mnd1 | 4 | fil3 | 0/30 | 2 | zygotene | not visible | 1 | 26-75 | no |
| mnd1 | 4 | ant2, fil1 | 0/30 | 1 | zygotene | not visible | 1 | 76-99 | yes |
| mnd1 | 4 | fil 2 | 0/30 | 1 | zygotene | not visible | 1 | 76-99 | yes |
| mnd1 | 4 | fil3 | 0/30 | 1 | zygotene | not visible | 1 | 76-99 | yes |
| mnd1 | 4 | ant3, fil1 | 0/30 | 1 | leptotene | circular , centric | 1 | 0 | no |
| mnd1 | 4 | fil 2 | 0/30 | 1 | leptotene | circular , centric | 1 | 0 | no |
| mnd1 | 4 | fil3 | 0/30 | 1 | leptotene | circular , centric | 1 | 0 | no |
| mnd1 | 4 | **bud 3**, ant1, fil1 | 30/30 | 0 | G2 | circular , centric | 1 | 0 | no |
| mnd1 | 4 | fil2 | 30/30 | 0 | G2 | circular , centric | 1 | 0 | no |
| mnd1 | 4 | fil3 | 30/30 | 0 | G2 | circular , centric | 1 | 0 | no |
| mnd1 | 4 | ant2, fil1 | 30/30 | 0 | G2 | circular , centric | 1 | 0 | no |
| mnd1 | 4 | fil 2 | 30/30 | 0 | G2 | circular , centric | 1 | 0 | no |
| mnd1 | 4 | fil3 | 30/30 | 0 | G2 | circular , centric | 1 | 0 | no |
| mnd1 | 4 | ant3, fil1 | 30/30 | 0 | G2 | circular , centric | 1 | 0 | no |
| mnd1 | 4 | fil 2 | 30/30 | 0 | G2 | circular , centric | 1 | 0 | no |
| mnd1 | 4 | fil3 | 30/30 | 0 | G2 | circular , centric | 1 | 0 | no |
| mnd1 | 4 | ant4, fil1 | 30/30 | 0 | G2 | circular , centric | 1 | 0 | no |
| mnd1 | 4 | fil 2 | 30/30 | 0 | G2 | circular , centric | 1 | 0 | no |
| mnd1 | 4 | fil3 | 30/30 | 0 | G2 | circular , centric | 1 | 0 | no |
| mnd1 | 4 | **bud4** ant1, fil1 | 0/30 | 0 | m-l P | circular,peripheral | 2 | 100 | yes |
| mnd1 | 4 | fil 2 | 0/30 | 0 | m-l P | circular,peripheral | 2 | 100 | yes |
| mnd1 | 4 | fil3 | 0/30 | 0 | m-l P | circular,peripheral | 2 | 100 | yes |
| mnd1 | 6 | **bud 1**, ant1, fil1 | 30/30 | 0 | leptotene | circular , centric | 1 | 0 | no |
| mnd1 | 6 | fil 2 | 30/30 | 0 | leptotene | circular , centric | 1 | 0 | no |
| mnd1 | 6 | fil3 | 30/30 | 0 | leptotene | circular , centric | 1 | 0 | no |

| | | | | | | | | | |
|---|---|---|---|---|---|---|---|---|---|
| mnd1 | 6 | **bud 2** ant1, fil1 | 0/30 | 0 | m-l P | circular,peripheral | 3 | 100 | no |
| mnd1 | 6 | fil 2 | 0/30 | 0 | m-l P | circular,peripheral | 3 | 100 | no |
| mnd1 | 6 | fil3 | 0/30 | 0 | m-l P | circular,peripheral | 3 | 100 | no |
| mnd1 | 6 | ant2, fil1 | 0/30 | 0 | m-l P | circular,peripheral | 3 | 100 | no |
| mnd1 | 6 | fil 2 | 0/30 | 0 | m-l P | circular,peripheral | 3 | 100 | no |
| mnd1 | 6 | fil3 | 0/30 | 0 | m-l P | circular,peripheral | 3 | 100 | no |
| mnd1 | 6 | ant3, fil1 | 0/30 | 2 | zygotene | not visible | 1 | 1 to 25 | yes |
| mnd1 | 6 | fil 2 | 0/30 | 2 | zygotene | not visible | 1 | 1 to 25 | yes |
| mnd1 | 6 | fil3 | 0/30 | 2 | zygotene | not visible | 1 | 1 to 25 | yes |
| mnd1 | 6 | ant4, fil1 | 0/30 | 0 | m-l P | circular,peripheral | 3 | 100 | no |
| mnd1 | 6 | fil 2 | 0/30 | 0 | m-l P | circular,peripheral | 3 | 100 | no |
| mnd1 | 6 | **bud 3**, ant1, fil1 | 0/30 | 1 | leptotene | circular , centric | 1 | 0 | no |
| mnd1 | 6 | fil2 | 0/30 | 1 | leptotene | circular , centric | 1 | 0 | no |
| mnd1 | 6 | fil3 | 0/30 | 1 | leptotene | circular , centric | 1 | 0 | no |
| mnd1 | 6 | ant2, fil1 | 25/30 | 1 | leptotene | circular , centric | 1 | 0 | no |
| mnd1 | 6 | fil 2 | 20/30 | 1 | leptotene | circular , centric | 1 | 0 | no |
| mnd1 | 6 | fil3 | 22/30 | 1 | leptotene | circular , centric | 1 | 0 | no |
| mnd1 | 6 | ant3, fil1 | 0/30 | 1 | leptotene | circular , centric | 1 | 0 | no |
| mnd1 | 6 | fil 2 | 0/30 | 1 | leptotene | circular , centric | 1 | 0 | no |
| mnd1 | 6 | fil3 | 0/30 | 1 | leptotene | circular , centric | 1 | 0 | no |
| mnd1 | 6 | ant4, fil1 | 0/30 | 1 | leptotene | circular , centric | 1 | 0 | no |
| mnd1 | 6 | fil 2 | 0/30 | 1 | leptotene | circular , centric | 1 | 0 | no |
| mnd1 | 6 | fil3 | 0/30 | 1 | leptotene | circular , centric | 1 | 0 | no |
| mnd1 | 6 | **bud 4**, ant1, fil1 | 0/30 | 1 | zygotene | not visible | 1 | 76-99 | no |
| mnd1 | 6 | fil2 | 0/30 | 1 | zygotene | not visible | 1 | 76-99 | no |
| mnd1 | 6 | fil3 | 0/30 | 1 | zygotene | not visible | 1 | 76-99 | no |
| mnd1 | 6 | ant2, fil1 | 0/30 | 1 | zygotene | not visible | 1 | 76-99 | no |
| mnd1 | 6 | fil 2 | 0/30 | 1 | zygotene | not visible | 1 | 76-99 | no |

| | | | | | | | | | |
|---|---|---|---|---|---|---|---|---|---|
| mnd1 | 6 | fil3 | 0/30 | 1 | zygotene | not visible | 1 | 76-99 | no |
| mnd1 | 6 | ant3, fil1 | 0/30 | 1 | zygotene | not visible | 1 | 76-99 | no |
| mnd1 | 6 | fil 2 | 0/30 | 1 | zygotene | not visible | 1 | 76-99 | no |
| mnd1 | 6 | fil3 | 0/30 | 1 | zygotene | not visible | 1 | 76-99 | no |
| mnd1 | 6 | **bud 5**, ant1, fil1 | 0/30 | 1 | zygotene | not visible | 1 | 76-99 | no |
| mnd1 | 6 | fil 2 | 0/30 | 1 | zygotene | not visible | 1 | 76-99 | no |
| mnd1 | 6 | fil3 | 0/30 | 1 | zygotene | not visible | 1 | 76-99 | no |
| mnd1 | 6 | ant2, fil1 | 0/30 | 1 | zygotene | not visible | 1 | 76-99 | no |
| mnd1 | 6 | fil 2 | 0/30 | 1 | zygotene | not visible | 1 | 76-99 | no |
| mnd1 | 6 | fil3 | 0/30 | 1 | zygotene | not visible | 1 | 76-99 | no |
| mnd1 | 6 | **bud 6** ant1, fil1 | 0/30 | 2 | zygotene | not visible | 1 | 26-75 | no |
| mnd1 | 6 | fil 2 | 0/30 | 2 | zygotene | not visible | 1 | 26-75 | no |
| mnd1 | 6 | fil3 | 0/30 | 2 | zygotene | not visible | 1 | 26-75 | no |
| mnd1 | 6 | ant2, fil1 | 0/30 | 1 | zygotene | not visible | 1 | 76-99 | no |
| mnd1 | 6 | fil 2 | 0/30 | 1 | zygotene | not visible | 1 | 76-99 | no |
| mnd1 | 6 | fil3 | 0/30 | 1 | zygotene | not visible | 1 | 76-99 | no |
| mnd1 | 6 | **infl 7**, ant1, fil1 | 0/30 | 1 | zygotene | not visible | 1 | 76-99 | no |
| mnd1 | 6 | fil 2 | 0/30 | 1 | zygotene | not visible | 1 | 76-99 | no |
| mnd1 | 6 | fil3 | 0/30 | 1 | zygotene | not visible | 1 | 76-99 | no |
| mnd1 | 6 | **infl 8**, ant1, fil1 | 0/30 | 2 | zygotene | not visible | 1 | 1 to 25 | no |
| mnd1 | 6 | fil2 | 0/30 | 2 | zygotene | not visible | 1 | 1 to 25 | no |
| mnd1 | 6 | fil3 | 0/30 | 2 | zygotene | not visible | 1 | 1 to 25 | no |
| mnd1 | 6 | ant2, fil1 | 0/30 | 0 | eP | circular,peripheral | 1 | 100 | no |
| mnd1 | 6 | fil 2 | 0/30 | 0 | eP | circular,peripheral | 1 | 100 | no |
| mnd1 | 6 | fil3 | 0/30 | 0 | eP | circular,peripheral | 1 | 100 | no |
| mnd1 | 6 | **bud 9** ant1, fil1 | 0/30 | 1 | eP | circular,peripheral | 1 | 100 | no |
| mnd1 | 6 | fil2 | 0/30 | 1 | eP | circular,peripheral | 1 | 100 | no |
| mnd1 | 6 | fil3 | 0/30 | 1 | eP | circular,peripheral | 1 | 100 | no |

| | | | | | | | | | |
|---|---|---|---|---|---|---|---|---|---|
| mnd1 | 6 | ant2, fil1 | 0/30 | 1 | eP | circular,peripheral | 1 | 100 | no |
| mnd1 | 6 | fil 2 | 0/30 | 1 | eP | circular,peripheral | 1 | 100 | no |
| mnd1 | 6 | fil3 | 0/30 | 1 | eP | circular,peripheral | 1 | 100 | no |
| mnd1 | 6 | ant3, fil1 | 0/30 | 0 | eP | circular,peripheral | 1 | 100 | no |
| mnd1 | 6 | fil 2 | 0/30 | 0 | eP | circular,peripheral | 1 | 100 | no |
| mnd1 | 6 | fil3 | 0/30 | 0 | eP | circular,peripheral | 1 | 100 | no |
| mnd1 | 6 | **bud 10** ant1, fil1 | 0/30 | 1 | eP | circular,peripheral | 1 | 100 | no |
| mnd1 | 6 | fil 2 | 0/30 | 1 | eP | circular,peripheral | 1 | 100 | no |
| mnd1 | 6 | fil3 | 0/30 | 1 | eP | circular,peripheral | 1 | 100 | no |
| mnd1 | 6 | ant2, fil1 | 0/30 | 1 | eP | circular,peripheral | 1 | 100 | no |
| mnd1 | 6 | fil 2 | 0/30 | 1 | eP | circular,peripheral | 1 | 100 | no |
| mnd1 | 6 | fil3 | 0/30 | 1 | eP | circular,peripheral | 1 | 100 | no |
| mnd1 | 6 | ant3, fil1 | 0/30 | 1 | eP | circular,peripheral | 1 | 100 | no |
| mnd1 | 6 | fil 2 | 0/30 | 1 | eP | circular,peripheral | 1 | 100 | no |
| mnd1 | 6 | fil3 | 0/30 | 1 | eP | circular,peripheral | 1 | 100 | no |
| mnd1 | 6 | bud **11** ant1, fil1 | 0/30 | 2 | zygotene | not visible | 1 | 2575 | no |
| mnd1 | 6 | fil 2 | 0/30 | 2 | zygotene | not visible | 1 | 26-75 | no |
| mnd1 | 6 | fil3 | 0/30 | 2 | zygotene | not visible | 1 | 26-75 | no |
| mnd1 | 6 | ant2, fil1 | 0/30 | 2 | zygotene | not visible | 1 | 2575 | no |
| mnd1 | 6 | fil 2 | 0/30 | 2 | zygotene | not visible | 1 | 26-75 | no |
| mnd1 | 6 | fil3 | 0/30 | 2 | zygotene | not visible | 1 | 26-75 | no |
| mnd1 | 6 | ant3, fil1 | 0/30 | 1 | zygotene | not visible | 1 | 76-99 | no |
| mnd1 | 6 | fil 2 | 0/30 | 1 | zygotene | not visible | 1 | 76-99 | no |
| mnd1 | 6 | fil3 | 0/30 | 1 | zygotene | not visible | 1 | 76-99 | no |
| mnd1 | 8 | **bud 1**, ant1, fil1 | 0/30 | 1 | eP | circular,peripheral | 1 | 100 | no |
| mnd1 | 8 | fil 2 | 0/30 | 1 | eP | circular,peripheral | 1 | 100 | no |
| mnd1 | 8 | fil3 | 0/30 | 1 | eP | circular,peripheral | 1 | 100 | no |

| | | | | | | | | | |
|---|---|---|---|---|---|---|---|---|---|
| mnd1 | 8 | ant 2, fil 1 | 0/30 | 1 | eP | circular,peripheral | 1 | 100 | no |
| mnd1 | 8 | fil2 | 0/30 | 1 | eP | circular,peripheral | 1 | 100 | no |
| mnd1 | 8 | fil3 | 0/30 | 1 | eP | circular,peripheral | 1 | 100 | no |
| mnd1 | 8 | ant 3 fil1 | 0/30 | 1 | eP | circular,peripheral | 1 | 100 | no |
| mnd1 | 8 | | 0/30 | 1 | eP | circular,peripheral | 1 | 100 | no |
| mnd1 | 8 | | 0/30 | 1 | eP | circular,peripheral | 1 | 100 | no |
| mnd1 | 8 | **bud 2** ant1, fil1 | 0/30 | 0 | m-l P | circular,peripheral | 3 | 100 | no |
| mnd1 | 8 | fil 2 | 0/30 | 0 | m-l P | circular,peripheral | 3 | 100 | no |
| mnd1 | 8 | fil3 | 0/30 | 0 | m-l P | circular,peripheral | 3 | 100 | no |
| mnd1 | 8 | ant2, fil1 | 0/30 | 0 | m-l P | circular,peripheral | 3 | 100 | no |
| mnd1 | 8 | fil 2 | 0/30 | 0 | m-l P | circular,peripheral | 3 | 100 | no |
| mnd1 | 8 | fil3 | 0/30 | 0 | m-l P | circular,peripheral | 3 | 100 | no |
| mnd1 | 8 | **bud 3**, ant1, fil1 | 0/30 | 0 | tetrad | circular , centric | 3 | 100 | no |
| mnd1 | 8 | fil 2 | 0/30 | 0 | tetrad | circular , centric | 3 | 100 | no |
| mnd1 | 8 | fil3 | 0/30 | 0 | tetrad | circular , centric | 3 | 100 | no |
| mnd1 | 8 | ant 2, fil 1 | 0/30 | 0 | anaphase 2 | not visible | 3 | 100 | no |
| mnd1 | 8 | fil2 | 0/30 | 0 | anaphase 2 | not visible | 3 | 100 | no |
| mnd1 | 8 | fil3 | 0/30 | 0 | anaphase 2 | not visible | 3 | 100 | no |
| mnd1 | 8 | ant 3 fil1 | 0/30 | 0 | diakenesis | not visible | 3 | 100 | no |
| mnd1 | 8 | fil 2 | 0/30 | 0 | diakenesis | not visible | 3 | 100 | no |
| mnd1 | 8 | fil 3 | 0/30 | 0 | diakenesis | not visible | 3 | 100 | no |
| mnd1 | 8 | ant 4 fil 1 | 0/30 | 0 | tetrad | circular , centric | 3 | 100 | no |
| mnd1 | 8 | fil2 | 0/30 | 0 | tetrad | circular , centric | 3 | 100 | no |
| mnd1 | 8 | fil3 | 0/30 | 0 | tetrad | circular , centric | 3 | 100 | no |
| mnd1 | 8 | **bud 4**, ant1, fil1 | 0/30 | 0 | leptotene | circular , centric | 1 | 0 | no |
| mnd1 | 8 | fil 2 | 0/30 | 0 | leptotene | circular , centric | 1 | 0 | no |
| mnd1 | 8 | fil3 | 0/30 | 0 | leptotene | circular , centric | 1 | 0 | no |
| mnd1 | 8 | bud 5, ant1, fil1 | 0/30 | 0 | tetrad | circular , centric | N/A | 100 | no |

| | | | | | | | | | |
|---|---|---|---|---|---|---|---|---|---|
| mnd1 | 8 | fil 2 | 0/30 | 0 | tetrad | circular , centric | N/A | 100 | no |
| mnd1 | 8 | fil3 | 0/30 | 0 | tetrad | circular , centric | N/A | 100 | no |
| mnd1 | 8 | **bud 6**, ant1, fil1 | 0/30 | 0 | diplotene | not visible | N/A | 100 | no |
| mnd1 | 8 | fil 2 | 0/30 | 0 | diplotene | not visible | N/A | 100 | no |
| mnd1 | 8 | fil3 | 0/30 | 0 | diakenesis | not visible | N/A | 100 | no |
| mnd1 | 8 | fil 4 | 0/30 | 0 | anaphase 1 | not visible | N/A | 100 | no |
| mnd1 | 8 | ant 2, fil 1 | 0/30 | 0 | m-l P | circular,peripheral | 2 | 100 | no |
| mnd1 | 8 | fil2 | 0/30 | 0 | m-l P | circular,peripheral | 2 | 100 | no |
| mnd1 | 8 | fil3 | 0/30 | 0 | m-l P | circular,peripheral | 2 | 100 | no |
| mnd1 | 8 | fil 4 | 0/30 | 0 | m-l P | circular,peripheral | 2 | 100 | no |
| mnd1 | 8 | ant 3 fil1 | 0/30 | 0 | m-l P | circular,peripheral | 2 | 100 | no |
| mnd1 | 8 | fil 2 | 0/30 | 0 | m-l P | circular,peripheral | 2 | 100 | no |
| mnd1 | 8 | fil 3 | 0/30 | 0 | m-l P | circular,peripheral | 2 | 100 | no |
| mnd1 | 8 | **bud 7** ant1, fil1 | 0/30 | 0 | m-l P | circular,peripheral | 2 | 100 | no |
| mnd1 | 8 | fil 2 | 0/30 | 0 | m-l P | circular,peripheral | 2 | 100 | no |
| mnd1 | 8 | fil3 | 0/30 | 0 | m-l P | circular,peripheral | 2 | 100 | no |
| mnd1 | 8 | ant2, fil1 | 0/30 | 0 | eP | circular,peripheral | 1 | 100 | no |
| mnd1 | 8 | fil 2 | 0/30 | 0 | eP | circular,peripheral | 1 | 100 | no |
| mnd1 | 8 | fil3 | 0/30 | 0 | eP | circular,peripheral | 1 | 100 | no |
| mnd1 | 8 | **bud 8**, ant1, fil1 | 0/30 | 2 | zygotene | not visible | 1 | 1 to 25 | no |
| mnd1 | 8 | fil 2 | 0/30 | 2 | zygotene | not visible | 1 | 1 to 25 | no |
| mnd1 | 8 | fil3 | 0/30 | 2 | zygotene | not visible | 1 | 1 to 25 | no |
| mnd1 | 8 | ant 2, fil 1 | 0/30 | 1 | zygotene | not visible | 1 | 26-75 | no |
| mnd1 | 8 | fil2 | 0/30 | 1 | zygotene | not visible | 1 | 26-75 | no |
| mnd1 | 8 | fil3 | 0/30 | 1 | zygotene | not visible | 1 | 26-75 | no |
| mnd1 | 8 | ant 3 fil1 | 0/30 | 1 | zygotene | not visible | 1 | 26-75 | no |
| mnd1 | 8 | fil 2 | 0/30 | 1 | zygotene | not visible | 1 | 26-75 | no |
| mnd1 | 8 | fil 3 | 0/30 | 1 | zygotene | not visible | 1 | 26-75 | no |

| | | | | | | | | | |
|---|---|---|---|---|---|---|---|---|---|
| mnd1 | 8 | **bud 9**, ant1, fil1 | 5/30 | 0 | leptotene | circular , centric | 1 | 0 | no |
| mnd1 | 8 | fil 2 | 7/30 | 0 | leptotene | circular , centric | 1 | 0 | no |
| mnd1 | 8 | fil3 | 10/30 | 0 | leptotene | circular , centric | 1 | 0 | no |
| mnd1 | 8 | ant 2, fil 1 | 0/30 | 0 | eP | circular,peripheral | 1 | 100 | no |
| mnd1 | 8 | fil2 | 0/30 | 0 | eP | circular,peripheral | 1 | 100 | no |
| mnd1 | 8 | fil3 | 0/30 | 0 | eP | circular,peripheral | 1 | 100 | no |
| mnd1 | 8 | ant 3 fil1 | 0/30 | 1 | zygotene | not visible | 1 | 1 to 25 | no |
| mnd1 | 8 | fil 2 | 0/30 | 1 | zygotene | not visible | 1 | 1 to 25 | no |
| mnd1 | 8 | fil 3 | 0/30 | 1 | zygotene | not visible | 1 | 1 to 25 | no |
| mnd1 | 8 | ant 4 fil 1 | 0/30 | 0 | leptotene | circular , centric | 1 | 0 | no |
| mnd1 | 8 | fil2 | 0/30 | 0 | leptotene | circular , centric | 1 | 0 | no |
| mnd1 | 8 | fil3 | 0/30 | 0 | leptotene | circular , centric | 1 | 0 | no |
| mnd1 | 8 | **bud 10**, ant1, fil1 | 0/30 | 0 | eP | circular,peripheral | 1 | 100 | no |
| mnd1 | 8 | fil 2 | 0/30 | 0 | eP | circular,peripheral | 1 | 100 | no |
| mnd1 | 8 | fil3 | 0/30 | 0 | eP | circular,peripheral | 1 | 100 | no |
| mnd1 | 10 | **bud 1**, ant1, fil1 | 30/30 | 1 | leptotene | circular , centric | 1 | 0 | no |
| mnd1 | 10 | fil 2 | 30/30 | 1 | leptotene | circular , centric | 1 | 0 | no |
| mnd1 | 10 | fil3 | 20/30 | 1 | leptotene | circular , centric | 1 | 0 | no |
| mnd1 | 10 | ant 2, fil 1 | 0/30 | 0 | leptotene | circular , centric | 1 | 0 | no |
| mnd1 | 10 | fil2 | 0/30 | 0 | leptotene | circular , centric | 1 | 0 | no |
| mnd1 | 10 | fil3 | 0/30 | 0 | leptotene | circular , centric | 1 | 0 | no |
| mnd1 | 10 | ant 3 fil1 | 30/30 | 1 | leptotene | circular , centric | 1 | 0 | no |
| mnd1 | 10 | fil 2 | 25/30 | 1 | leptotene | circular , centric | 1 | 0 | no |
| mnd1 | 10 | fil 3 | 30/30 | 1 | leptotene | circular , centric | 1 | 0 | no |
| mnd1 | 10 | ant 4 fil 1 | 30/30 | 1 | leptotene | circular , centric | 1 | 0 | no |
| mnd1 | 10 | fil2 | 30/30 | 1 | leptotene | circular , centric | 1 | 0 | no |
| **mnd1** | **10** | **bud 2, ant1, fil1** | **30/30** | **1** | **zygotene** | **not visible** | **1** | **1 to 25** | **yes** |
| **mnd1** | **10** | **fil 2** | **30/30** | **1** | **zygotene** | **not visible** | **1** | **1 to 25** | **yes** |

| | | | | | | | | | |
|---|---|---|---|---|---|---|---|---|---|
| **mnd1** | **10** | **fil3** | **30/30** | **1** | **zygotene** | **not visible** | **1** | **1 to 25** | **yes** |
| **mnd1** | **10** | **ant 2, fil 1** | **30/30** | **1** | **zygotene** | **not visible** | **1** | **1 to 25** | **yes** |
| **mnd1** | **10** | **fil2** | **30/30** | **1** | **zygotene** | **not visible** | **1** | **1 to 25** | **yes** |
| **mnd1** | **10** | **fil3** | **30/30** | **1** | **zygotene** | **not visible** | **1** | **1 to 25** | **yes** |
| mnd1 | 10 | ant 3 fil1 | 30/30 | 1 | leptotene | circular , centric | 1 | 0 | no |
| mnd1 | 10 | fil 2 | 30/30 | 1 | leptotene | circular , centric | 1 | 0 | no |
| mnd1 | 10 | fil 3 | 30/30 | 1 | leptotene | circular , centric | 1 | 0 | no |
| mnd1 | 10 | bud 3, ant1, fil1 | 0/30 | 0 | m-l P | circular,peripheral | 3 | 100 | no |
| mnd1 | 10 | fil 2 | 0/30 | 0 | m-l P | circular,peripheral | 3 | 100 | no |
| mnd1 | 10 | fil3 | 0/30 | 0 | m-l P | circular,peripheral | 3 | 100 | no |
| mnd1 | 10 | ant 2, fil 1 | 0/30 | 0 | m-l P | circular,peripheral | 3 | 100 | no |
| mnd1 | 10 | fil2 | 0/30 | 0 | m-l P | circular,peripheral | 3 | 100 | no |
| mnd1 | 10 | fil3 | 0/30 | 0 | m-l P | circular,peripheral | 3 | 100 | no |
| mnd1 | 10 | ant 3 fil1 | 0/30 | 0 | tetrad | circular , centric | 3 | 100 | no |
| mnd1 | 10 | fil 2 | 0/30 | 0 | tetrad | circular , centric | 3 | 100 | no |
| mnd1 | 10 | fil 3 | 0/30 | 0 | tetrad | circular , centric | 3 | 100 | no |
| mnd1 | 10 | bud 4, ant1, fil1 | 0/30 | 1 | eP | circular,peripheral | 1 | 100 | no |
| mnd1 | 10 | fil 2 | 0/30 | 1 | eP | circular,peripheral | 1 | 100 | no |
| mnd1 | 10 | fil3 | 0/30 | 0 | m-l P | circular,peripheral | 3 | 100 | no |
| mnd1 | 10 | ant 2, fil 1 | 5/30 | 1 | leptotene | circular , pericentric | 1 | 0 | no |
| mnd1 | 10 | fil2 | 0/30 | 1 | leptotene | circular , pericentric | 1 | 0 | no |
| mnd1 | 10 | fil3 | 3/30 | 1 | leptotene | circular , pericentric | 1 | 0 | no |
| mnd1 | 10 | ant 3 fil1 | 3/30 | 1 | leptotene | circular , pericentric | 1 | 0 | no |
| mnd1 | 10 | fil 2 | 5/30 | 1 | leptotene | circular , pericentric | 1 | 0 | no |
| mnd1 | 10 | fil 3 | 10/30 | 1 | leptotene | circular , pericentric | 1 | 0 | no |
| mnd1 | 10 | bud 5, ant1, fil1 | 0/30 | 0 | m-l P | circular,peripheral | 2 | 100 | no |
| mnd1 | 10 | fil 2 | 0/30 | 0 | m-l P | circular,peripheral | 2 | 100 | no |
| mnd1 | 10 | fil3 | 0/30 | 0 | m-l P | circular,peripheral | 2 | 100 | no |
| mnd1 | 10 | ant 2, fil 1 | 5/30 | 2 | zygotene | not visible | 1 | 1 to 25 | no |

| | | | | | | | | | |
|---|---|---|---|---|---|---|---|---|---|
| mnd1 | 10 | fil2 | 5/30 | 2 | zygotene | not visible | 1 | 1 to 25 | no |
| mnd1 | 10 | fil3 | 3/30 | 2 | zygotene | not visible | 1 | 1 to 25 | no |
| mnd1 | 10 | bud 6, ant1, fil1 | 20/30 | 2 | zygotene | not visible | 1 | 1 to 25 | yes |
| mnd1 | 10 | fil 2 | 30/30 | 2 | zygotene | not visible | 1 | 1 to 25 | yes |
| mnd1 | 10 | fil3 | 30/30 | 2 | zygotene | not visible | 1 | 1 to 25 | yes |
| mnd1 | 10 | ant 2, fil 1 | 30/30 | 2 | zygotene | not visible | 1 | 1 to 25 | yes |
| mnd1 | 10 | fil2 | 20/30 | 2 | zygotene | not visible | 1 | 1 to 25 | yes |
| mnd1 | 10 | fil3 | 30/30 | 2 | zygotene | not visible | 1 | 1 to 25 | yes |
| mnd1 | 12 | **bud 1**, ant1, fil1 | 0/30 | 0 | anaphase II | not visible | N/A | 100 | yes |
| mnd1 | 12 | fil 2 | 0/30 | 0 | anaphase II | not visible | N/A | 100 | yes |
| mnd1 | 12 | fil3 | 0/30 | 0 | anaphase II | not visible | N/A | 100 | yes |
| mnd1 | 12 | ant 2, fil 1 | 0/30 | 0 | anaphase I | not visible | N/A | 100 | yes |
| mnd1 | 12 | fil2 | 0/30 | 0 | anaphase I | not visible | N/A | 100 | yes |
| mnd1 | 12 | fil3 | 0/30 | 0 | anaphase I | not visible | N/A | 100 | yes |
| mnd1 | 12 | ant 3 fil1 | 0/30 | 0 | metaphase I | not visible | N/A | 100 | yes |
| mnd1 | 12 | fil 2 | 0/30 | 0 | metaphase I | not visible | N/A | 100 | yes |
| mnd1 | 12 | fil 3 | 0/30 | 0 | metaphase I | not visible | N/A | 100 | yes |
| mnd1 | 12 | ant 4 fil 1 | 0/30 | 0 | m-l P | circular,peripheral | 3 | 100 | no |
| mnd1 | 12 | fil2 | 0/30 | 0 | m-l P | circular,peripheral | 3 | 100 | no |
| mnd1 | 12 | fil3 | 0/30 | 0 | m-l P | circular,peripheral | 3 | 100 | no |
| mnd1 | 12 | **bud 2**, ant1, fil1 | 30/30 | 2 | zygotene | not visible | 1 | 26-75 | yes |
| mnd1 | 12 | fil 2 | 30/30 | 2 | zygotene | not visible | 1 | 26-75 | yes |
| mnd1 | 12 | fil3 | 30/30 | 2 | zygotene | not visible | 1 | 26-75 | yes |
| mnd1 | 12 | ant 2, fil 1 | 0/30 | 0 | m-l P | circular,peripheral | 3 | 100 | no |
| mnd1 | 12 | fil2 | 0/30 | 0 | m-l P | circular,peripheral | 3 | 100 | no |
| mnd1 | 12 | fil3 | 0/30 | 0 | m-l P | circular,peripheral | 3 | 100 | no |
| mnd1 | 12 | ant 3 fil1 | 30/30 | 2 | zygotene | not visible | 1 | 2575 | yes |
| mnd1 | 12 | fil 2 | 30/30 | 2 | zygotene | not visible | 1 | 26-75 | yes |
| mnd1 | 12 | fil 3 | 30/30 | 2 | zygotene | not visible | 1 | 26-75 | yes |

| | | | | | | | | | |
|---|---|---|---|---|---|---|---|---|---|
| mnd1 | 12 | ant 4 fil 1 | 30/30 | 2 | zygotene | not visible | 1 | 1 to 25 | yes |
| mnd1 | 12 | fil2 | 30/30 | 2 | zygotene | not visible | 1 | 1 to 25 | yes |
| mnd1 | 12 | fil3 | 30/30 | 2 | zygotene | not visible | 1 | 1 to 25 | yes |
| mnd1 | 12 | **bud 3**, ant1, fil1 | 30/30 | 1 | zygotene | not visible | 1 | 76-99 | yes |
| mnd1 | 12 | fil 2 | 30/30 | 1 | zygotene | not visible | 1 | 76-99 | yes |
| mnd1 | 12 | fil3 | 30/30 | 1 | zygotene | not visible | 1 | 76-99 | yes |
| mnd1 | 12 | ant 2, fil 1 | 30/30 | 2 | zygotene | not visible | 1 | 1 to 25 | yes |
| mnd1 | 12 | fil2 | 30/30 | 2 | zygotene | not visible | 1 | 1 to 25 | yes |
| mnd1 | 12 | fil3 | 30/30 | 2 | zygotene | not visible | 1 | 1 to 25 | yes |
| mnd1 | 12 | **bud 4**, ant1, fil1 | 0/30 | 1 | eP | circular,peripheral | 1 | 100 | no |
| mnd1 | 12 | fil 2 | 0/30 | 1 | eP | circular,peripheral | 1 | 100 | no |
| mnd1 | 12 | fil3 | 0/30 | 1 | eP | circular,peripheral | 1 | 100 | no |
| mnd1 | 12 | ant 2, fil 1 | 0/30 | 0 | m-l P | circular,peripheral | 2 | 100 | no |
| mnd1 | 12 | fil2 | 0/30 | 0 | m-l P | circular,peripheral | 2 | 100 | no |
| mnd1 | 12 | **bud 5**, ant1, fil1 | 0/30 | 0 | tetrad | circular , centric | 3 | 100 | no |
| mnd1 | 14 | **bud 1**, ant1, fil1 | 30/30 | 1 | zygotene | not visible | 1 | 76-99 | yes |
| mnd1 | 14 | fil 2 | 30/30 | 1 | zygotene | not visible | 1 | 76-99 | yes |
| mnd1 | 14 | fil3 | 30/30 | 1 | zygotene | not visible | 1 | 76-99 | yes |
| mnd1 | 14 | ant 2, fil 1 | 10/30 | 1 | zygotene | not visible | 1 | 76-99 | yes |
| mnd1 | 14 | fil2 | 15/30 | 1 | zygotene | not visible | 1 | 76-99 | yes |
| mnd1 | 14 | fil3 | 15/30 | 1 | zygotene | not visible | 1 | 76-99 | yes |
| mnd1 | 14 | ant 3 fil1 | 0/30 | 0 | metaphase I | not visible | N/A | 100 | yes |
| mnd1 | 14 | fil 2 | 0/30 | 0 | metaphase I | not visible | N/A | 100 | yes |
| mnd1 | 14 | fil 3 | 0/30 | 0 | metaphase I | not visible | N/A | 100 | yes |
| mnd1 | 14 | **bud 2** ant 1 fil1 | 0/30 | 0 | metaphase I | not visible | N/A | 100 | yes |
| mnd1 | 16 | **bud 1**, ant1, fil1 | 30/30 | 1 | leptotene | circular , pericentric | 1 | 0 | yes |
| mnd1 | 16 | fil 2 | 30/30 | 1 | leptotene | circular , pericentric | 1 | 0 | yes |
| mnd1 | 16 | fil3 | 30/30 | 1 | leptotene | circular , pericentric | 1 | 0 | yes |

| | | | | | | | | | |
|---|---|---|---|---|---|---|---|---|---|
| mnd1 | 16 | ant 2, fil 1 | 30/30 | 2 | zygotene | not visible | 1 | 1 to 25 | yes |
| mnd1 | 16 | fil2 | 30/30 | 2 | zygotene | not visible | 1 | 1 to 25 | yes |
| mnd1 | 16 | fil3 | 30/30 | 2 | zygotene | not visible | 1 | 1 to 25 | yes |
| mnd1 | 16 | ant 3 fil1 | 0/30 | 2 | zygotene | not visible | 1 | 2575 | no |
| mnd1 | 16 | fil 2 | 0/30 | 2 | zygotene | not visible | 1 | 26-75 | no |
| mnd1 | 16 | ant 4 fil 1 | 0/30 | 0 | m-l P | circular,peripheral | 2 | 100 | no |
| mnd1 | 16 | fil2 | 0/30 | 0 | m-l P | circular,peripheral | 2 | 100 | no |
| mnd1 | 16 | fil3 | 0/30 | 0 | m-l P | circular,peripheral | 2 | 100 | no |
| mnd1 | 16 | ant 5 fil 1 | 0/30 | 0 | m-l P | circular,peripheral | 2 | 100 | no |
| mnd1 | 16 | fil2 | 0/30 | 0 | m-l P | circular,peripheral | 2 | 100 | no |
| mnd1 | 16 | fil3 | 0/30 | 0 | m-l P | circular,peripheral | 2 | 100 | no |
| mnd1 | 16 | **bud 2**, ant1, fil1 | 0/30 | 0 | m-l P | circular,peripheral | 3 | 100 | no |
| mnd1 | 16 | fil 2 | 0/30 | 0 | m-l P | circular,peripheral | 3 | 100 | no |
| mnd1 | 16 | fil3 | 0/30 | 0 | m-l P | circular,peripheral | 3 | 100 | no |
| mnd1 | 16 | **bud 3**, ant1, fil1 | 0/30 | 0 | tetrad | circular , centric | 3 | 100 | no |
| mnd1 | 16 | fil 2 | 0/30 | 0 | tetrad | circular , centric | 3 | 100 | no |
| mnd1 | 16 | fil3 | 0/30 | 0 | tetrad | circular , centric | 3 | 100 | no |
| mnd1 | 16 | **bud 4**, ant1, fil1 | 30/30 | 1 | leptotene | circular , centric | 1 | 0 | no |
| mnd1 | 16 | fil 2 | 25/30 | 1 | leptotene | circular , centric | 1 | 0 | no |
| mnd1 | 16 | fil3 | 30/30 | 1 | leptotene | circular , centric | 1 | 0 | no |
| mnd1 | 16 | **bud 5**, ant1, fil1 | 0/30 | 1 | zygotene | not visible | 1 | 76-99 | no |
| mnd1 | 16 | fil 2 | 0/30 | 1 | zygotene | not visible | 1 | 76-99 | no |
| mnd1 | 16 | fil3 | 0/30 | 1 | zygotene | not visible | 1 | 76-99 | no |
| mnd1 | 16 | **bud 6** ant 1, fil 1 | 0/30 | 0 | m-l P | circular,peripheral | 2 | 100 | no |
| mnd1 | 16 | fil2 | 0/30 | 0 | m-l P | circular,peripheral | 2 | 100 | no |
| mnd1 | 16 | fil3 | 0/30 | 0 | m-l P | circular,peripheral | 2 | 100 | no |
| mnd1 | 16 | ant 2 fil1 | 0/30 | 0 | m-l P | circular,peripheral | 2 | 100 | no |
| mnd1 | 16 | fil 2 | 0/30 | 0 | m-l P | circular,peripheral | 2 | 100 | no |

| | | | | | | | | |
|---|---|---|---|---|---|---|---|---|
| mnd1 | 16 | | 0/30 | 0 | m-l P | circular,peripheral | 2 | 100 | no |
| mnd1 | 16 | ant 3 fil 1 | 0/30 | 0 | m-l P | circular,peripheral | 2 | 100 | no |
| mnd1 | 16 | fil2 | 0/30 | 0 | m-l P | circular,peripheral | 2 | 100 | no |
| mnd1 | 16 | fil3 | 0/30 | 0 | m-l P | circular,peripheral | 2 | 100 | no |
| mnd1 | 16 | ant 4 fil 1 | 0/30 | 0 | eP | circular,peripheral | 1 | 100 | no |
| mnd1 | 16 | fil2 | 0/30 | 0 | eP | circular,peripheral | 1 | 100 | no |
| mnd1 | 16 | fil3 | 0/30 | 0 | eP | circular,peripheral | 1 | 100 | no |
| mnd1 | 16 | **bud 7**, ant1, fil1 | 0/30 | 1 | eP | circular,peripheral | 1 | 100 | no |
| mnd1 | 16 | fil 2 | 0/30 | 1 | eP | circular,peripheral | 1 | 100 | no |
| mnd1 | 16 | fil3 | 0/30 | 1 | eP | circular,peripheral | 1 | 100 | no |
| mnd1 | 16 | ant 2, fil 1 | 0/30 | 1 | eP | circular,peripheral | 1 | 100 | no |
| mnd1 | 16 | fil2 | 0/30 | 1 | eP | circular,peripheral | 1 | 100 | no |
| mnd1 | 16 | fil3 | 0/30 | 1 | eP | circular,peripheral | 1 | 100 | no |
| mnd1 | 16 | **bud 8**, ant1, fil1 | 30/30 | 1 | leptotene | circular , pericentric | 1 | 0 | yes |
| mnd1 | 16 | fil 2 | 30/30 | 1 | leptotene | circular , pericentric | 1 | 0 | yes |
| mnd1 | 16 | fil3 | 30/30 | 1 | leptotene | circular , pericentric | 1 | 0 | yes |
| mnd1 | 16 | ant 2, fil 1 | 30/30 | 1 | leptotene | circular , pericentric | 1 | 0 | yes |
| mnd1 | 16 | fil2 | 30/30 | 1 | leptotene | circular , pericentric | 1 | 0 | yes |
| mnd1 | 16 | fil3 | 30/30 | 1 | leptotene | circular , pericentric | 1 | 0 | yes |
| mnd1 | 16 | **bud 9**, ant1, fil1 | 30/30 | 2 | zygotene | not visible | 1 | 1 to 25 | yes |
| mnd1 | 16 | fil 2 | 30/30 | 2 | zygotene | not visible | 1 | 1 to 25 | yes |
| mnd1 | 16 | fil3 | 30/30 | 2 | zygotene | not visible | 1 | 1 to 25 | yes |
| mnd1 | 16 | ant 2, fil 1 | 30/30 | 2 | zygotene | not visible | 1 | 1 to 25 | yes |
| mnd1 | 16 | fil2 | 30/30 | 2 | zygotene | not visible | 1 | 1 to 25 | yes |
| mnd1 | 16 | fil3 | 30/30 | 2 | zygotene | not visible | 1 | 1 to 25 | yes |
| mnd1 | 16 | ant 3 fil1 | 30/30 | 2 | zygotene | not visible | 1 | 1 to 25 | yes |
| mnd1 | 16 | fil 2 | 30/30 | 2 | zygotene | not visible | 1 | 1 to 25 | yes |
| mnd1 | 16 | fil 3 | 30/30 | 2 | zygotene | not visible | 1 | 1 to 25 | yes |
| mnd1 | 16 | fil 4 | 30/30 | 2 | zygotene | not visible | 1 | 1 to 25 | yes |

| | | | | | | | | | |
|---|---|---|---|---|---|---|---|---|---|
| mnd1 | 18 | **bud 1**, ant1, fil1 | 0/30 | 0 | G2 | circular , centric | 1 | 0 | no |
| mnd1 | 18 | fil 2 | 0/30 | 0 | G2 | circular , centric | 1 | 0 | no |
| mnd1 | 18 | fil3 | 0/30 | 0 | G2 | circular , centric | 1 | 0 | no |
| mnd1 | 18 | **bud 2**, ant1, fil1 | 0/30 | 0 | tetrad | circular , centric | 3 | 100 | no |
| mnd1 | 18 | fil 2 | 0/30 | 0 | tetrad | circular , centric | 3 | 100 | no |
| mnd1 | 18 | fil3 | 0/30 | 0 | tetrad | circular , centric | 3 | 100 | no |
| mnd1 | 18 | **bud 3**, ant1, fil1 | 30/30 | 1 | zygotene | not visible | 1 | 26-75 | yes |
| mnd1 | 18 | fil 2 | 30/30 | 1 | zygotene | not visible | 1 | 26-75 | yes |
| mnd1 | 18 | fil3 | 30/30 | 1 | zygotene | not visible | 1 | 26-75 | yes |
| mnd1 | 18 | **bud 4**, ant1, fil1 | 30/30 | 1 | leptotene | circular , pericentric | 1 | 0 | yes |
| mnd1 | 18 | fil 2 | 30/30 | 1 | leptotene | circular , pericentric | 1 | 0 | yes |
| mnd1 | 18 | fil3 | 30/30 | 1 | leptotene | circular , pericentric | 1 | 0 | yes |
| mnd1 | 18 | ant 2, fil 1 | 30/30 | 1 | leptotene | circular , pericentric | 1 | 0 | yes |
| mnd1 | 18 | fil2 | 30/30 | 1 | leptotene | circular , pericentric | 1 | 0 | yes |
| mnd1 | 18 | fil3 | 30/30 | 1 | leptotene | circular , pericentric | 1 | 0 | yes |
| mnd1 | 18 | **bud 5**, ant1, fil1 | 30/30 | 2 | leptotene | circular , peripheral | 1 | 0 | yes |
| mnd1 | 18 | fil 2 | 30/30 | 2 | leptotene | circular , peripheral | 1 | 0 | yes |
| mnd1 | 18 | fil3 | 30/30 | 2 | leptotene | circular , peripheral | 1 | 0 | yes |
| mnd1 | 18 | ant 2, fil 1 | 0/30 | 1 | zygotene | not visible | 1 | 76-99 | yes |
| mnd1 | 18 | fil2 | 0/30 | 1 | zygotene | not visible | 1 | 76-99 | yes |
| mnd1 | 18 | fil3 | 0/30 | 1 | zygotene | not visible | 1 | 76-99 | yes |
| mnd1 | 18 | ant 3 fil1 | 30/30 | 2 | zygotene | not visible | 1 | 2575 | yes |
| mnd1 | 18 | fil 2 | 30/30 | 2 | zygotene | not visible | 1 | 26-75 | yes |
| mnd1 | 18 | fil 3 | 30/30 | 2 | zygotene | not visible | 1 | 26-75 | yes |
| mnd1 | 18 | ant 4 fil 1 | 0/30 | 0 | eP | circular,peripheral | 1 | 100 | no |
| mnd1 | 18 | fil2 | 0/30 | 0 | eP | circular,peripheral | 1 | 100 | no |
| mnd1 | 18 | fil3 | 0/30 | 0 | eP | circular,peripheral | 1 | 100 | no |
| mnd1 | 18 | **bud 6**, ant1, fil1 | 30/30 | 2 | zygotene | not visible | 1 | 2575 | yes |
| mnd1 | 18 | fil 2 | 30/30 | 2 | zygotene | not visible | 1 | 26-75 | yes |

| | | | | | | | | | |
|---|---|---|---|---|---|---|---|---|---|
| mnd1 | 18 | fil3 | 30/30 | 2 | zygotene | not visible | 1 | 26-75 | yes |
| mnd1 | 18 | ant 2, fil 1 | 30/30 | 2 | leptotene | circular , peripheral | 1 | 0 | yes |
| mnd1 | 18 | fil2 | 30/30 | 2 | leptotene | circular , peripheral | 1 | 0 | yes |
| mnd1 | 18 | fil3 | 30/30 | 2 | leptotene | circular , peripheral | 1 | 0 | yes |
| mnd1 | 18 | ant 3 fil1 | 30/30 | 2 | leptotene | circular , peripheral | 1 | 0 | yes |
| mnd1 | 18 | fil 2 | 30/30 | 2 | leptotene | circular , peripheral | 1 | 0 | yes |
| mnd1 | 18 | fil 3 | 30/30 | 2 | leptotene | circular , peripheral | 1 | 0 | yes |
| mnd1 | 18 | ant 4 fil 1 | 30/30 | 2 | leptotene | circular , peripheral | 1 | 0 | yes |
| mnd1 | 18 | fil2 | 30/30 | 2 | leptotene | circular , peripheral | 1 | 0 | yes |
| mnd1 | 18 | fil3 | 30/30 | 2 | leptotene | circular , peripheral | 1 | 0 | yes |
| mnd1 | 18 | ant 5 fil 1 | 30/30 | 1 | leptotene | circular , pericentric | 1 | 0 | yes |
| mnd1 | 18 | fil2 | 30/30 | 1 | leptotene | circular , pericentric | 1 | 0 | yes |
| mnd1 | 18 | fil3 | 30/30 | 1 | leptotene | circular , pericentric | 1 | 0 | yes |
| mnd1 | 18 | **bud 7**, ant1, fil1 | 0/30 | 0 | m-l P | circular,peripheral | 2 | 100 | yes |
| mnd1 | 18 | fil 2 | 0/30 | 0 | m-l P | circular,peripheral | 2 | 100 | yes |
| mnd1 | 18 | fil3 | 0/30 | 0 | m-l P | circular,peripheral | 2 | 100 | yes |
| mnd1 | 18 | ant 2, fil 1 | 0/30 | 0 | m-l P | circular,peripheral | 3 | 100 | yes |
| mnd1 | 18 | fil2 | 0/30 | 0 | m-l P | circular,peripheral | 3 | 100 | yes |
| mnd1 | 18 | fil3 | 0/30 | 0 | m-l P | circular,peripheral | 3 | 100 | yes |
| mnd1 | 18 | ant 3 fil1 | 0/30 | 0 | m-l P | circular,peripheral | 3 | 100 | yes |
| mnd1 | 18 | fil 2 | 0/30 | 0 | m-l P | circular,peripheral | 3 | 100 | yes |
| mnd1 | 18 | fil 3 | 0/30 | 0 | m-l P | circular,peripheral | 3 | 100 | yes |
| mnd1 | 18 | **bud 8**, ant1, fil1 | 0/30 | 0 | tetrad | circular , centric | 3 | 100 | no |
| mnd1 | 18 | fil 2 | 0/30 | 0 | tetrad | circular , centric | 3 | 100 | no |
| mnd1 | 18 | fil3 | 0/30 | 0 | tetrad | circular , centric | 3 | 100 | no |
| mnd1 | 18 | **bud 9**, ant1, fil1 | 0/30 | 1 | leptotene | circular , centric | 1 | 0 | yes |
| mnd1 | 18 | fil 2 | 0/30 | 1 | leptotene | circular , centric | 1 | 0 | yes |
| mnd1 | 18 | fil3 | 0/30 | 1 | leptotene | circular , centric | 1 | 0 | yes |
| mnd1 | 18 | ant 2, fil 1 | 0/30 | 1 | leptotene | circular , centric | 1 | 0 | yes |

| | | | | | | | | | |
|---|---|---|---|---|---|---|---|---|---|
| mnd1 | 18 | fil2 | 0/30 | 1 | leptotene | circular , centric | 1 | 0 | yes |
| mnd1 | 18 | fil3 | 0/30 | 1 | leptotene | circular , centric | 1 | 0 | yes |
| mnd1 | 18 | ant 3 fil1 | 30/30 | 1 | zygotene | not visible | 1 | 1 to 25 | yes |
| mnd1 | 18 | fil 2 | 30/30 | 1 | zygotene | not visible | 1 | 1 to 25 | yes |
| mnd1 | 18 | fil 3 | 30/30 | 1 | zygotene | not visible | 1 | 1 to 25 | yes |
| mnd1 | 18 | ant 4 fil 1 | 30/30 | 2 | zygotene | not visible | 1 | 1 to 25 | yes |
| mnd1 | 18 | fil2 | 30/30 | 2 | zygotene | not visible | 1 | 1 to 25 | yes |
| mnd1 | 18 | fil3 | 30/30 | 2 | zygotene | not visible | 1 | 1 to 25 | yes |
| mnd1 | 18 | ant 5 fil 1 | 30/30 | 1 | zygotene | not visible | 1 | 1 to 25 | yes |
| mnd1 | 18 | fil2 | 30/30 | 1 | zygotene | not visible | 1 | 1 to 25 | yes |
| mnd1 | 18 | fil3 | 30/30 | 1 | zygotene | not visible | 1 | 1 to 25 | yes |
| mnd1 | 18 | **bud 10**, ant1, fil1 | 30/30 | 1 | leptotene | circular , centric | 1 | 0 | yes |
| mnd1 | 18 | fil 2 | 30/30 | 1 | leptotene | circular , centric | 1 | 0 | yes |
| mnd1 | 18 | fil3 | 30/30 | 1 | leptotene | circular , centric | 1 | 0 | yes |
| mnd1 | 18 | **bud 11**, ant1, fil1 | 0/30 | 1 | zygotene | not visible | 1 | 76-99 | yes |
| mnd1 | 18 | fil 2 | 0/30 | 1 | zygotene | not visible | 1 | 76-99 | yes |
| mnd1 | 18 | fil3 | 0/30 | 1 | zygotene | not visible | 1 | 76-99 | yes |
| mnd1 | 20 | **bud 1**, ant1, fil1 | 0/30 | 0 | m-l P | circular,peripheral | 2 | 100 | no |
| mnd1 | 20 | fil 2 | 0/30 | 0 | m-l P | circular,peripheral | 2 | 100 | no |
| mnd1 | 20 | ant 2, fil 1 | 0/30 | 0 | m-l P | circular,peripheral | 2 | 100 | yes |
| mnd1 | 20 | fil2 | 0/30 | 0 | m-l P | circular,peripheral | 2 | 100 | yes |
| mnd1 | 20 | fil3 | 0/30 | 0 | m-l P | circular,peripheral | 2 | 100 | yes |
| mnd1 | 20 | ant 3 fil1 | 0/30 | 0 | eP | circular,peripheral | 1 | 100 | no |
| mnd1 | 20 | fil 2 | 0/30 | 0 | eP | circular,peripheral | 1 | 100 | no |
| mnd1 | 20 | ant 4 fil 1 | 0/30 | 1 | zygotene | not visible | 1 | 76-99 | yes |
| mnd1 | 20 | fil2 | 0/30 | 1 | zygotene | not visible | 1 | 76-99 | yes |
| mnd1 | 20 | **bud 2**, ant1, fil1 | 20/30 | 1 | zygotene | not visible | 1 | 26-75 | yes |

| | | | | | | | | | |
|---|---|---|---|---|---|---|---|---|---|
| mnd1 | 20 | fil 2 | 25/30 | 1 | zygotene | not visible | 1 | 26-75 | yes |
| mnd1 | 20 | ant 2 fil1 | 25/30 | 2 | zygotene | not visible | 1 | 1 to 25 | yes |
| mnd1 | 20 | fil 2 | 25/30 | 2 | zygotene | not visible | 1 | 1 to 25 | yes |
| mnd1 | 20 | ant 3, fil 1 | 0/30 | 0 | m-l P | circular,peripheral | 2 | 100 | no |
| mnd1 | 20 | fil2 | 0/30 | 0 | m-l P | circular,peripheral | 2 | 100 | no |
| mnd1 | 20 | fil3 | 0/30 | 0 | m-l P | circular,peripheral | 2 | 100 | no |
| mnd1 | 20 | ant 4, fil 1 | 25/30 | 2 | zygotene | not visible | 1 | 1 to 25 | yes |
| mnd1 | 20 | fil2 | 20/30 | 2 | zygotene | not visible | 1 | 1 to 25 | yes |
| mnd1 | 20 | fil3 | 25/30 | 2 | zygotene | not visible | 1 | 1 to 25 | yes |
| mnd1 | 20 | **bud 3**, ant1, fil1 | 0/30 | 1 | eP | circular,peripheral | 1 | 100 | yes |
| mnd1 | 20 | fil 2 | 0/30 | 1 | eP | circular,peripheral | 1 | 100 | yes |
| mnd1 | 20 | fil3 | 0/30 | 1 | eP | circular,peripheral | 1 | 100 | yes |
| mnd1 | 20 | ant 2, fil 1 | 0/30 | 0 | m-l P | circular,peripheral | 2 | 100 | yes |
| mnd1 | 20 | fil2 | 0/30 | 0 | m-l P | circular,peripheral | 2 | 100 | yes |
| mnd1 | 20 | fil3 | 0/30 | 0 | m-l P | circular,peripheral | 2 | 100 | yes |
| mnd1 | 20 | **bud 4**, ant1, fil1 | 0/30 | 0 | eP | circular,peripheral | 1 | 100 | no |
| mnd1 | 20 | fil 2 | 0/30 | 0 | eP | circular,peripheral | 1 | 100 | no |
| mnd1 | 20 | **bud 5**, ant1, fil1 | 0/30 | 0 | metaphase I | not visible | N/A | 100 | no |
| mnd1 | 20 | fil 2 | 0/30 | 0 | metaphase I | not visible | N/A | 100 | no |
| mnd1 | 20 | fil3 | 0/30 | 0 | tetrad | circular , centric | 3 | 100 | no |
| mnd1 | 20 | **bud 6**, ant1, fil1 | 30/30 | 1 | leptotene | circular , pericentric | 1 | 0 | yes |
| mnd1 | 20 | fil 2 | 30/30 | 1 | leptotene | circular , pericentric | 1 | 0 | yes |
| mnd1 | 20 | fil 3 | 30/30 | 1 | leptotene | circular , pericentric | 1 | 0 | yes |
| mnd1 | 20 | ant 2, fil 1 | 30/30 | 2 | zygotene | not visible | 1 | 1 to 25 | yes |
| mnd1 | 20 | fil2 | 30/30 | 2 | zygotene | not visible | 1 | 1 to 25 | yes |
| mnd1 | 20 | fil3 | 30/30 | 2 | zygotene | not visible | 1 | 1 to 25 | yes |
| mnd1 | 20 | ant 3 fil1 | 30/30 | 2 | zygotene | not visible | 1 | 1 to 25 | yes |
| mnd1 | 20 | fil 2 | 30/30 | 2 | zygotene | not visible | 1 | 1 to 25 | yes |
| mnd1 | 20 | fil 3 | 30/30 | 2 | zygotene | not visible | 1 | 1 to 25 | yes |

| | | | | | | | | | |
|---|---|---|---|---|---|---|---|---|---|
| mnd1 | 22 | **bud 1**, ant1, fil1 | 0/30 | 0 | G2 | circular , centric | 1 | 0 | no |
| mnd1 | 22 | fil 2 | 0/30 | 0 | G2 | circular , centric | 1 | 0 | no |
| mnd1 | 22 | fil 3 | 0/30 | 0 | G2 | circular , centric | 1 | 0 | no |
| mnd1 | 22 | ant 2, fil 1 | 0/30 | 0 | G2 | circular , centric | 1 | 0 | no |
| mnd1 | 22 | fil2 | 0/30 | 0 | G2 | circular , centric | 1 | 0 | no |
| mnd1 | 22 | ant 3 fil1 | 0/30 | 0 | G2 | circular , centric | 1 | 0 | no |
| mnd1 | 22 | fil 2 | 0/30 | 0 | G2 | circular , centric | 1 | 0 | no |
| mnd1 | 22 | fil 3 | 0/30 | 0 | G2 | circular , centric | 1 | 0 | no |
| mnd1 | 22 | **bud 2**, ant1, fil1 | 0/30 | 0 | tetrad | circular , centric | 3 | 100 | no |
| mnd1 | 22 | fil 2 | 0/30 | 0 | tetrad | circular , centric | 3 | 100 | no |
| mnd1 | 22 | fil 3 | 0/30 | 0 | tetrad | circular , centric | 3 | 100 | no |
| mnd1 | 22 | **bud 3**, ant1, fil1 | 30/30 | 2 | leptotene | circular , pericentric | 1 | 0 | yes |
| mnd1 | 22 | fil 2 | 30/30 | 2 | leptotene | circular , pericentric | 1 | 0 | yes |
| mnd1 | 22 | fil 3 | 30/30 | 2 | leptotene | circular , pericentric | 1 | 0 | yes |
| mnd1 | 22 | ant 2, fil 1 | 30/30 | 1 | leptotene | circular , pericentric | 1 | 0 | yes |
| mnd1 | 22 | fil2 | 30/30 | 1 | leptotene | circular , pericentric | 1 | 0 | yes |
| mnd1 | 22 | fil3 | 30/30 | 1 | leptotene | circular , pericentric | 1 | 0 | yes |
| **mnd1** | **22** | **bud 4, ant1, fil1** | **30/30** | **0** | **eP** | **circular,peripheral** | **1** | **100** | **yes** |
| **mnd1** | **22** | **fil 2** | **30/30** | **0** | **eP** | **circular,peripheral** | **1** | **100** | **yes** |
| **mnd1** | **22** | **fil 3** | **30/30** | **0** | **eP** | **circular,peripheral** | **1** | **100** | **yes** |
| mnd1 | 22 | ant 2, fil 1 | 7/30 | 1 | zygotene | not visible | 1 | 76-99 | yes |
| mnd1 | 22 | fil2 | 10/30 | 1 | zygotene | not visible | 1 | 76-99 | yes |
| mnd1 | 22 | fil3 | 5/30 | 1 | zygotene | not visible | 1 | 76-99 | yes |
| mnd1 | 22 | **bud 5**, ant1, fil1 | 0/30 | 0 | G2 | circular , centric | 1 | 0 | no |
| mnd1 | 22 | fil 2 | 0/30 | 0 | G2 | circular , centric | 1 | 0 | no |
| mnd1 | 22 | fil 3 | 0/30 | 0 | G2 | circular , centric | 1 | 0 | no |
| mnd1 | 22 | ant 2, fil 1 | 0/30 | 0 | G2 | circular , centric | 1 | 0 | no |
| mnd1 | 22 | fil2 | 0/30 | 0 | G2 | circular , centric | 1 | 0 | no |

| | | | | | | | | | |
|---|---|---|---|---|---|---|---|---|---|
| mnd1 | 22 | fil3 | 0/30 | 0 | G2 | circular , centric | 1 | 0 | no |
| mnd1 | 22 | **bud 6**, ant1, fil1 | 0/30 | 1 | leptotene | circular , centric | 1 | 0 | yes |
| mnd1 | 22 | fil 2 | 0/30 | 1 | leptotene | circular , centric | 1 | 0 | yes |
| mnd1 | 22 | fil 3 | 0/30 | 1 | leptotene | circular , centric | 1 | 0 | yes |
| mnd1 | 22 | **bud 7**, ant1, fil1 | 30/30 | 0 | eP | circular,peripheral | 1 | 100 | yes |
| mnd1 | 22 | fil 2 | 30/30 | 0 | eP | circular,peripheral | 1 | 100 | yes |
| mnd1 | 22 | fil 3 | 30/30 | 0 | eP | circular,peripheral | 1 | 100 | yes |
| mnd1 | 22 | ant 2, fil 1 | 30/30 | 0 | eP | circular,peripheral | 1 | 100 | yes |
| mnd1 | 22 | fil 2 | 30/30 | 0 | eP | circular,peripheral | 1 | 100 | yes |
| mnd1 | 22 | fil 3 | 30/30 | 0 | eP | circular,peripheral | 1 | 100 | yes |
| mnd1 | 22 | fil 4 | 30/30 | 0 | eP | circular,peripheral | 1 | 100 | yes |
| mnd1 | 22 | ant 3 fil1 | 30/30 | 0 | eP | circular,peripheral | 1 | 100 | yes |
| mnd1 | 22 | fil 2 | 30/30 | 0 | eP | circular,peripheral | 1 | 100 | yes |
| mnd1 | 22 | fil 3 | 30/30 | 0 | eP | circular,peripheral | 1 | 100 | yes |
| mnd1 | 22 | fil 4 | 30/30 | 0 | eP | circular,peripheral | 1 | 100 | yes |
| mnd1 | 22 | ant 4 fil 1 | 30/30 | 0 | eP | circular,peripheral | 1 | 100 | yes |
| mnd1 | 22 | fil 2 | 30/30 | 0 | eP | circular,peripheral | 1 | 100 | yes |
| mnd1 | 22 | fil 3 | 30/30 | 0 | eP | circular,peripheral | 1 | 100 | yes |
| mnd1 | 22 | fil 4 | 30/30 | 0 | eP | circular,peripheral | 1 | 100 | yes |
| mnd1 | 22 | **bud 8**, ant1, fil1 | 0/30 | 2 | leptotene | circular , pericentric | 1 | 0 | yes |
| mnd1 | 22 | fil 2 | 0/30 | 2 | leptotene | circular , pericentric | 1 | 0 | yes |
| mnd1 | 22 | fil3 | 0/30 | 2 | leptotene | circular , pericentric | 1 | 0 | yes |
| mnd1 | 22 | ant 2, fil 1 | 0/30 | 2 | leptotene | circular , pericentric | 1 | 0 | yes |
| mnd1 | 22 | fil2 | 0/30 | 2 | leptotene | circular , pericentric | 1 | 0 | yes |
| mnd1 | 22 | fil3 | 0/30 | 2 | leptotene | circular , pericentric | 1 | 0 | yes |
| mnd1 | 22 | ant 3 fil1 | 0/30 | 1 | leptotene | circular , pericentric | 1 | 0 | yes |
| mnd1 | 22 | fil 2 | 5/30 | 1 | leptotene | circular , pericentric | 1 | 0 | yes |
| mnd1 | 22 | fil 3 | 10/30 | 1 | leptotene | circular , pericentric | 1 | 0 | yes |

| | | | | | | | | | |
|---|---|---|---|---|---|---|---|---|---|
| mnd1 | 24 | **bud 1**, ant1, fil1 | 5/30 | 1 | leptotene | circular , pericentric | 1 | 0 | yes |
| mnd1 | 24 | fil 2 | 10/30 | 1 | leptotene | circular , pericentric | 1 | 0 | yes |
| mnd1 | 24 | fil3 | 5/30 | 1 | leptotene | circular , pericentric | 1 | 0 | yes |
| mnd1 | 24 | **bud 2**, ant1, fil1 | 0/30 | 1 | leptotene | circular , pericentric | 1 | 0 | yes |
| mnd1 | 24 | fil 2 | 0/30 | 1 | leptotene | circular , pericentric | 1 | 0 | yes |
| mnd1 | 24 | fil3 | 0/30 | 1 | leptotene | circular , pericentric | 1 | 0 | yes |
| mnd1 | 24 | ant 2, fil 1 | 0/30 | 1 | leptotene | circular , pericentric | 1 | 0 | yes |
| mnd1 | 24 | fil2 | 0/30 | 1 | leptotene | circular , pericentric | 1 | 0 | yes |
| mnd1 | 24 | fil3 | 0/30 | 1 | leptotene | circular , pericentric | 1 | 0 | yes |
| mnd1 | 24 | ant 3 fil1 | 0/30 | 1 | leptotene | circular , pericentric | 1 | 0 | yes |
| mnd1 | 24 | fil 2 | 0/30 | 1 | leptotene | circular , pericentric | 1 | 0 | yes |
| mnd1 | 24 | fil 3 | 0/30 | 1 | leptotene | circular , pericentric | 1 | 0 | yes |
| mnd1 | 26 | **bud 1** ant 1, fil 1 | 30/30 | 1 | zygotene | not visible | 1 | 76-99 | yes |
| mnd1 | 26 | fil 2 | 30/30 | 1 | zygotene | not visible | 1 | 76-99 | yes |
| mnd1 | 26 | fil 3 | 30/30 | 1 | zygotene | not visible | 1 | 76-99 | yes |
| mnd1 | 26 | fil 4 | 30/30 | 1 | zygotene | not visible | 1 | 76-99 | yes |
| mnd1 | 26 | ant 2 fil1 | 30/30 | 1 | zygotene | not visible | 1 | 76-99 | yes |
| mnd1 | 26 | fil 2 | 30/30 | 1 | zygotene | not visible | 1 | 76-99 | yes |
| mnd1 | 26 | fil 3 | 30/30 | 1 | zygotene | not visible | 1 | 76-99 | yes |
| mnd1 | 26 | fil 4 | 30/30 | 1 | zygotene | not visible | 1 | 76-99 | yes |
| mnd1 | 28 | **bud 1** ant 1, fil 1 | 30/30 | 2 | zygotene | not visible | 1 | 1 to 25 | yes |
| mnd1 | 28 | fil 2 | 30/30 | 2 | zygotene | not visible | 1 | 1 to 25 | yes |
| mnd1 | 28 | fil 3 | 30/30 | 2 | zygotene | not visible | 1 | 1 to 25 | yes |
| mnd1 | 28 | fil 4 | 30/30 | 2 | zygotene | not visible | 1 | 1 to 25 | yes |
| mnd1 | 28 | ant 2 fil1 | 30/30 | 2 | zygotene | not visible | 1 | 1 to 25 | yes |
| mnd1 | 28 | fil 2 | 30/30 | 2 | zygotene | not visible | 1 | 1 to 25 | yes |
| mnd1 | 28 | fil 3 | 30/30 | 2 | zygotene | not visible | 1 | 1 to 25 | yes |

| | | | | | | | | | |
|---|---|---|---|---|---|---|---|---|---|
| mnd1 | 28 | fil 4 | 30/30 | 2 | zygotene | not visible | 1 | 1 to 25 | yes |
| | | **bud 2** ant 1, fil | | | | | | | |
| mnd1 | 28 | 1 | 30/30 | 0 | m-l P | circular,peripheral | 2 | 100 | yes |
| mnd1 | 28 | fil 2 | 30/30 | 0 | m-l P | circular,peripheral | 2 | 100 | yes |
| mnd1 | 28 | fil 3 | 30/30 | 0 | m-l P | circular,peripheral | 2 | 100 | yes |
| mnd1 | 28 | fil 4 | 30/30 | 0 | m-l P | circular,peripheral | 2 | 100 | yes |
| | | **bud 3** ant 1, fil | | | | | | | |
| mnd1 | 28 | 1 | 0/30 | 0 | tetrad | circular , centric | 3 | 100 | no |
| mnd1 | 28 | fil 2 | 0/30 | 0 | tetrad | circular , centric | 3 | 100 | no |
| mnd1 | 28 | fil 3 | 0/30 | 0 | tetrad | circular , centric | 3 | 100 | no |
| mnd1 | 28 | fil 4 | 0/30 | 0 | tetrad | circular , centric | 3 | 100 | no |
| | | **bud 4** ant 1, fil | | | | | | | |
| mnd1 | 28 | 1 | 30/30 | 1 | leptotene | circular , pericentric | 1 | 0 | yes |
| mnd1 | 28 | fil 2 | 30/30 | 1 | leptotene | circular , pericentric | 1 | 0 | yes |
| mnd1 | 28 | fil 3 | 30/30 | 1 | leptotene | circular , pericentric | 1 | 0 | yes |
| mnd1 | 28 | fil 4 | 30/30 | 1 | leptotene | circular , pericentric | 1 | 0 | yes |
| | | **bud 5** ant 1, fil | | | | | | | |
| mnd1 | 28 | 1 | 30/30 | 0 | m-l P | circular,peripheral | 2 | 100 | yes |
| mnd1 | 28 | fil 2 | 30/30 | 0 | m-l P | circular,peripheral | 2 | 100 | yes |
| mnd1 | 28 | fil 3 | 30/30 | 0 | m-l P | circular,peripheral | 2 | 100 | yes |
| mnd1 | 28 | fil 4 | 30/30 | 0 | m-l P | circular,peripheral | 2 | 100 | yes |
| | | **bud 6** ant 1, fil | | | | | | | |
| mnd1 | 28 | 1 | 0/30 | 0 | anaphase I | not visible | N/A | 100 | no |
| mnd1 | 28 | fil 2 | 0/30 | 0 | anaphase I | not visible | N/A | 100 | no |
| mnd1 | 28 | fil 3 | 0/30 | 0 | anaphase I | not visible | N/A | 100 | no |
| mnd1 | 28 | fil 4 | 0/30 | 0 | anaphase I | not visible | N/A | 100 | no |
| | | **bud 1** ant 1, fil | | | | | | | |
| mnd1 | 30 | 1 | 0/30 | 0 | m-l P | circular,peripheral | 3 | 100 | yes |
| mnd1 | 30 | fil 2 | 0/30 | 0 | m-l P | circular,peripheral | 3 | 100 | yes |
| mnd1 | 30 | fil 3 | 0/30 | 0 | m-l P | circular,peripheral | 3 | 100 | yes |
| mnd1 | 30 | fil 4 | 0/30 | 0 | m-l P | circular,peripheral | 3 | 100 | yes |

| | | | | | | | | | |
|---|---|---|---|---|---|---|---|---|---|
| mnd1 | 30 | **bud 2** ant 1, fil 1 | 0/30 | 0 | G2 | circular , centric | 1 | 0 | no |
| mnd1 | 30 | fil 2 | 0/30 | 0 | G2 | circular , centric | 1 | 0 | no |
| mnd1 | 30 | fil 3 | 0/30 | 0 | G2 | circular , centric | 1 | 0 | no |
| mnd1 | 30 | fil 4 | 0/30 | 0 | G2 | circular , centric | 1 | 0 | no |
| mnd1 | 30 | ant 2, fil 1 | 20/30 | 1 | leptotene | circular , centric | 1 | 0 | yes |
| mnd1 | 30 | fil 2 | 25/30 | 1 | leptotene | circular , centric | 1 | 0 | yes |
| mnd1 | 30 | fil 3 | 25/30 | 1 | leptotene | circular , centric | 1 | 0 | yes |
| mnd1 | 30 | fil 4 | 10/30 | 1 | leptotene | circular , centric | 1 | 0 | yes |
| mnd1 | 30 | ant 3 fil1 | 10/30 | 1 | leptotene | circular , centric | 1 | 0 | yes |
| mnd1 | 30 | fil 2 | 5/30 | 1 | leptotene | circular , centric | 1 | 0 | yes |
| mnd1 | 30 | fil 3 | 10/30 | 1 | leptotene | circular , centric | 1 | 0 | yes |
| mnd1 | 30 | ant 4 fil 1 | 10/30 | 1 | leptotene | circular , centric | 1 | 0 | yes |
| mnd1 | 30 | fil 2 | 10/30 | 1 | leptotene | circular , centric | 1 | 0 | yes |
| mnd1 | 30 | fil 3 | 5/30 | 1 | leptotene | circular , centric | 1 | 0 | yes |
| mnd1 | 30 | **buds 3** ant 1, fil 1 | 30/30 | 0 | m-l P | circular,peripheral | 3 | 100 | yes |
| mnd1 | 30 | fil 2 | 30/30 | 0 | m-l P | circular,peripheral | 3 | 100 | yes |
| mnd1 | 30 | fil 3 | 30/30 | 0 | m-l P | circular,peripheral | 3 | 100 | yes |
| mnd1 | 30 | fil 4 | 30/30 | 0 | m-l P | circular,peripheral | 3 | 100 | yes |
| mnd1 | 30 | ant 2, fil 1 | 30/30 | 0 | m-l P | circular,peripheral | 3 | 100 | yes |
| mnd1 | 30 | fil 2 | 30/30 | 0 | m-l P | circular,peripheral | 3 | 100 | yes |
| mnd1 | 30 | fil 3 | 30/30 | 0 | m-l P | circular,peripheral | 3 | 100 | yes |
| mnd1 | 30 | fil 4 | 30/30 | 0 | m-l P | circular,peripheral | 3 | 100 | yes |
| mnd1 | 32 | **bud 1** ant 1, fil 1 | 5/30 | 1 | leptotene | circular , centric | 1 | 0 | yes |
| mnd1 | 32 | fil 2 | 0/30 | 1 | leptotene | circular , centric | 1 | 0 | yes |
| mnd1 | 32 | fil 3 | 5/30 | 1 | leptotene | circular , centric | 1 | 0 | yes |
| mnd1 | 32 | **bud 2**, ant1, fil1 | 30/30 | 1 | eP | circular,peripheral | 1 | 100 | yes |
| mnd1 | 32 | fil 2 | 30/30 | 1 | eP | circular,peripheral | 1 | 100 | yes |

| | | | | | | | | | |
|---|---|---|---|---|---|---|---|---|---|
| mnd1 | 32 | fil3 | 30/30 | 1 | eP | circular,peripheral | 1 | 100 | yes |
| mnd1 | 32 | fil 4 | 30/30 | 1 | eP | circular,peripheral | 1 | 100 | yes |
| mnd1 | 32 | ant2, fil1 | 30/30 | 1 | eP | circular,peripheral | 1 | 100 | yes |
| mnd1 | 32 | fil 2 | 30/30 | 1 | eP | circular,peripheral | 1 | 100 | yes |
| mnd1 | 32 | fil3 | 30/30 | 1 | eP | circular,peripheral | 1 | 100 | yes |
| mnd1 | 32 | **bud 3** ant 1, fil 1 | 0/30 | 0 | m-l P | circular,peripheral | 3 | 100 | no |
| mnd1 | 32 | fil 2 | 0/30 | 0 | m-l P | circular,peripheral | 3 | 100 | no |
| mnd1 | 32 | fil 3 | 0/30 | 0 | m-l P | circular,peripheral | 3 | 100 | no |
| mnd1 | 32 | **bud 4**, ant1, fil1 | 0/30 | 1 | leptotene | circular , centric | 1 | 0 | yes |
| mnd1 | 32 | fil 2 | 0/30 | 1 | leptotene | circular , centric | 1 | 0 | yes |
| mnd1 | 32 | fil3 | 0/30 | 1 | leptotene | circular , centric | 1 | 0 | yes |
| mnd1 | 32 | ant2, fil1 | 0/30 | 1 | leptotene | circular , centric | 1 | 0 | yes |
| mnd1 | 32 | fil 2 | 0/30 | 1 | leptotene | circular , centric | 1 | 0 | yes |
| mnd1 | 32 | fil3 | 0/30 | 1 | leptotene | circular , centric | 1 | 0 | yes |
| mnd1 | 32 | **bud 5**, ant1, fil1 | 0/30 | 0 | m-l P | circular,peripheral | 3 | 100 | no |
| mnd1 | 32 | fil 2 | 0/30 | 0 | m-l P | circular,peripheral | 3 | 100 | no |
| mnd1 | 32 | fil3 | 0/30 | 0 | m-l P | circular,peripheral | 3 | 100 | no |
| mnd1 | 32 | ant 2, fil 1 | 0/30 | 0 | m-l P | circular,peripheral | 3 | 100 | no |
| mnd1 | 32 | fil2 | 0/30 | 0 | m-l P | circular,peripheral | 3 | 100 | no |
| mnd1 | 32 | fil3 | 0/30 | 0 | m-l P | circular,peripheral | 3 | 100 | no |
| mnd1 | 32 | ant 3 fil1 | 30/30 | 1 | eP | circular,peripheral | 1 | 100 | yes |
| mnd1 | 32 | fil 2 | 30/30 | 1 | eP | circular,peripheral | 1 | 100 | yes |
| mnd1 | 32 | fil 3 | 30/30 | 1 | eP | circular,peripheral | 1 | 100 | yes |
