## Supplemental Table S2 for "The Arabidopsis *HOP2* Gene Has a Role in Preventing illegitimate Exchanges between Nonhomologous Chromosomes"

**Supplemental Table S2.** Primers for SSLP genotyping

| Chromosome/position | Primer name | Sequence |
|---|---|---|
| Chr1: 9.62Mb | CIW12 FOR | AGGTTTTATTGCTTTTCACA |
| | CIW12 BACK | CTTTCAAAAGCACATCACA |
| Chr2: 6.4Mb | CIW3 FOR | GAAACTCAATGAAATCCACTT |
| | CIW3 BACK | TGAACTTGTTGTGAGCTTTGA |
| Chr3: 9.8Mb | CIW11 FOR | CCCCGAGTTGAGGTATT |
| | CIW11 BACK | TGAACTTGTTGTGAGCTTTGA |
| Chr4: 16.4Mb | NGA1139 FOR | TAGCCGGATGAGTTGGTACC |
| | NGA1139 BACK | TTTTTCCTTGTGTTGCATTCC |
| Chr5: 14Mb | ATPHYC FOR | CTCAGAGAATTCCCAGAAAAATCT |
| | ATPHYC BACK | AAACTCGAGAGTTTTGTCTAGATC |
