## Supplemental Figures S1 and S2 and S3 for "The Arabidopsis *HOP2* Gene Has a Role in Preventing illegitimate Exchanges between Nonhomologous Chromosomes"

**Figure S1. Timing of γH2AX foci in genotypes Columbia (Col) and *mnd1* (Columbia background), Landsberg *erecta* (L*er*), *hop2-1*(L*er* background) from all samples collected.**

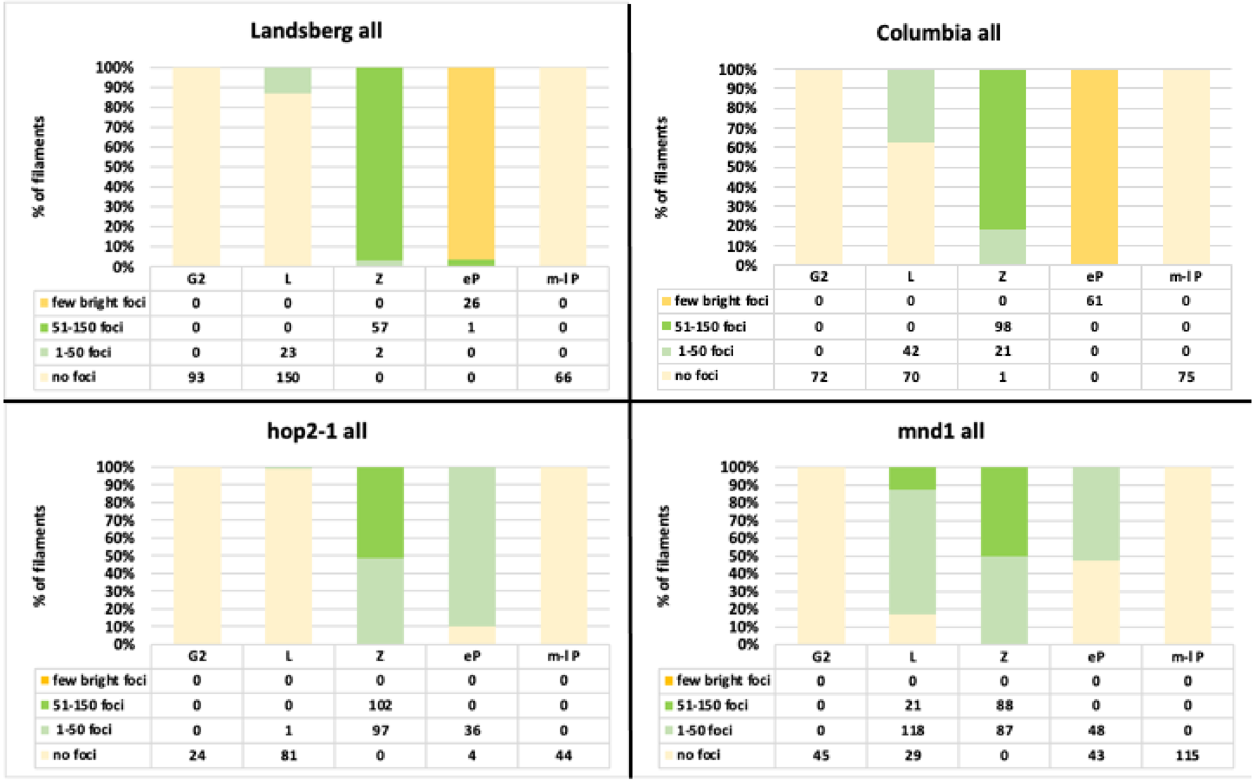


Figure S1. Timing of γH2AX foci in genotypes Columbia (Col) and *mnd1* (Columbia background), Landsberg *erecta* (L*er*), *hop2-1*(L*er background)*. All filaments of meiotic cells for the relevant meiotic stages were included in this analysis regardless of whether or not they had incorporated EdU. Each filament was classified as having 'no foci', '1- 50 small foci', '51-150 small foci', or 'a few brighter larger foci'. The number of filaments within each category are given in tabular form below each genotype's percentage of nuclei in each category. Stages assessed were G2 preceding meiosis, leptotene (L), zygotene (Z), early pachytene (eP) and mid-to late pachytene (m-lP). The mutant *hop2-1* had significantly fewer foci than L*er* at both leptotene and zygotene and the differences were statistically significant ($p<0.002$ and $p<0.00001$ respectively). In contrast the *mnd1* mutant had significantly more foci visible than Col at leptotene ($p<0.00001$), but significantly fewer than Col by zygotene ($p<0.0001$).

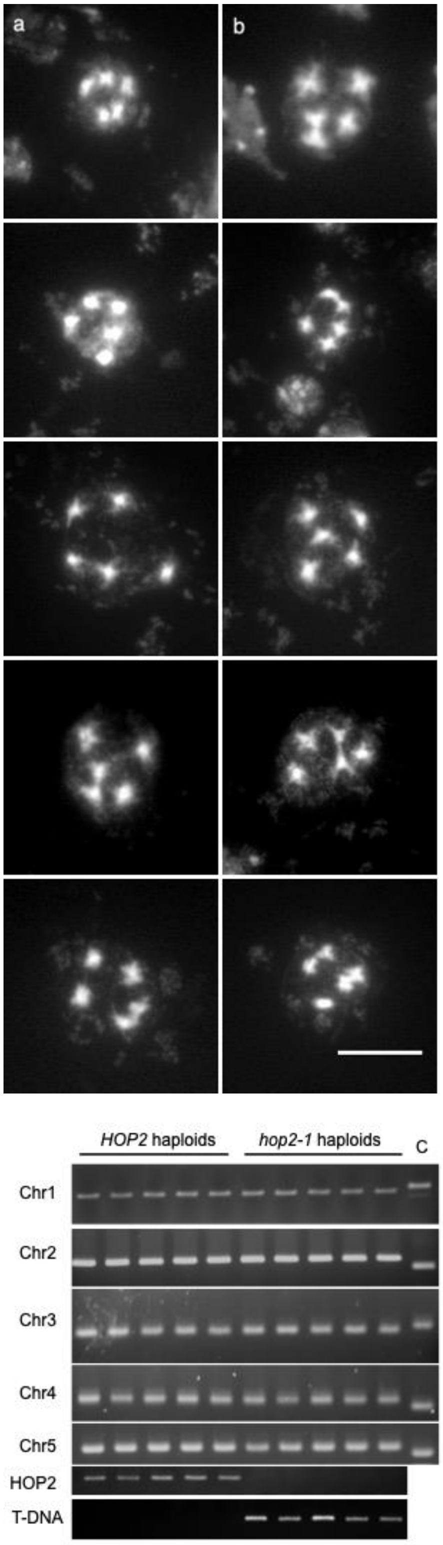


**Figure S2: Haploid plant genotyping.**
The five images of Fig.S2a (left side) are from plants genotyped as having the wild type *HOP2* allele, those in S2b (right side) are from haploids with the *hop2-1* allele. In Fig.S2c the putative haploid plants were subjected to SSLP analysis to ascertain parentage, and in all cases, gave rise to only L*er*-specific banding patterns. The first five lanes represent the five *HOP2* wildtype haploids; the five *hop2-1* mutant haploids are in lanes 6-10. The lane designated 'C' is a reaction with a Columbia template. The markers for each of the chromosomes are as follows 1, ciw12; 2, ciw3; 3, ciw11; 4, nga1139; 5, AthPHYC. The row labelled HOP2 represents reactions with primers designed to give a signal with wildtype DNA, while the row labelled T-DNA represents reactions in which a wildtype primer was used with a T-DNA primer to screen for the presence of a *hop2-1* T-DNA insertion. Primer sequences are given in the Experimental Procedures section and in Supplemental Table 2.

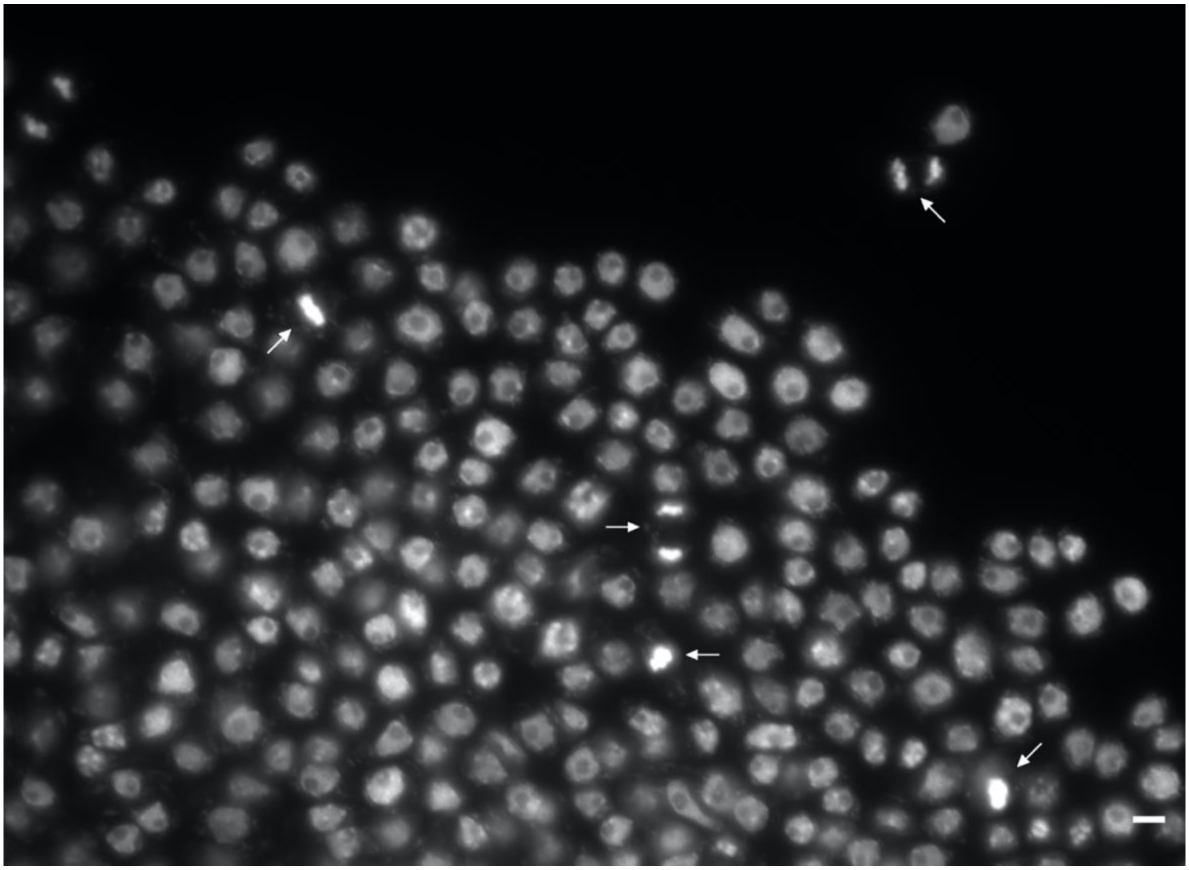

Figure S3. Petal cells from a segment of one petal of a Ler control plant. Arrows point to mitotic figures. Bar =10 μm
